# When does temporal resolution matter? Including detection covariates in discrete- versus continuous-time occupancy and N-mixture models

**DOI:** 10.1101/2025.08.31.673411

**Authors:** Léa Pautrel, Sylvain Moulherat, Benoit Charrasse, Guillaume Debat, Lucie Gendron, Kenneth Kellner, Marie-Pierre Etienne, Olivier Gimenez

**Author notes:** Co-senior authors (Marie-Pierre Etienne, Olivier Gimenez).

## Abstract

Camera traps and other sensors allow continuous-time biodiversity observation, raising new questions and opportunities for modelling detection in hierarchical models such as occupancy (for species presence) and N-mixture models (for abundance). We focused on a rarely considered aspect: how the temporal treatment of detection covariates affects inference. Through simulations and a five-month case study on an research center, we examined the effects of covariate temporal resolution, discretisation scale in discrete-time (DT) models, and interpolation methods in continuous-time (CT) models. While occupancy and abundance estimates were largely unaffected by these choices, detection estimates were more sensitive to them. DT models with fine temporal discretisation closely matched CT models. Simulations showed that when detection covariates had no effect on detectability, the considered modelling choices had little impact. But when covariates did influence detection, bias and error increased if their temporal variation was not accurately retained. The case study revealed more complex patterns, highlighting the consequences of temporally simplifying both observations and detection covariates. Overall, our results suggest that when detectability is of ecological interest, exploring a range of temporal treatments of detection covariates, from fine-scale to coarser resolutions, can reveal complementary insights into scale-dependent patterns in detection.

## 1 Introduction

Technological advancements have significantly influenced methodologies for addressing ecological challenges, particularly in the acquisition of fauna observation data (Besson et al., 2022; Lahoz-Monfort & Magrath, 2021). Sensors, such as camera traps (Burton et al., 2015) and acoustic devices (Biffi et al., 2024), enable monitoring a wide range of species across diverse habitats. For most species, sensor-based wildlife surveys do not allow individual identification (Tuia et al., 2022). In such cases, models suited to non-individualised animals (often referred to as unmarked data) are used. These typically estimate either species presence (Guillera-Arroita, 2017) or abundance (Gilbert et al., 2021) in relation to environmental factors, offering practical insights that inform conservation efforts. The area of occupancy describes the spatial distribution of a species and is one of the criteria used by the IUCN to establish the Red List of Ecosystems (Rodríguez et al., 2015). Estimating absolute abundance is crucial for developing broad-scale conservation strategies (Callaghan et al., 2024), as it can detect population changes even when the area of occupancy remains stable (Dennis et al., 2019). While not as informative as absolute abundance at broader scales, spatial relative abundance is especially valuable at local levels to prioritise conservation or mitigation actions (*e.g.* (Boussarie et al., 2023; Moulherat et al., 2024)).

In this work, we focus on hierarchical models that use non-individually identified observations, with two sub-models: an ecological state process and a detection process. Specifically, we consider two widely used categories of site-based, single-species, static models: occupancy (MacKenzie et al., 2002) and N-mixture (Royle, 2004) models. The ecological process represents either the presence/absence (occupancy) or the abundance of a species at a site. We restrict ourselves to static models, meaning we assume the ecological state remains constant over the study period. Occupancy and abundance are typically estimated in relation to spatial environmental covariates, like the habitat type.

These hierarchical models also explicitly account for imperfect detection, since for the vast majority of observation methods, not all individuals present are observed during sampling (Guillera-Arroita & Lahoz-Monfort, 2012; Mackenzie, 2005). The detection process in these hierarchical models is associated with the format of the observation data. Traditionally, detection is modelled in discrete-time (DT). Yet, with the recent increase in continuous monitoring via sensors, continuous-time (CT) detection processes have gained attention (Pautrel et al., 2024). Like the ecological state process, detection can also depend on environmental covariates that vary over time or space-time, such as weather conditions or human disturbance. Continuously collected observations require model users to make decisions during data preparation and model specification, and these choices differ depending on whether the modeller uses a DT or CT framework. In this work, we investigate how such decisions, which affect the detection process of these hierarchical models, influence inference in occupancy and N-mixture models. These decisions include how to incorporate detection covariates, commonly used in DT models, but still very rare in the few existing CT model applications, despite being formally described in previous studies (Guillera-Arroita et al., 2011; Kellner, Parsons, et al., 2022).

CT models offer several theoretical advantages over DT models for analysing continuously collected data (Guillera-Arroita et al., 2011; Pautrel et al., 2024; Rushing, 2023). These models retain all available information and enhance objectivity and reproducibility by eliminating the arbitrary discretisation step, although they still require decisions about the temporal resolution of covariates. DT occupancy and N-mixture models, developed for fieldwork sampling occasions separated in time, assume independence between sampling occasions (Bailey et al., 2014). This assumption is sensible in that context, but less relevant for consecutive discretised sessions. These theoretical advantages may encourage modellers to adopt CT models for analysing sensor-collected observation data.

However, CT models are not necessarily the panacea in all situations. Previous work on occupancy models without covariates found that DT and CT occupancy models provided equivalently accurate estimates of occupancy probability in most cases (Pautrel et al., 2024). This work did not consider covariates, which can significantly impact the detection process. Time-varying data such as weather conditions or human disturbance can be included as detection covariates, influencing species detectability. For instance, a lizard might be inactive during rain, rendering it undetectable.

Accounting for covariates involves different methodological considerations in DT and CT models. Covariates are often collected at regular intervals, such as hourly temperature measurements. In DT models, one must decide how to aggregate these covariates within the chosen discretisation intervals. In CT models, detection times may not align with covariate measurements. Therefore, it is necessary to interpolate covariate values to estimate their values at the times of detection and between detections. This is crucial because CT models relies on the integral value of covariates between each pair of detections to capture the full information, not just the covariate value at the time of detection. Numerous interpolation techniques are available. Simple methods include assuming constant covariate values until the next measurement or linear interpolation between successive measurements. More sophisticated approaches can also be used, such as constructing specific linear or non-linear, parametric or non-parametric regression models on covariates. For example, Gaussian process regression, also called Kriging in geostatistics, is particularly used for interpolation.

Building on previous comparisons that showed few differences between DT and CT models in estimating occupancy without covariates (Pautrel et al., 2024), we investigate in which cases CT models offer advantages when detection covariates are included and in which cases they do not, or even potentially degrade the quality of the results. Additionally, given the variety of possible interpolation methods, we aim to investigate how the interpolation method affects model outcomes for CT models. Our main objective is to provide recommendations for choosing hierarchical ecological models when analysing continuously collected fauna observations. Our assumptions are that CT models can be beneficial under the following conditions:

- If detectability is strongly influenced by covariates, CT models could provide better estimates and insights into species behaviour, such as reactions to human disturbance.
- The interpolation must be accurate. Inaccurate interpolation, due to sparse covariate data, could introduce errors or bias, negating potential improvements.
- CT models are most useful when the temporal scale of the covariate’s effect aligns with the scale at which the covariate is measured. For example, human presence might have a short-term impact (an animal hides for a few minutes, until the human leaves) or a longer-term one (the animal avoids the area for hours, days, or weeks). To detect these effects, the covariate must be measured at a matching temporal resolution. If human presence is recorded every minute, short-term effects can be detected. But if it is only measured or supplied to the model in an aggregated form – such as the sum or mean of passages per hour, day, or week – this fine-scale temporal variability is lost, and the model cannot distinguish between short- and long-term effects. This measurement scale depends both on methodological choices (*e.g.* aggregation) and on data availability (*e.g.* if only one measure per day is available, short-term variation cannot be captured). If the covariate does not have a short-term effect, or if it does not vary at short timescales (*e.g.* seasonal sunlight duration), the benefit of using CT models is likely to be limited.

To address these questions, we will utilise both simulated data and real observations. Simulated data will allow us to test different impacts of time-varying covariates on detectability under controlled conditions. This approach enables us to systematically vary the temporal precision of covariates measured at different time scales and the temporal scale at which these covariates fluctuate. For the case study, we will use real observations from a 5-months long camera trap study on an research center.

## 2 Material and methods

### 2.1 Models

We considered four types of site-based, mono-species, static hierarchical models (recapitulated in Table 1, Appendix S1: Figure S1). All models include a state process (occupancy or abundance) and a detection process (discrete- or continuous-time). Below, we briefly outline each component. A practical guide to our implementation of these models in Nimble (De Valpine et al., 2017), along with further explanation of the mathematical formulation, is provided in Appendix S1. The code is also available in a git repository, see supplementary material.

**Table 1:** The four compared models. With subscript *_i_* for “in site *i*”, subscript *_j_* for “during session *j*”, *z* the occupancy state, *ψ* the occupancy probability, *N* the abundance, *ϕ* the expected abundance, *µ* the species detection rate, *λ* the individual detection rate, *T* the duration of a session, *y* the count data.

| Model | State process | Detection process | Data |
| --- | --- | --- | --- |
| **CToccu** | *Occupancy*<br>$z_i \sim \text{Bernoulli}(\psi_i)$ | *Continuous-time*<br>If $z_i = 1$: Poisson process<br>with rate $\mu(t)$ | Time-to-each-detection |
| **DToccu** | *Occupancy*<br>$z_i \sim \text{Bernoulli}(\psi_i)$ | *Discrete-time*<br>$y_{i,j} \sim \text{Poisson}(\mu_{i,j}T_{i,j}z_i)$ | Detection<br>counts |
| **CTabun** | *Abundance*<br>$N_i \sim \text{Poisson}(\phi_i)$ | *Continuous-time*<br>If $z_i = 1$: Poisson process<br>with rate $\lambda(t)$ | Time-to-each-detection |
| **DTabun** | *Abundance*<br>$N_i \sim \text{Poisson}(\phi_i)$ | *Discrete-time*<br>$y_{i,j} \sim \text{Poisson}(\lambda_{i,j}T_{i,j}N_i)$ | Detection<br>counts |

#### 2.1.1 State process

The state process describes the latent presence or abundance at each site. We use single-season (static) models, which do not account for temporal changes in occupancy or abundance. In both cases, we assume that sites are independent. Site-level covariates can be included through a linear model, using the appropriate link function. As we focus on static models, these covariates are assumed to be constant over time and vary only across space.

##### Occupancy

The occupancy state of site *i*, *z_i_* (present = 1, absent = 0), follows a Bernoulli distribution with parameter *ψ_i_*, the occupancy probability: *z_i_* ∼ Bernoulli(*ψ_i_*). We assume that there are no false positives. If the species is detected at least once in a site, that site is considered occupied. We can model the occupancy probability as a logit-linear function of site covariates: logit(*ψ_i_*) = *α*^⊤^*v_i_*, where *α* is the coefficients vector and *v_i_* = (1, state cov_1_*_,i_, …,* state cov*_n,i_*)^⊤^ is the vector of state covariate values at site *i*, including an intercept.

##### Abundance

The population size at site *i*, *N_i_*, follows a Poisson distribution with expected abundance *ϕ_i_*: *N_i_* ∼ Poisson(*ϕ_i_*). This distribution could be replaced by another non negative discrete distribution, such as the negative binomial or its zero-inflated variant, when more appropriate (Joseph et al., 2009; Stoklosa et al., 2022). We model *ϕ_i_* as a log-linear function of site covariates: log(*ϕ_i_*) = *γ*^⊤^*w_i_*, where *γ* is the coefficients vector and *w_i_*= (1, state cov_1_*_,i_, …,* state cov*_n,i_*)^⊤^ is the vector of state covariate values at site *i*, including an intercept.

#### 2.1.2 Detection process

##### Classical detection representation

Conditioned on the ecological state, classical occupancy models (MacKenzie et al., 2002) and N-mixture models (Royle, 2004) typically represent detection using Bernoulli or Binomial random variables. In these models, the detection probability during a visit is denoted by *p*. However, this probability is inherently linked to the visit duration: shorter visits typically lead to smaller *p* values. This relationship creates a challenge when combining data collected under different survey protocols, especially those with varying visit durations.

To incorporate visit duration explicitly, the detection process can be modelled using an instantaneous detection rate *µ*(*t*). This is defined such that the probability of a detection occurring around time *t* in a small interval of duration *h* is approximately *µ*(*t*)*h*. Assuming *µ*(*t*) is constant over time, this formulation yields a cumulative detection probability over a visit of duration *T* given by *p* = 1 − *e*^−^*^µT^*. This representation provides a principled way to account for heterogeneous visit durations in detection models. In all models considered below, detection is described by a detection rate. We define rates per day for consistency. In occupancy models, we use the **species detection rate** *µ*, representing the average number of detections of the species per day. In abundance models, we use the **individual detection rate** *λ*, representing the average number of detections per individual per day.

##### Discrete-time detection process

DT models aggregate time-varying data per session, modelling detection counts *y_i,j_* at site *i* and session *j*. These detection counts are assumed to follow a Poisson distribution: *y_i,j_* ∼ Poisson(*µ_i,j_T_i,j_*) in occupancy models and *y_i,j_* ∼ Poisson(*λ_i,j_T_i,j_*) in N-mixture models, with *T_i,j_* the duration of session *j* at site *i*. Detection rates with covariates are modelled as log(*µ_i,j_*) = *β*^⊤^*x_i,j_* or log(*λ_i,j_*) = *β*^⊤^*x_i,j_*, where *β* is the detection coefficients vector and *x_i,j_* = (1, det_cov_1_*_,i,j_, …,* det_cov*_n,i,j_*)^⊤^ is the vector of detection covariate values at site *i* and session *j*.

##### Continuous-time detection process

In CT models, we use detection times, often transformed into interdetection times, also referred to as time-to-each-detection data (Guillera-Arroita et al., 2011; Haines et al., 2023). These are modelled as an inhomogeneous Poisson point process with a time-varying detection rate. Covariates must therefore be defined continuously over time. The instantaneous detection rate at time *t* at site *i* follows a log-linear model: log(*µ_i_*(*t*)) = *β*^⊤^*x_i_*(*t*) for occupancy and log(*λ_i_*(*t*)) = *β*^⊤^*x_i_*(*t*) for abundance, where *β* is the detection coefficients vector and *x_i_*(*t*) = (1, det_cov_1_*_,i_*(*t*)*, …,* det_cov*_n,i_*(*t*))^⊤^ is the vector of detection covariate functions at time *t* at site *i*, including an intercept.

##### Similarities and differences between CT and DT formulations

In the CT detection process, the number of detections over a deployment of duration *T* follows a Poisson distribution with parameter equal to the integral of the detection rate: 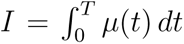 (or 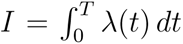 in abundance models), where *I* denotes the total detection intensity. If the detection rate is constant (*i.e.*, homogeneous Poisson process), this simplifies to *I* = *µT*, and detections follow Poisson(*µT*). When *µ*(*t*) varies, we can define an average detection rate 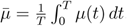, so that 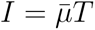. This equivalence shows that CT and DT models are mathematically identical when the detection rate is constant: both model detection counts as a Poisson process with parameter rate × duration (Zhang & Bonner, 2019). Thus, regression coefficients *β* are directly comparable across CT and DT models. When detection rates vary, differences arise not in model structure, but in how time-varying data –detections and detection covariates– are supplied. For instance, a DT model with 1-day sessions could be equivalent to a CT model with one covariate data point per day.

##### Advantage in using instantaneous detection rate

As noted above, if the instantaneous detection rate *µ* is constant over a session of duration *T*, the detection probability is *p* = 1 − *e*^−^*^µT^*. However, this non-linear relationship combined with the linear modelling of *µ* provides straightforward comparison of covariate effects across models. Furthermore, CT models require detection rates by construction, as probabilities inherently assume discrete sampling. Focusing exclusively on detection-rate-based models thus allows a consistent scale for covariate effects and enables direct comparison across CT and DT frameworks. Moreover, we argue that detection rates can be advantageous over probabilities – provided model identifiability holds (see Haines et al., 2023) – since they offer greater flexibility and remain meaningful regardless of discretisation scale (Pautrel et al., 2024).

### 2.2 Simulation study workflow

Figure 1 illustrates the steps of our simulation study, detailed in the following section. We repeated the simulation workflow 100 times and simulated data over a 3-month period.

**Figure 1:**
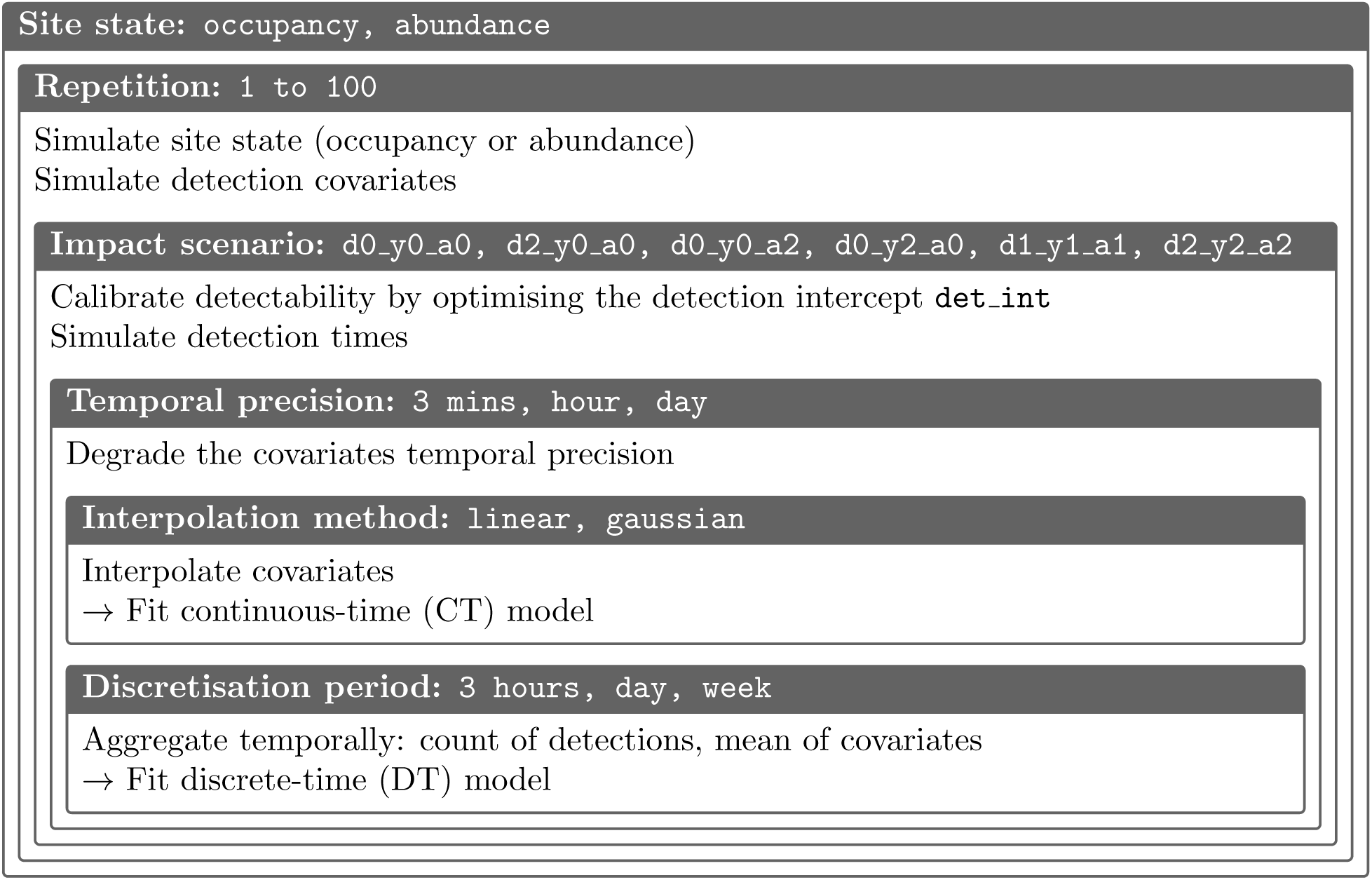
Overview of the simulation study workflow.

#### Sites state simulation

The occupancy or abundance at each site was simulated according to the models described in Section 2.1.1. We used the same parameter values for all sites, without state covariates: *ψ* = 0.5 for occupancy, and *ϕ* = 5 for abundance.

##### Detection covariates simulation

We simulated values for covariates every 3 minutes, ensuring that they were mutually uncorrelated, with three distinct temporal patterns: covday: constant across sites, daily and yearly variation (e.g. temperature-like); covyear: constant across sites, yearly variation only (e.g. seasonal signal); and covarma: site-specific, generated with an ARIMA model (AR = 0.9999, MA = 1) using stats::arima.sim in R (v4.4.0). All covariates were centered and scaled.

##### Detection covariate effect scenarios

We defined six scenarios by varying the strength of regression coefficients in the linear equation for detection rate, for three types of simulated covariates: covday, with daily variation (d); covyear, with yearly variation (y); and covarma, a site-specific ARIMA covariate (a). In each scenario, the detection rate intensity – *log*(*µ*(*t*)) for occupancy or *log*(*λ*(*t*)) for abundance – was defined as det_int + *β_d_*covday(*t*) + *β_y_*covyear(*t*) + *β_a_*covarma(*t*), where the coefficients *β_d_*, *β_y_*, and *β_a_* varied across scenarios (e.g. in scenario d2 y0 a0, *β_d_* = 2, *β_y_* = 0, and *β_a_* = 0).

##### Detection intensity calibration

Because covariate effects on detection were fixed within each scenario, we optimised the intercept parameter (det_int) to ensure comparability of detectability across scenarios. Specifically, we adjusted det_int so that the average number of detections matched 10 detections during the deployment per occupied site (occupancy model); 2 detections per individual (abundance model).

##### Observation simulation

Detection events were then simulated in continuous time, using rejection sampling to generate inhomogeneous Poisson point processes (Illian, 2008, p. 119–120). The intensity function used in this process was defined by the covariates and their scenario-specific coefficients, as described above, with linear interpolation between 3-minute time points.

##### Temporal degradation of covariates

To assess how coarser temporal resolutions affect model performance, we compared three levels of covariate resolution: using all values at 3-minute intervals, or degrading them by subsampling one value per hour or per day. Only the covariate values at the retained time points were kept. This degradation aimed to evaluate the limits of CT models under varying data availability and to test how the loss of fine-scale covariate variation influences inference accuracy.

##### Discretisation

For the DT models, the detections were aggregated into counts per site and session. Covariates were summarised by their mean within each session. Sessions were defined as 3-hour periods, days, or weeks. This allowed us to assess how well DT models perform when fitted on increasingly coarse temporal units, in line with the assumption that the temporal scale of aggregation may affect the accuracy of inference. Notably, there is an equivalence between approaches: in CT models, interpolated covariates are integrated over time, and the integral of a covariate over a session of length *T* is mathematically equivalent to the mean value of that covariate during *T*, multiplied by *T*.

##### Covariate interpolation

For the CT models, covariates, whether degraded or not, were interpolated to provide continuous input over time. Two interpolation methods were used: linearly between successive points, and with a Gaussian process (via gausspr from the R package kernlab, v0.9.33) (Karatzoglou et al., 2004). For the latter, we used the default parameters, which resulted in a smoothed interpolation (*e.g.* Appendix S2: Figures S19, S16 for the use case). The aim was to explore how different interpolation approaches might influence model performance, given the assumption that interpolation quality can affect CT model results.

##### Model fitting

Both DT and CT models were fitted using the R package nimble (v1.3.0) (De Valpine et al., 2017). Initial values were roughly estimated from the simulated data (see Appendix 2: Section S2.1). We applied a burn-in period of 7,500 iterations, followed by 50,000 additional iterations for occupancy models and 100,000 for N-mixture models. Three chains were run with a thinning interval of 25. Analyses were distributed across three Linux Debian 12 computing servers, with hardware specifications detailed in Appendix S2: Section S2.1.

##### Model evaluation

For each simulation scenario, we calculated four complementary metrics. For point estimates (defined as the median of the posterior distribution), we compared the estimated value to the simulated parameter using root mean square error (RMSE) and bias. To assess the posterior distribution’s uncertainty, we measured coverage (the proportion of simulations where the simulated parameter was within the 95% credible interval) and the average range of the 95% credible interval (ARCI) (Pautrel et al., 2024).

### 2.3 Case study

#### 2.3.1 Study area

The case study was conducted in the Cadarache Center of the French Alternative Energies and Atomic Energy Commission (CEA), located in the French Mediterranean region (Bouches-du-Rhône). This site covers 867 hectares and hosts around 2,400 workers. The area was previously used as agricultural land, then as a game reserve for hunting (1924–1958), and, since 1963, as a nuclear research centre, and has been fully enclosed ever since. Since the perimeter fence is regularly monitored and maintained, large species cannot move in or out of the study area, meaning there is no exchange with external populations.

A vector land-use map (Appendix S2: Figure S10) combined multiple data sources, including Corine Land Cover (Büttner et al., 2017), RPG (IGN, 2020), and BD TOPO (IGN, 2021), along with photointerpretation and dedicated fieldwork. The compilation began with the coarser Corine Land Cover, which provides full coverage of the study area, and progressively incorporated more detailed layers. Habitat types were grouped into six broad classes: aquatic (covering 0.5% of the study area), without vegetation (3.7%), shrubby (8.5%), urban (18.5%), herbaceous (25.4%) and woody (43.4%).

To guide camera placement, a preliminary landscape analysis ensured coverage of the site’s habitat diversity. Final placement was constrained by access rights due to the site’s military use (Appendix S2: Figure S10). Cameras were angled slightly downward to detect both small ground-dwelling and larger species, while avoiding vegetation that could cause false triggers. Efforts were made to maintain a consistent field of view across devices. In total, 59 cameras were deployed from June to November 2023, operating for 64 to 143 days each (mean = 135), totalling 7,994 trap-days.

#### 2.3.2 Selecting focal taxa

Camera traps recorded 795,159 images, initially processed with MegaDetector v5. A first filter removed 346,231 images without any detection box, even at low confidence. The remaining 448,928 images were uploaded to the collaborative annotation platform OCAPI (https://ocapi.terroiko.fr/). OCAPI used an artificial intelligence (AI) pipeline based on MegaDetector v5 to detect animals, vehicles, and humans. Animal boxes predicted by MegaDetector are then cropped and classified with FaunIA, a model based on Efficient-NetV2 (Moulherat et al., 2024; Tan & Le, 2021).

To identify which species were predicted reliably enough, a subset of 9,180 images was manually annotated to build a confusion matrix (Appendix S2: Figure S5). Since the models account for imperfect detection but not false positives, selection focused on taxa with high precision. To improve reliability, detections were grouped into three-image sequences per trigger event. Only events where the target taxon was predicted by the AI in all three were retained. Detection rates in the models thus combines camera trap triggering and AI detection.

We selected four taxa with over 92 % image-level precision and over 98 % sequence-level precision (Appendix S2: Tables S1, S2): wild boar (*Sus scrofa*, 5,219 detections on 58 cameras), mouflon (*Ovis aries*, 4,032 detections on 59), red fox (*Vulpes vulpes*, 689 detections on 59), and small mustelids (*Martes* and *Mustela* spp., grouped due to difficulty distinguishing them on camera-trap images, 109 detections on 32) (Appendix S2: Figures S6, S7, S8, S9). Using AI was not the main focus of our study, so we kept the approach simple. We recommend model users treat AI-derived outputs not as ground truth but as uncertain data, as poor handling of prediction uncertainty can lead to biased inference (Katsis et al., 2025; Monchy et al., 2025).

#### 2.3.3 Gridding the study area for site-based modelling

All focal taxa have wide home ranges. Wild boar can cover about 30 ha daily (Russo et al., 1997), with seasonal home ranges up to 400 ha (Fattebert et al., 2017). Mouflons also use up to 400 ha in spring (Dubois et al., 1992, 1993). Red fox home ranges vary widely between individuals, from 375 ha (Lucherini et al., 1995) to 350 km² (Walton et al., 2017). Small mustelid species have varying home ranges, with examples including 250 ha for *Martes martes* (Zalewski et al., 2004) and 2,900 ha for *Mustela lutreola* (Fournier et al., 2008).

Given these movement capacities, our 867-ha study area is too small to define spatially independent sampling units as required by standard occupancy or N-mixture models, which assume closure and independence across sites. We therefore divided the area into 1-ha hexagonal cells. As often in such contexts, we interpret model outputs as indicators of site use, rather than true occupancy or abundance (Nichols et al., 2008). This is an oversimplification for mobile species (Valente et al., 2024), but necessary in our case.

#### 2.3.4 Covariates

Covariates were grouped into three categories: spatial habitat covariates affecting site state (occupancy or abundance), which was assumed static and thus modelled using time-invariant covariates; temporal weather covariates influencing detection rate; and spatiotemporal anthropic disturbance covariates (per camera trap), which also affected detection.

We derived **habitat covariates** from the land-use map (Section 2.3.1). For each location and habitat type, we calculated distance to the nearest habitat polygon and percentage cover within buffers of 50, 150, 447, 1337, and 4000 metres. **Weather data** came from a single CEA station recording air temperature, pressure, humidity, wind direction and speed, and rainfall every minute. We assumed weather conditions were spatially uniform across the study area. For each variable, we computed rolling statistics over multiple windows (from 3 minutes to 7 days), including centred and trailing rolling means, maxima, and sums. We measured **anthropic disturbance** from human detections on cameras. Cumulative Gaussian and half-Gaussian kernel smoothing over various time windows (from 3 minutes to 90 days) captured both immediate and long-term disturbance.

These different time windows were used to characterise various temporal scales of variation in the raw covariates. They retain information useful for the model at different scales. Without these temporal variants, the model would lack the data necessary to identify potential punctual effects (*e.g.* an animal hiding during rain but soon reappearing) versus more sustained responses (*e.g.* lower activity and thus detectability during generally cold weather, regardless of brief warmer periods). However, variants of the same variable are often inter-correlated. To reduce dimensionality and collinearity, we summarised habitat, weather, and disturbance covariates separately via PCA, yielding four habitat, four weather, and two disturbance uncorrelated covariates (Appendix S2: Section S2.2.2).

#### 2.3.5 Modelling the taxa presence/abundance and detection rate

Our modelling framework followed the simulation study setup, varying temporal precision of covariates (3 minutes, 1 hour, or 1 day; Appendix S2: Figures S16, S19), interpolation method for CT models (linear or Gaussian, same figures), and discretisation period for DT models (3 hours, 1 day, or 1 week). For each, we fitted a model without covariates and one including all PCA-derived covariates. CT models at 3-minute intervals were run at 15-minute intervals instead, due to RAM limits that prevented model fitting. Site state (occupancy or abundance) was defined per cell, but because some cells had multiple cameras (50 with one, two with two, one with five), detection was modelled per camera.

## 3 Results

### 3.1 Simulation study

#### 3.1.1 Coefficient estimation

##### No covariate effect on detection

When detection covariates had no effect, all models performed well. State estimates were near their simulated values. Occupancy estimates ranged from –0.612 to 0.653 on the logit scale (backtransformed: 0.35–0.66). Abundance estimates ranged 1.140–1.878 on the log scale (backtransformed: 2.13–6.54). RMSE remained low and similar across scenarios (Figure 2). Higher temporal precision improved RMSE and bias for covday (Appendix S2: Figure S1). Coverage exceeded 90% for occupancy and 82% for abundance (Appendix S2: Figure S2). Credible intervals stayed narrow (Appendix S2: Figure S3). Abundance models showed slight bias, underestimating abundance and overestimating detection intercepts, likely due to known identifiability issues in Poisson–Poisson models without covariates (Haines et al., 2023).

**Figure 2:**
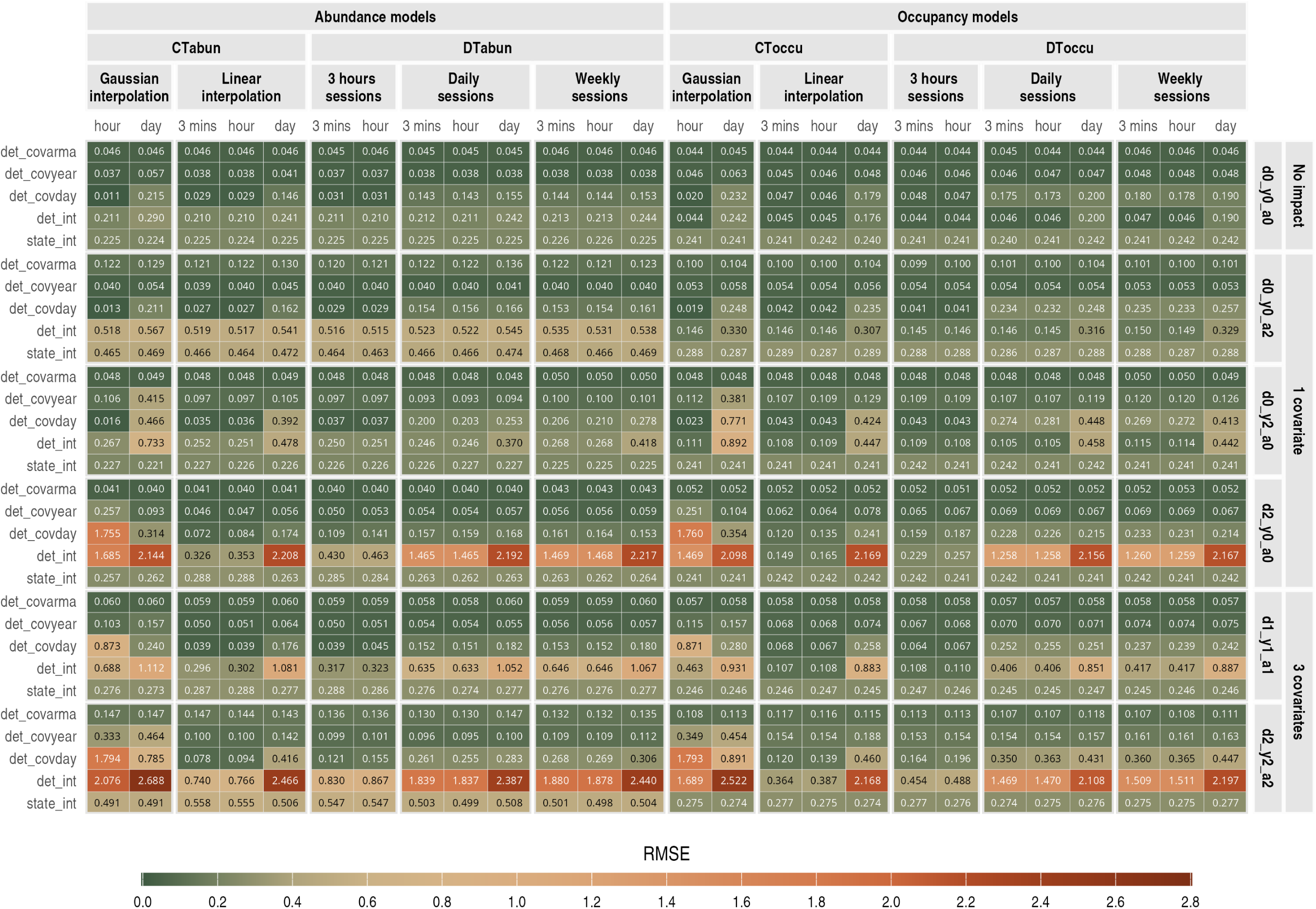
Root mean square error (RMSE) for occupancy and abundance models. Rows correspond to model terms. Columns represent model types and configurations, as indicated in the top facets: occupancy or abundance; CT or DT; interpolation method (CT) or discretisation (DT); and covariate temporal resolution (one value every 3 minutes, hour, or day). Right-hand facets show different simulated detection covariate effect scenarios, as described in Section 2.2. For example, scenario d2 y0 a0 indicates a coefficient of 2 for covday (d2) on the detection rate, with no effect of covyear (y0) or covarma (a0). Cell colours and values indicate the RMSE.

##### Effect of covarma

When detection depended on the covarma covariate, bias and RMSE increased in abundance models (Appendix S2: Figure S1). Coverage dropped to 54–60% (abundance intercept) and 28–60% (detection intercept), with wider credible intervals (Appendix S2: Figures S2, S3). Occupancy models were less affected.

##### Effect of covyear

When only covyear influenced detection, model performance stayed close to the no-effect scenario. Models capturing finer temporal structure performed slightly better (Figure 2, Appendix S2: Figures S1, S2).

##### Effect of covday

Detection performance depended heavily on retaining fine-scale variation. The detection intercept’s RMSE was the lowest with 1-hour or 3-minute temporal resolution for CT models with linear interpolation or DT models with 3-hour sessions (0.149–0.463). It was above 1.258 in other cases, and exceeded 2 with daily temporal resolution (Figure 2). Coverage dropped to zero in models unable to capture the fine-scale temporal variation (Appendix S2: Figure S2). The covday coefficient was still generally well estimated, likely thanks to its yearly pattern. CT models with linear interpolation showed better coverage than DT models with 3-hour sessions, even when RMSE was similar, indicating more appropriate uncertainty quantification.

##### Effect of multiple covariates

With all three covariates affecting detection, bias and uncertainty increased, especially with stronger effects. Poor retention of fine-scale temporal information caused the largest performance drops.

#### 3.1.2 Fitting metrics

##### Convergence quality

Trace plots confirmed sufficient burn-in. Most models converged well, though a few had only marginally acceptable diagnostics. Effective sample sizes ranged from 1878–6000 (5th–95th percentile) across all occupancy model parameters, and from 672–11904 for abundance (N-mixture) models. The R^ remained below 1.007 (occupancy) and 1.084 (abundance).

##### Fitting duration

Fitting times ranged from 1 minute 27 seconds to 173 hours 37 minutes. Continuous-time models took longer to fit when covariates had finer temporal precision, due to the increased number of steps in the Riemann sum used to approximate the detection rate (Appendix S2: Figure S4). Discrete-time models took longer to fit as the number of sessions increased (Appendix S2: Figure S4).

### 3.2 Case study

#### 3.2.1 Models without covariates

Models without covariates gave consistent results across model types (CT or DT) and DT discretisation scales (Appendix S2: Figures S20, S21, S23, S25, S27, S29, S31, S33).

#### 3.2.2 State parameters estimation

Habitat covariates rarely influenced occupancy or abundance (Appendix S2: Figures S22, S26, S28, S30, S32, S33). Estimates were robust to detection variation. For boar, mouflon, and fox occupancy, nearly all sites had detections, leading to occupancy probabilities close to 1 and thus independent of covariates (Figures S22, S26, S30). Covariates had little impact on small mustelid occupancy or abundance (Figures S34, S32) and fox abundance (Figure S28). Wild boar abundance (Figure 3) was negatively linked to abun[1] (-0.462 to -0.195), and positively to abun[2] (0.106 to 0.429) and abun[4] (0.163 to 0.410). Mouflon abundance (Appendix S2: Figure S24) showed similar trends: abun[1] had a negative effect (-0.273 to 0.001), abun[2] a positive one (0.242 to 0.547). Dimensions are interpretable from PCA (Appendix S2: Figures S11, S13): abun[1] opposes urban (low) to wooded/shrubby (high) cover; abun[2] opposes wooded (low) to aquatic/herbaceous (high) areas; abun[4] contrasts

**Figure 3:**
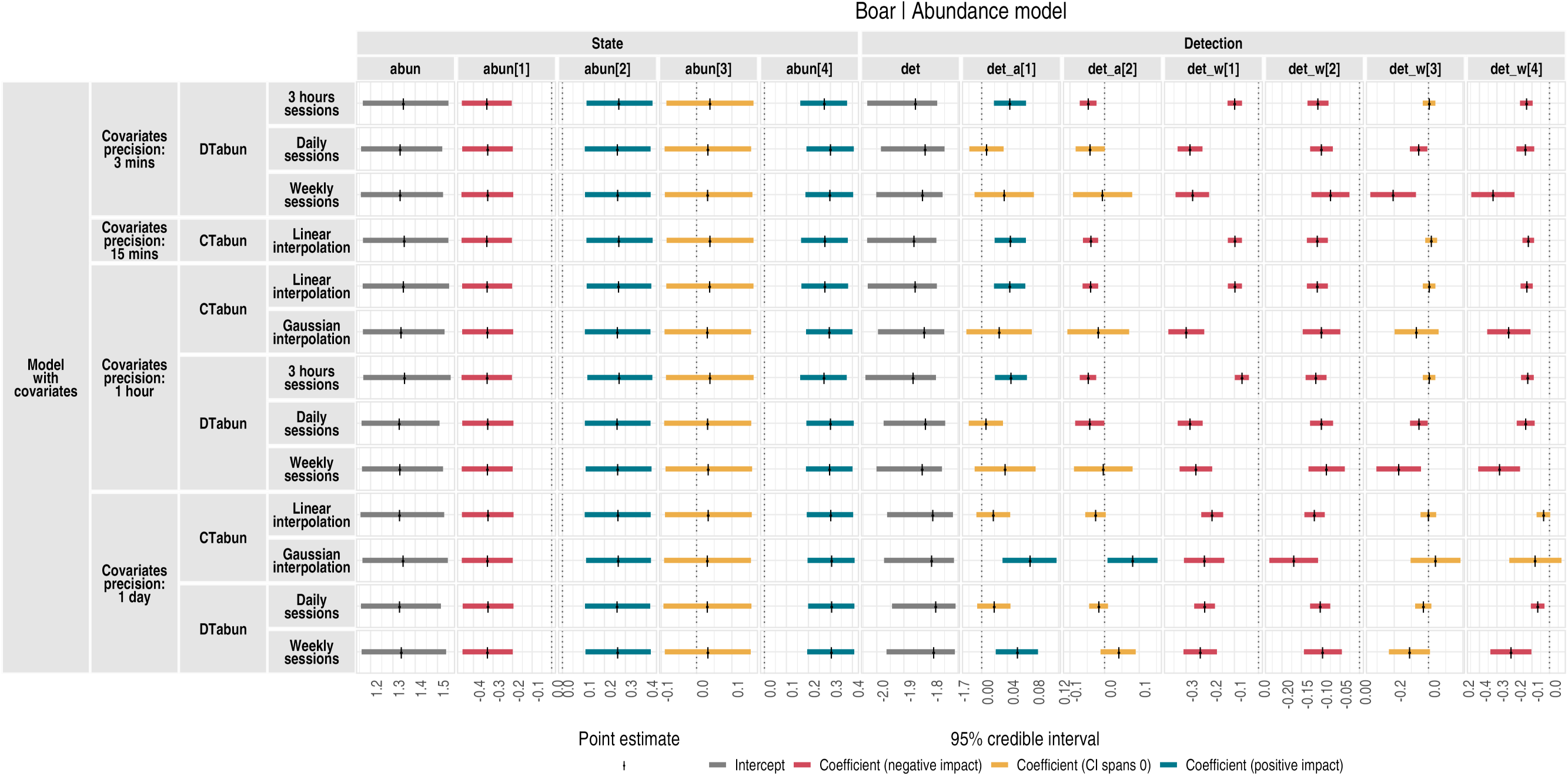
Abundance models results for the wild boar nearby urban areas (low) with aquatic proximity and shrubby/wooded structure at moderate distance (high).

#### 3.2.3 Detection parameters estimation

Detection estimates varied with how temporal covariates were processed. Results depended on covariate resolution, DT discretisation period, and CT interpolation method. Occupancy and abundance models with the same covariate processing produced similar estimates, but the level of temporal detail retained produced different results.

##### Retaining fine-scale temporal detail can increase covariate coefficient estimates

Some covariate effects were stronger when fine-scale detail was retained: at least hourly covariate measurements, CT with linear interpolation, or DT with 3-hour sessions. Effects weakened when covariates were smoothed (e.g., Gaussian interpolation or daily/weekly aggregation). For instance, det_a[1] negatively impacted fox detection only with high temporal detail (Appendix S2: Figures S30, S28). This suggests that human presence briefly reduces fox detectability. This effect is revealed only when both covariates and detections retain fine temporal resolution. If these are smoothed or aggregated, the short-term effect is masked or lost, preventing detection of the fine-scale correlation between the covariate and reduced detectability.

##### Retaining fine-scale temporal detail can decrease covariate coefficient estimates

In contrast, some effects increased with smoothing. In wild boar models (Figure 3, Appendix S2: Figure S22), det_w[1] (positively correlated with temperature and wind, negatively with hygrometry; Appendix S2: Figures S14, S15) showed a stronger negative effect on detection when smoothed. This implies either *(i)* prolonged dry, hot, windy conditions reduce detectability or *(ii)* effects occur near time *t*, not necessarily at *t*.

##### Temporal scale of covariate can affect both coefficient estimate sign and strength

In certain cases, the impact of a covariate on detection varied depending on the scale of temporal processing not only in strength, but also in direction. For small mustelids (Appendix S2: Figures S32, S34), the effect of det_w[1] was negative with fine-scale resolution, null at intermediate, and positive with heavy smoothing.

## 4 Discussion

Performance and relevance of CT over DT models depend on the modelling aim and on methodological choices beyond just CT vs DT. Occupancy and abundance were estimated consistently, but detection parameters depended strongly on the temporal characteristics of detection covariates and the quality of their interpolation, in both occupancy and N-mixture models.

In simulations, bias, higher error, and lower coverage arose when fine-scale covariate variation affected detection but was poorly retained. This occurred with limited data points in time, coarse discretisation for DT models, or heavily smoothed interpolation for CT models. When the covariate had broad-scale variation, no effect, or was well preserved, models performed similarly. N-mixture models were sensitive to identifiability issues when covariates had no effect (Haines et al., 2023), or when using site-varying covariates, likely due to detection–abundance confounding.

Our case study supported these results, although we lacked ground truth to assess which model was closest to reality. Detection estimates clearly varied across models (CT/DT, discretisation, interpolation, temporal resolution), while state estimates remained stable.

Retaining fine-scale variation led to mixed outcomes:

- In some cases, it increased the estimated covariate effect, which may reflect a punctual influence on detection.
- In other cases, it decreased the effect. This could mean that the covariate has a broader-scale effect, or that its influence on detection is shifted in time, either before or after the observed variation. Testing temporal offsets might help clarify this.
- In a few cases, the direction of the effect changed. This may reflect more complex or non-linear processes, potentially dependent on the temporal scale, that warrant further exploration.

### 4.1 Model recommendations with continuously collected data

#### 4.1.1 Modelling detectability can be a research focus

If detectability is modelled only to correct bias in occupancy or abundance estimates, model choice (among the tested ones) has limited impact. Both CT and DT approaches gave similar occupancy or abundance results under most conditions. In such cases, fine-scale modelling of detection may not be necessary. Efforts might be better directed at improving the state process, for example with dynamic or spatially explicit models.

In contrast, if detectability is the research focus, retaining temporal detail becomes important. Detectability is no longer a nuisance parameter but a process with ecological meaning. It relates to imperfect detection but also reflects activity patterns, behaviour, and responses to environmental conditions. Our simulation and case studies showed that modelling choices affect detectability estimates. In DT models, the discretisation interval matters. In CT models, the interpolation method plays a key role. We only tested two methods, selected arbitrarily and not tailored to the structure of the covariates. More sophisticated approaches, such as wavelet analysis, could better capture multi-scale signals (Cazelles et al., 2008). These choices can also inform study design—for instance, required covariate resolution may guide sensor selection (e.g. weather sensors with high-frequency output). Both simulations and the case study suggest that DT models with fine bins can approximate CT performance. As long as temporal detail is retained, choosing between CT and DT is not critical for studying detectability. Continuous-time alternatives such as Hawkes processes may offer further insights, for instance in predator–prey dynamics (Nicvert et al., 2024).

#### 4.1.2 Practical considerations

##### Software and implementation

DT models are widely supported and easier to implement. R packages such as unmarked (Fiske & Chandler, 2011; Kellner et al., 2023), spOccupancy (Doser et al., 2022), spAbundance (Doser et al., 2024), and ubms (Kellner, Fowler, et al., 2022) are flexible, user-friendly tools. CT models still require custom code, limiting accessibility. However, recent interest in continuous-time frameworks (Kellner, Parsons, et al., 2022; Priyadarshani et al., 2024; Rushing, 2023) is driving software development perspectives (Kellner et al., 2023).

##### Computation time

Computation times remained manageable for most models, though they increased with finer temporal precision. In our simulation study, CT models with 3-minute precision took between 8 hours 50 minutes and 173 hours 36 minutes; DT models with 3-hour sessions took between 56 minutes and 27 hours 36 minutes. Other models ran between 1 minute and 6 hours 49 minutes (Appendix S2: Figure S4). CT and DT models both had longer runtimes at higher temporal resolution, so runtime is unlikely to be a main driver of model choice between them.

##### Memory use

Memory demand was limiting for CT models at high covariate resolution. In our case study, we had to reduce resolution from 3 to 15 minutes, even on servers with over 500 GB RAM. For large datasets (many sites, long periods, or dense covariates), memory-efficient implementation will be essential. Alternatively, users can reduce resolution or partition the analysis (e.g. by season).

### 4.2 Suggestions for future directions

#### Embedding interpolation within model fitting

An interesting direction for CT models could be to integrate the interpolation step directly into the model-fitting process, effectively adding another layer to the hierarchical framework. Although we did not test this, our initial thought was that it could reduce the influence of arbitrary methodological choices and support a more objective modelling process. This, in turn, could improve comparability across studies and help develop a more generalisable framework that adapts to diverse contexts. However, results from our simulation study suggest this may be less straightforward. The effect of fine-scale covariate temporal resolution on detectability estimates was inconsistent. It could increase, decrease, or even reverse the estimated effect. This implies that multiple equally plausible optimised equilibrium states might exist. This complexity may limit the benefits of integrating the interpolation directly into the model.

#### Improving convergence and MCMC sampling strategies

In some models, convergence diagnostics were only marginally acceptable. To address this, we increased the number of iterations post burn-in for our abundance models and our case study (100,000 iterations vs 50,000 in occupancy models). However, alternative strategies could be considered to improve convergence more efficiently, such as block sampling or changing the sampling algorithm (*e.g.* Hoffman and Gelman, 2014), which we did not explore in this study, keeping the default Nimble assignment of sampler algorithms. These avenues may offer gains in computational performance or stability.

#### Alternative methods for integral estimation

In the CT model, the integral of the detection rate was estimated using a Riemann sum. Although we tested three levels of temporal resolution (3-minute, 1-hour, 1-day steps), we did not explore alternative numerical integration methods. These could influence the effective temporal precision that the model can retain and might further impact detection estimates. Exploring different integration methods could offer improvements in both computational efficiency and modelling accuracy, particularly if they allow to better capture how covariates vary rapidly at a fine temporal scale.

## Acknowledgments

This work was supported by the Association Nationale de la Recherche et de la Technologie (PhD Grant CIFRE N°2022/0428). This work is part of the PSI-BIOM project granted by the French PIA 3 under grant number 2182D0406-A. This research is partly funded by Biodiversa+, the European Biodiversity Partnership, in the context of the Big Picture project under the 2022-2023 BiodivMon joint call. It is co-funded by the European Commission (GA No. 101052342) and the French Agence Nationale de la Recherche.

## Author Contributions

**Léa Pautrel**: Conceptualisation, Data Curation, Formal Analysis, Methodology, Visualisation, Writing – Original Draft Preparation. **Sylvain Moulherat**: Conceptualisation, Methodology, Supervision, Writing – Review & Editing. **Benoit Charrasse**: Resources (fieldwork), Writing – Review & Editing. **Guillaume Debat**: Resources (AI), Writing – Review & Editing. **Lucie Gendron**: Data Curation (land-use map), Writing – Review & Editing. **Kenneth Kellner**: Methodology, Writing – Review & Editing. **Marie-Pierre Etienne**: Conceptualisation, Methodology, Supervision, Writing – Review & Editing. **Olivier Gimenez**: Conceptualisation, Methodology, Supervision, Writing – Review & Editing.

