## Appendix 1 for "When does temporal resolution matter? Including detection covariates in discrete- versus continuous-time occupancy and N-mixture models"

### Appendix S1: Practical guide for implementing discrete- and continuous-time hierarchical models in Nimble

#### When does temporal resolution matter? Including detection covariates in discrete- versus continuous-time occupancy and N-mixture models

Léa Pautrel[1, 2], Sylvain Moulherat[2], Benoit Charrasse[3], Guillaume Debat[2], Lucie Gendron[2], Kenneth Kellner[4], Marie-Pierre Etienne[5, *], and Olivier Gimenez[1, *]

[1]Centre d'Ecologie Fonctionnelle et Evolutive (CEFE), University of Montpellier, CNRS, EPHE, IRD, 1919 Route de Mende, 34000 Montpellier, France
[2]OïkoLab, TerrOïko, 2 Place Dom Devic, BP 26, 81540 Sorèze, France
[3]CEA, DES, IRESNE, DTN, Laboratory for Environmental Transfer Modeling, Cadarache, 13108 Saint-Paul-lès-Durance, France
[4]Department of Fisheries and Wildlife, Michigan State University, 480 Wilson Rd, East Lansing, Michigan, USA
[5]Univ Rennes, Ensai, CNRS, CREST – UMR 9194, F-35000 Rennes, France
[*]Co-senior authors

01 September 2025

This appendix is not intended as a general introduction to using Nimble. For that purpose, the official Nimble documentation (https://r-nimble.org/documentation) is an excellent resource. We also do not aim to provide a full overview of Bayesian methods (see e.g., King et al., 2010), nor a general tutorial on hierarchical modelling.

In this guide, we briefly introduce the implementation of discrete-time hierarchical models with count data, which are well documented in the literature and serve as a baseline. We then focus on the more specific issue of **implementing continuous-time hierarchical models in Nimble**. In our experience, examples of such models are rare, at least in ecology, where most resources focus on discrete-time formulations. We encountered challenges bridging this gap while developing the models presented in this paper, which motivated writing this guide.

To help others facing similar questions as we encountered, we provide this guide as a complement to the code, which is also available in a git repository: oikolab.terroiko.fr/...occupancy-and-n-mixture-models. We have aimed to keep explanations accessible for ecological modellers who are more familiar with discrete-time approaches. We extract and explain the most relevant components related to continuous-time implementation, presenting them in a standalone format. Our aim is to offer a clearer starting point for those wishing to implement such models in Nimble, or in similar Bayesian frameworks such as JAGS or BUGS, without needing to explore the full code provided with our paper, which contains other code related to the simulation study and the case study. We also include mathematical explanations behind our implementation choices, clarifying why the approach works.

We describe only the method used in this paper. To our knowledge, no established alternative resources currently exist for implementing these continuous-time hierarchical models in a Bayesian framework. However, we believe that more elegant or optimised solutions are possible, and we hope this guide can serve as a useful basis for future developments.

For those interested in implementing these models within a **frequentist framework**, we recommend reading Haines et al. (2023), who describe the likelihood for N-mixture models depending on the detection data: discrete-time with binary or count data (*count data corresponding to the discrete-time models considered here*), time-to-first-detection, and time-to-all-detection approaches (*the latter corresponding to what we refer to here as continuous-time models*). Their work does not, however, directly incorporate detection covariates. For models that include covariates, we refer to Guillera-Arroita et al. (2011), who explain how to include continuous covariates in the likelihood for a detection process modelled continuously along a transect. Although their formulation uses space (transect length), the mathematical structure is equivalent to a temporal process and can therefore be directly applied to continuous-time detection.

We present here the Nimble code used for our case study. The code used in the simulation study is nearly identical, with the only differences relating to covariates, and the detection process (which is defined per site in the simulation study, but per camera trap and site in the case study, since some sites had several camera traps). We follow the same nomenclature as in the main text: `DT` stands for discrete-time, `CT` for continuous-time, `occu` refers to occupancy models, and `abun` refers to abundance models using an N-mixture approach. The four models (`DToccu`, `CToccu`, `DTabun`, and `CTabun`) are described in the paper. Figure S1 provides another representation of those hierarchical models.

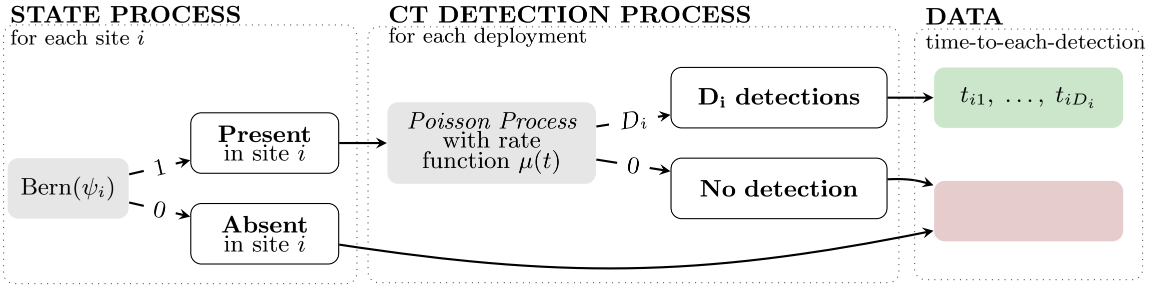


(a) CToccu. Continuous-time occupancy model

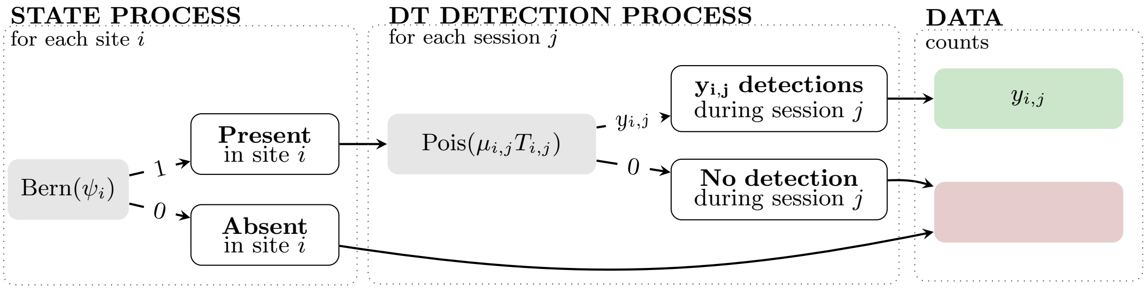


(b) DToccu. Discrete-time occupancy model

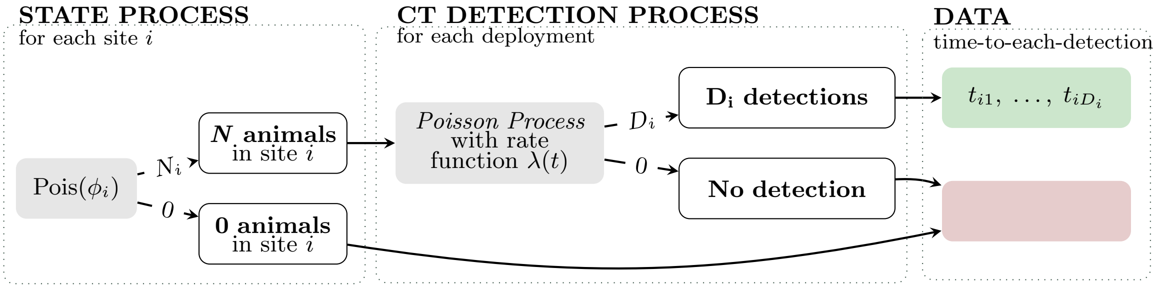


(c) CTabun. Continuous-time abundance model

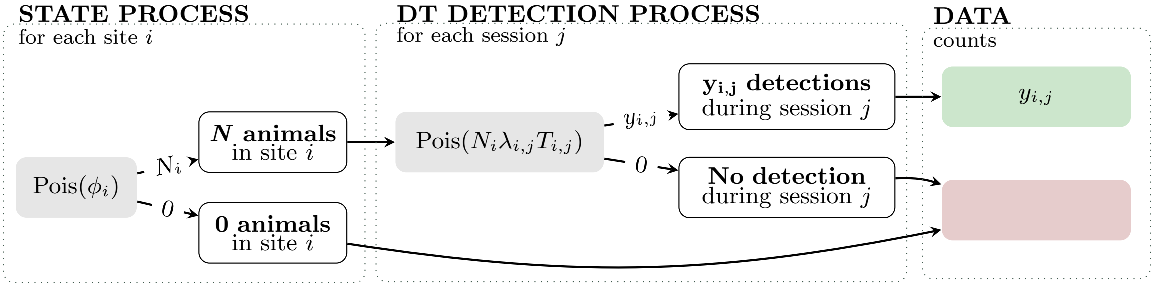


(d) DTabun. Discrete-time abundance model (Poisson-Poisson N-mixture)

Figure S1: **Models formulation**. With: subscript $_i$ as "in site $i$", subscript $_j$ as "during session $j$" (a **session** as a discretised time interval), a **deployment** as an uninterrupted monitoring period by a given sensor, $\psi$ the occupancy probability, $\phi$ the expected abundance, $\mu$ the species detection rate, $\lambda$ the individual detection rate, $T$ the duration of a session, $N$ the abundance, $D$ the total count of detections, $y$ the discretised count of detections, $t$ the time of a detection (easily transformed to interdetection times, or time-to-each-detection).

#### S1.1 Continuous-time detection process with a non-homogeneous Poisson process

The most common approach to model detection is to use discrete-time (DT) models. In these, temporally varying data are aggregated into fixed-length sessions. Detections are typically reduced to binary data indicating presence or absence within each session, from which a detection probability is modelled. However, in our study, we considered detection counts, the number of detection events summed within each session. These counts are then modelled using a Poisson distribution, and any time-varying covariates influencing detectability are summarised over the session (*e.g.*, by the mean, median, standard deviation...).

In contrast, continuous-time (CT) models do not require aggregation. We model detections as a non-homogeneous Poisson process (NHPP), also referred to as an inhomogeneous Poisson point process. This framework allows the detection rate to vary continuously over time, described by a function $\lambda(t)$ that depends on time-varying covariates. For further background on NHPP, we refer to Drazek (2013), Illian (2008), and Ross (1996). Below, we focus only on the key properties relevant to our implementation.

Let:

- $N$ be the number of detections during the deployment,
- $\lambda(t)$ the detection rate at time $t$,
- $\Lambda = \int_D \lambda(t)\, dt$ the integrated detection rate over deployment $D$,
- $\{t_1, \dots, t_n\}$ the observed detection times.

The NHPP has two key properties:

1. $N \sim \text{Poisson}(\Lambda)$: the count of events in $D$ follows a Poisson distribution with rate $\Lambda$.
2. Conditional on the count of events $N = n$, the detection times are independent and identically distributed with density $f(t_i) = \lambda(t_i)/\Lambda$.

Combining these gives the likelihood of observing $n$ detections at times $t_1, \dots, t_n$:

$$
\begin{aligned}
L(\lambda; \{t_1, \dots, t_n\}) &= \underbrace{e^{-\Lambda}\frac{\Lambda^n}{n!}}_{\text{Poisson count}} \times \underbrace{\prod_{i=1}^{n} \frac{\lambda(t_i)}{\Lambda}}_{\text{Conditional times}} \\
&= \frac{e^{-\Lambda}}{n!} \prod_{i=1}^{n} \lambda(t_i)
\end{aligned}
$$

The $1/n!$ disappears if the times are ordered (which is the case in our implementation), as ordering the sample multiplies the likelihood by $n!$ (Ross, 1996, pp. 66–67). Thus, the simplified likelihood becomes:

$$
L(\lambda) = e^{-\Lambda} \prod_{i=1}^{n} \lambda(t_i)
$$

##### Connection to implementation in Nimble

We calculate the detection rate at punctual times over fixed interval $m$ (e.g., per minute), depending on the detection covariates (in the example below, we use only one detection covariate for simplification).

$$
log(\lambda_m) = log(\lambda(t_m)) = \beta_0 + \beta_1 \cdot \text{covariate}_m
$$

The corresponding Nimble code is:

```
# Detection rate per minute
for (m in 1:nb_min) {
  log(lambda_min[m]) <- beta0 + beta1 * covar_min[m]
}
```

Then, we approximate $\Lambda = \int_D \lambda(t)\, dt$ the integral of the detection rate over the whole deployment, using a Riemann sum over regularly spaced intervals:

$$\Lambda \approx \sum_m \lambda_m \cdot \Delta t$$

The corresponding Nimble code is:

```
# Riemann sum: integral of the detection rate during the whole deployment
Lambda <- sum(lambda_min[1:nb_min]) * (1/60)
```

The total number of detections is modelled via a Poisson draw: `Ntot ~dpois(lambda = Lambda)`.

Each detection time $t_i$ is modelled via a Bernoulli draw with success probability $\lambda(t_i)/\Lambda$, using a ones trick: we condition on observing 1, allowing us to evaluate the density at $t_i$ without having to sample it. The corresponding Nimble code is:

```
# Intensity at detection times
for (i in 1:Ntot) {
  log(lambda_evt[i]) <- beta0 + beta1 * covar_evt[i]
  det_evt[i] ~ dbern(prob = lambda_evt[i] / Lambda)
}
```

The full Nimble code for this minimal NHPP example is thus presented in Model 1.

Model 1: Non homogeneous Poisson Process in Nimble

```
nhpp_code <- nimble::nimbleCode({
  # Priors
  beta0 ~ dnorm(0, sd = 1)
  beta1 ~ dnorm(0, sd = 1)

  # Detection rate per minute
  for (m in 1:nb_min) {
    log(lambda_min[m]) <- beta0 + beta1 * covar_min[m]
  }

  # Riemann sum: integral of the detection rate during the whole deployment
  Lambda <- sum(lambda_min[1:nb_min]) * (1/60)

  # Total count of detections
  Ntot ~ dpois(lambda = Lambda)

  # Intensity at detection times
  for (i in 1:Ntot) {
    log(lambda_evt[i]) <- beta0 + beta1 * covar_evt[i]
    det_evt[i] ~ dbern(prob = lambda_evt[i] / Lambda)
  }
})
```

Where:

- `covar_min[m]` is the covariate value at minute $m$,
- `lambda_min[m]` is the detection rate at minute $m$,
- `covar_evt[i]` is the covariate at detection time $t_i$,
- `det_evt[i]` is always 1, used for the one-trick in the Bernoulli draw.

The data provided for CT models thus differs substantially from that used in DT models. For this reason, we explain below not only the model structures but also give guidance on data preparation and formatting.

#### S1.2 Discrete-time occupancy model without covariates

Model 2 gives the code used for our discrete-time occupancy NULL model, without covariates. Occupancy status is modelled at the site level, while detection is modelled for each camera trap at each session.

Model 2: DToccu NULL model

```
dt_occu_null_code <- nimble::nimbleCode({
  # Priors
  ## Occupancy parameters
  occu_int ~ dnorm(0, sd = 1.25)

  ## Detection parameters
  det_int ~ dnorm(0, sd = 1)

  # Hierarchical occupancy model
  for (i in 1:nb_sites) {
    ## Occupancy submodel (per site)
    logit(psi[i]) <- occu_int
    z[i] ~ dbern(psi[i])

    ## Detection submodel (per camtrap and session)
    for (ct in 1:nb_cams[i]) {
      for (j in 1:nb_sessions) {
        log(mu[ct, j, i]) <- det_int
        nb_detecs[ct, j, i] ~ dpois(lambda = (
          mu[ct, j, i] *
            ndays_monitored[ct, j, i] *
            z[i]
        ))
      }
    }
  }
})
```

We define uninformative priors for both the occupancy intercept (`occu_int`) and the detection intercept (`det_int`).

**Occupancy submodel.** The occupancy state $z_i \sim \text{Bernoulli}(\psi_i)$ depends on a site-specific occupancy probability $\psi_i$, transformed from the logit scale. This format was chosen to be homogeneous with the model with covariates, but because $\psi$ does not vary across sites when no covariates are included, this could be simplified to:

```
logit(psi) <- occu_int
z[i] ~dbern(psi)
```

**Detection submodel.** Detection follows a Poisson distribution. This formulation assumes independent Poisson detection events, conditional on occupancy, and no covariates. The number of detections (discrete count data, `nb_detecs`) depends on:

- the species daily detection rate $\mu$ (on a log scale),
- the number of days monitored by this camera in this site during this session,
- whether the species was present at that site (via multiplication by $z[i]$): at least one detection implies that the site is assumed occupied.

**Required data.** The model expects the following data structures. This includes a dummy example with 3 sites and 4 sessions.

- `nb_sites`: Integer. Number of sites. For example, 3.

- `nb_sessions`: Integer. Number of sites. For example, 4.
- `nb_cams[i]`: An integers vector. Specifies the number of camera traps deployed per site (`i`). In the example, site 1 has two camera traps, while sites 2 and 3 each have one.

```
nb_cams = c('site1' = 2, 'site2' = 1, 'site3' = 1)
```

- `nb_detecs[ct, j, i]`: An integers 3D array. Number of detections per camera trap (`ct`), session (`j`), and site (`i`). If a camera trap was not active during a session (e.g. site2, cam1, session3), it can contain `NA`. In the example below, each row corresponds to a camera trap. Since sites 2 and 3 each have only one trap, the second row (`cam2`) contains only `NA`.

```
nb_detecs[ , , site1]
#           session1 session2 session3 session4
# cam1          0        1        2        0
# cam2         NA       NA        3        1

nb_detecs[ , , site2]
#           session1 session2 session3 session4
# cam1          4        1       0        0
# cam2         NA       NA       NA       NA

nb_detecs[ , , site3]
#           session1 session2 session3 session4
# cam1          0        0        0        0
# cam2         NA       NA       NA       NA
```

- `ndays_monitored[ct, j, i]`: A doubles 3D array. Number of days each camera was active for each site-session combination. In the example below, each session has a nominal duration of 7 days, though some sessions were incomplete. *Note: we used the number of days since we chose the time unit for all rates to be daily. However, this can be changed to any other time unit.*

```
ndays_monitored[ , , site1]
#           session1 session2 session3 session4
# cam1        6.5        7        7        7
# cam2          0        0        5.5      7

ndays_monitored[ , , site2]
#           session1 session2 session3 session4
# cam1          7       7       7       6.6
# cam2         NA       NA       NA      NA

ndays_monitored[ , , site3]
#           session1 session2 session3 session4
# cam1        6.9        7        7        7
# cam2         NA       NA       NA       NA
```

#### S1.3 Discrete-time occupancy model with covariates

We extend the previous NULL model to include covariates (Model 3). Occupancy is still modelled at the site level, and detection is modelled for each camera trap per session. We now include covariates to explain variation in both occupancy and detection processes.

Model 3: DToccu model with covariates

```
dt_occu_code <- nimble::nimbleCode({
  # Priors
  ## Occupancy parameters
  occu_int ~ dnorm(0, sd = 1.25)
  for (ks in 1:ncov_state) {
    occu_beta[ks] ~ dnorm(0, sd = 1)
  }

  ## Detection parameters
  det_int ~ dnorm(0, sd = 1)
  for (ka in 1:ncov_anthro) {
    det_a_beta[ka] ~ dnorm(0, sd = 1)
  }
  for (kw in 1:ncov_weather) {
    det_w_beta[kw] ~ dnorm(0, sd = 1)
  }

  for (i in 1:nb_sites) {
    ## Occupancy submodel (per site)
    logit(psi[i]) <- occu_int + sum(
      occu_beta[1:ncov_state] * state_site[i, 1:ncov_state]
    )
    z[i] ~ dbern(psi[i])

    ## Detection submodel (per camtrap and session)
    for (ct in 1:nb_cams[i]) {
      for (j in 1:nb_sessions) {
        log(mu[ct, j, i]) <- det_int +
          sum(
            det_a_beta[1:ncov_anthro] * anthro_discr[ct, j, i, 1:ncov_anthro]
          ) +
          sum(
            det_w_beta[1:ncov_weather] * weather_discr[1:ncov_weather, j]
          )
        nb_detecs[ct, j, i] ~ dpois(lambda = (
          mu[ct, j, i] *
            ndays_monitored[ct, j, i] *
            z[i]
        ))
      }
    }
  }
})
```

Occupancy probability $\psi_i$ depends on `ks` site-level covariates (`state_site`), while detection rate $\mu_{ctji}$ is modelled using two types of discrete-time covariates: `ka` anthropogenic disturbance covariates (`anthro_discr`) and `kw` weather covariates (`weather_discr`). These two sets of covariates are modelled separately because of structural differences: anthropogenic variables vary in both space and time (one value per camera per session), while weather variables vary only in time (same value for all cameras in a session). Uninformative priors are defined for all regression coefficients, including intercepts (`occu_int`, `det_int`) and slopes (`occu_beta[ks]`, `det_a_beta[ka]`, `det_w_beta[kw]`).

**Occupancy submodel.** Occupancy is modelled at the site level. The latent occupancy status $z_i$ is a Bernoulli random variable. The occupancy probability $\psi_i$ in site $i$ is linked to site-level covariates using a logit link function. Here, $x_{i,k}$ denotes the $k$th covariate value at site $i$. In our case study, these covariates originate from habitat typology, extracted from a land-use map, and summarised using PCA. The PCA scores are centred and scaled.

$$z_i \sim \text{Bernoulli}(\psi_i)$$
$$\text{logit}(\psi_i) = \beta_0 + \sum_{k=1}^{ks} \beta_k \cdot x_{i,k}$$

**Detection submodel.** Detection is modelled at the level of each camera trap ($ct$) and session ($j$) within a site ($i$). Detection counts are linked to covariates through a log-linear model. Two types of covariates are supported: spatiotemporal covariates $a_{ct,j,i,k}$ that vary by camera, session, and site, and temporal covariates $w_{jk}$ that vary by session only. This structure allows the model to accommodate covariates measured at different resolutions. In all cases, time-varying covariates influencing detectability are summarised over the session (e.g., by the mean or standard deviation).

Detection rate $\mu_{ct,j,i}$ is scaled by the number of monitoring days and multiplied by the occupancy indicator $z_i$, so that detections only occur at occupied sites. In our case study, spatiotemporal covariates represent anthropogenic disturbance (based on human detections per camera trap), and temporal covariates represent weather conditions (based on weather data collected at a unique weather station for the whole study area). These are used illustratively, but the model can incorporate any covariates following the same structural format.

$$\log(\mu_{ct,j,i}) = \alpha_0 + \sum_{k=1}^{ka} \alpha_k \cdot a_{ct,j,i,k} + \sum_{k=1}^{kw} \gamma_k \cdot w_{j,k}$$
$$y_{ct,j,i} \sim \text{Poisson}\left(\mu_{ct,j,i} \cdot \text{ndays}_{ct,j,i} \cdot z_i\right)$$

**Required data.** The model requires the following input data. The structure of `nb_sites`, `nb_sessions`, `nb_cams`, `nb_detecs`, and `ndays_monitored` is identical to that used in DToccu NULL model (see Section S1.2). Additional input is needed for covariates. These are structured according to the level at which the covariates vary: spatial (site), spatiotemporal (camera × session × site), or temporal (session). The specific covariates described here are illustrative. They are those from our case study.

- `ncov_state`, `ncov_anthro`, `ncov_weather`: Integers. Number of covariates for each covariate type.
- `state_site[i, k]`: A doubles 2D array (matrix). Static covariates at the site level (`i`), with one row per site and one column per covariate.

```
state_site
#           occu_cov1   occu_cov2
# site1          1.07        0.12
# site2         -0.89       -1.05
# site3         -0.18        0.93
```

- `anthro_discr[ct, j, i, k]`: A doubles 4D array. Spatiotemporal covariates that vary across cameras (`ct`), sessions (`j`), and sites (`i`). The fourth dimension (`k`) indexes covariates. In our case study, this includes the first two principal components of our anthropic disturbance PCA, based on human detections from camera traps (centred and scaled).
- `weather_discr[k, j]`: A doubles 2D array (or matrix). Temporal covariates that vary across sessions (`j`) but not spatially. The first dimension (`k`) indexes the covariates. In our case study, this includes the first four principal components from a PCA of weather data collected from a single weather station (centred and scaled).

#### S1.4 Continuous-time occupancy model without covariates

Model 4 gives the code used for our continuous-time occupancy NULL model, without covariates. Occupancy status is modelled at the site level, while detection is modelled for each camera trap and session.

Model 4: CToccu NULL model

```
ct_occu_null_code <- nimble::nimbleCode({
  # Priors
  ## Occupancy parameters
  occu_int ~ dnorm(0, sd = 1.25)

  ## Detection parameters
  det_int ~ dnorm(0, sd = 1)

  # Hierarchical occupancy model
  for (i in 1:nb_sites) {
    ## Occupancy submodel
    logit(psi[i]) <- occu_int
    z[i] ~ dbern(psi[i])

    ## Detection submodel per camtrap
    for (ct in 1:nb_cams[i]) {
      # Detection rate per timeint
      for (ti in 1:nb_monitored[ct, i]) {
        log(mu_by_timeint[ct, ti, i]) <- det_int
      }

      # Detection rate integral during monitoring with a Riemann sum
      mu_integral[ct, i] <- sum(mu_by_timeint[ct, 1:nb_monitored[ct, i], i]) *
        tempprec / (60 * 60 * 24)

      # Total number of detections
      nb_detec[ct, i] ~ dpois(lambda = mu_integral[ct, i] * z[i])

      for (td in 1:cste_nb_detec[ct, i]) {
        # Detection rate at the time of each detection
        log(mu_detecs[ct, td, i]) <- det_int

        # Proba of having those detections at those times (with a 1-trick)
        detecs_1[ct, td, i] ~ dbern(
          prob = mu_detecs[ct, td, i] / mu_integral[ct, i]
        )
      }
    }
  }
})
```

**Occupancy submodel.** The occupancy submodel remains exactly the same as for the NULL DToccu model (see Section S1.2).

**Detection submodel.** Detection follows a NHPP, as described in Section S1.1. However, because no covariates are included, this is equivalent to a homogeneous Poisson process and could therefore be simplified. We kept this formulation to maintain consistency between models with and without covariates.

**Required data.** The model expects the following data structures. This includes a dummy example with 3 sites, each monitored with one or two camera traps. Detection times are modelled as a continuous process, and data are structured to reflect this.

- `nb_sites`: Integer. Number of sites. For example, 3.

- `tempprec`: Integer. Temporal resolution of the discretised time intervals (in seconds). For example, `tempprec = 180` for 3-minutes time intervals.
- `nb_cams[i]`: Integer vector. Number of camera traps at each site `i`. For example:

```
nb_cams = c('site1' = 2, 'site2' = 1, 'site3' = 1)
```

- `nb_detec[ct, i]`: Integer matrix. Total number of detections per camera trap `ct` at site `i`. Following on from our previous example in Section S1.2, with 2 cameras at site 1 and 1 camera at sites 2 and 3, this would give:

```
nb_detec
#          site1    site2   site3
# cam1       3        5        0
# cam2       4        NA       NA
```

- `cste_nb_detec[ct, i]`: Same dimensions and values as `nb_detec[ct, i]`, but required both as data and as a constant in `Nimble`. `nb_detec[ct, i]` is treated as data because it is a stochastic node in the model (`nb_detec[ct, i] ~ dpois(...)`). In contrast, `cste_nb_detec[ct, i]` is treated as a constant to allow iteration in `Nimble` loops (`for (td in 1:cste_nb_detec[ct, i]) { ... }`).
- `detecs_1[ct, td, i]`: Integer 3D array of dimension `max(nb_cams)` by `max_nb_detec` by `nb_sites`. It contains only values equal to 1. Used in the ones trick to assign probability to detection times under the NHPP.
- `nb_monitored[ct, i]`: Integer matrix of dimension `max(nb_cams)` by `nb_sites`. For each camera trap `ct` at site `i`, it gives the number of time intervals used to approximate the detection rate integral with the Riemann sum. These intervals are always indexed starting from 1, representing the first monitored time interval for that specific camera and site.

  Following on our example, if daily intervals are used, and camera 1 at site 1 monitored from 1 January 2025 to 27 January 2025, `nb_monitored[1, 1]` will be 27, as 27 days are monitored by camera 1 in site 1. If camera 2 at site 1 started later, on 16 January 2025, `nb_monitored[2, 1]` will be 13.

```
nb_monitored
#          site1    site2   site3
# cam1       27       28      28
# cam2       13       NA      NA
```

  To avoid confusion and to keep track of the matching between time interval indexes and actual times, we created an additional object. Although not used directly in the model, `monitored_timeints[ct, ti, i]` is an integer 3D array with dimensions `max(nb_cams)` by `max_nb_timeint` by `nb_sites`. This array stores the actual time interval (Unix epoch timestamps) for each camera trap `ct`, time interval `ti`, and site `i`. It allows precise mapping of each time interval to calendar time, accounting for staggered start times across cameras and sites. To understand why we formatted this way, see the section on `CToccu` with covariates (Section S1.5), where it is necessary to match covariate times to each time interval.

#### S1.5 Continuous-time occupancy model with covariates

Model 5: CToccu model

```
ct_occu_code <- nimble::nimbleCode({
  # Priors
  ## Occupancy parameters
  occu_int ~ dnorm(0, sd = 1.25)
  for (ks in 1:ncov_state) {
    occu_beta[ks] ~ dnorm(0, sd = 1)
  }

  ## Detection parameters
  det_int ~ dnorm(0, sd = 1)
  for (ka in 1:ncov_anthro) {
    det_a_beta[ka] ~ dnorm(0, sd = 1)
  }
  for (kw in 1:ncov_weather) {
    det_w_beta[kw] ~ dnorm(0, sd = 1)
  }

  # Hierarchical occupancy model
  for (i in 1:nb_sites) {
    ## Occupancy submodel
    logit(psi[i]) <- occu_int + sum(
      occu_beta[1:ncov_state] * state_site[i, 1:ncov_state]
    )
    z[i] ~ dbern(psi[i])

    ## Detection submodel per camtrap
    for (ct in 1:nb_cams[i]) {
      # Detection rate per timeint
      for (ti in 1:nb_monitored[ct, i]) {
        log(mu_by_timeint[ct, ti, i]) <- det_int +
          sum(det_a_beta[1:ncov_anthro] *
                anthro_by_timeint[ct, ti, i, 1:ncov_anthro]) +
          sum(det_w_beta[1:ncov_weather] *
                weather_by_timeint[ct, ti, i, 1:ncov_weather])
      }

      # Detection rate integral during monitoring with a Riemann sum
      mu_integral[ct, i] <- sum(mu_by_timeint[ct, 1:nb_monitored[ct, i], i]) *
        tempprec / (60 * 60 * 24)

      # Total number of detections
      nb_detec[ct, i] ~ dpois(lambda = mu_integral[ct, i] * z[i])

      for (td in 1:cste_nb_detec[ct, i]) {
        # Detection rate at the time of each detection
        log(mu_detecs[ct, td, i]) <- det_int +
          sum(det_a_beta[1:ncov_anthro] *
                anthro_at_detecs[ct, td, i, 1:ncov_anthro]) +
          sum(det_w_beta[1:ncov_weather] *
                weather_at_detecs[ct, td, i, 1:ncov_weather])

        # Proba of having those detections at those times (with a 1-trick)
        detecs_1[ct, td, i] ~ dbern(
          prob = mu_detecs[ct, td, i] / mu_integral[ct, i]
        )
      }
    }
  }
})
```

**Occupancy submodel.** The occupancy submodel remains exactly the same as for the DToccu model with covariates (see Section S1.3).

**Detection submodel.** Detection follows a NHPP, as described in Section S1.1.

**Required data.** The model requires the following input data. The structure of `nb_sites`, `tempprec`, `nb_cams`, `nb_detecs`, `cste_nb_detecs`, `detecs_1`, and `nb_monitored` is identical to that used in the CToccu NULL model (see Section S1.4). Additional input is needed for covariates. These are structured according to the level at which the covariates vary: spatial, spatiotemporal, or temporal. The specific covariates described here are illustrative. They are those from our case study.

- `ncov_state`, `ncov_anthro`, `ncov_weather`: Integers. Number of covariates for each covariate type.
- `state_site[i, k]`: A doubles 2D array (matrix). This is the same as for the DToccu model with covariates (Section S1.3). Static covariates at the site level (`i`), with one row per site and one column per covariate.

```
state_site
#             occu_cov1    occu_cov2
# site1            1.07         0.12
# site2           -0.89        -1.05
# site3           -0.18         0.93
```

- `anthro_by_timeint[ct, ti, i, k]` and `weather_by_timeint[ct, ti, i, k]`: 4D arrays of doubles. Temporal or spatiotemporal detection covariates at each time interval (in our use case, describing human disturbance or weather). Each entry represents the value of covariate `k` at time interval `ti` for camera trap `ct` at site `i`.

  These arrays must be filled only for the monitored time intervals, as specified by `nb_monitored[ct, i]` (see Section S1.4 for the CToccu model without covariates). That is, for each camera `ct` at site `i`, only the first `nb_monitored[ct, i]` time intervals `ti` are meaningful and used in the model. Covariate values outside this range are ignored.

  To ensure that each covariate value is associated with the correct time, the entries must align exactly with the true time intervals. This is optional in theory, but we chose to store this information explicitly in a matching array: `monitored_timeints[ct, ti, i]` (also described in Section S1.4). This object stores the actual index (e.g., day number or timestamp) corresponding to each `ti`. It allows alignment of each covariate value to the correct point in time, accounting for staggered monitoring periods across cameras and sites.

  This matching is essential: if the covariate arrays do not align with the correct time intervals, the model will associate covariate effects with the wrong detection times. This leads to incorrect estimation of detection parameters and invalid inference about the influence of covariates on detectability.

- `anthro_at_detecs[ct, td, i, k]` and `weather_at_detecs[ct, td, i, k]`: 4D arrays of doubles. Temporal or spatiotemporal detection covariates at each detection time (in our use case, describing human disturbance or weather). Each entry represents the value of covariate `k` at detection `td` for camera trap `ct` at site `i`.

  Once again, there is a matching process to align covariate values with the actual detection times. To facilitate this, we created a matching array, `time_at_detecs[ct, td, i]`, which stores the actual time (Unix epoch timestamps) of each detection `td` for camera `ct` at site `i`. This object is not used directly in the model but is essential for preparing the covariate arrays. It ensures that each detection is associated with the correct covariate value. This is particularly important since cameras produce variable numbers of detections.

  However, covariate values are not always available exactly at the detection times. Here, our interpolation method comes into play. The covariates by time interval (as described above) are used to create interpolation functions, which are then evaluated at the detection times stored in `time_at_detecs`. This produces the interpolated covariate values used to fill `anthro_at_detecs[ct, td, i, k]`, ensuring precise alignment of covariates with each detection event.

#### S1.6 N-mixture models

The process will not be detailed separately for each model (DTabun vs CTabun, with and without covariates), as much of it is common to both occupancy and N-mixture models. The main differences are:

- In the **state submodel**: Instead of modelling site occupancy as a Bernoulli variable, we model site abundance as a Poisson-distributed count.
- In the **detection submodel**: The detection rate is no longer $\mu(t)$, the species detection rate, but $\lambda(t)$, the individual detection rate. To model the number of detections, we multiply the detection rate (of the session for DTabun, or its integral over deployment for CTabun) by `N[i]`, the number of individuals at site $i$ (0, 1, 2, ...), instead of by `z[i]`, the occupancy state (0/1).

To illustrate these changes, the N-mixture model codes are provided below:

- Model 6: DTabun NULL model, without covariates
- Model 7: DTabun model with covariates
- Model 8: CTabun NULL model, without covariates
- Model 9: CTabun model with covariates

Model 6: DTabun NULL model

```
dt_abun_null_code <- nimble::nimbleCode({
  # Priors
  ## abundance parameters
  abun_int ~ dnorm(0, sd = 1.25)

  ## Detection parameters
  det_int ~ dnorm(0, sd = 1)

  # Hierarchical N-mixture model with a detection process per camtrap, not site
  for (i in 1:nb_sites) {
    ## Abundance submodel (per site)
    log(phi[i]) <- abun_int
    N[i] ~ dpois(phi[i])

    ## Detection submodel (per camtrap and session)
    for (ct in 1:nb_cams[i]) {
      for (j in 1:nb_sessions) {
        log(lambda[ct, j, i]) <- det_int
        nb_detecs[ct, j, i] ~ dpois(lambda = (
          lambda[ct, j, i] *
            ndays_monitored[ct, j, i] *
            N[i]
        ))
      }
    }
  }
})
```

Model 7: DTabun model with covariates

```
dt_abun_code <- nimble::nimbleCode({
  # Priors
  ## abundance parameters
  abun_int ~ dnorm(0, sd = 1.25)
  for (ks in 1:ncov_state) {
    abun_beta[ks] ~ dnorm(0, sd = 1)
  }

  ## Detection parameters
  det_int ~ dnorm(0, sd = 1)
  for (ka in 1:ncov_anthro) {
    det_a_beta[ka] ~ dnorm(0, sd = 1)
  }
  for (kw in 1:ncov_weather) {
    det_w_beta[kw] ~ dnorm(0, sd = 1)
  }

  # Hierarchical N-mixture model with a detection process per camtrap, not site
  for (i in 1:nb_sites) {
    ## Abundance submodel (per site)
    log(phi[i]) <- abun_int + sum(
      abun_beta[1:ncov_state] * state_cams[i, 1:ncov_state]
    )
    N[i] ~ dpois(phi[i])

    ## Detection submodel (per camtrap and session)
    for (ct in 1:nb_cams[i]) {
      for (j in 1:nb_sessions) {
        log(lambda[ct, j, i]) <- det_int +
          sum(
            det_a_beta[1:ncov_anthro] * anthro_discr[ct, j, i, 1:ncov_anthro]
          ) +
          sum(
            det_w_beta[1:ncov_weather] * weather_discr[1:ncov_weather, j]
          )
        nb_detecs[ct, j, i] ~ dpois(lambda = (
          lambda[ct, j, i] *
            ndays_monitored[ct, j, i] *
            N[i]
        ))
      }
    }
  }
})
```

Model 8: CTabun NULL model

```
ct_abun_null_code <- nimble::nimbleCode({
  # Priors
  ## Abundance parameters
  abun_int ~ dnorm(0, sd = 1.25)

  ## Detection parameters
  det_int ~ dnorm(0, sd = 1)

  # Hierarchical abunpancy model
  for (i in 1:nb_sites) {
    ## Abundance submodel (per site)
    log(phi[i]) <- abun_int
    N[i] ~ dpois(phi[i])

    ## Detection submodel per camtrap
    for (ct in 1:nb_cams[i]) {
      # Detection rate per timeint
      for (ti in 1:nb_monitored[ct, i]) {
        log(lambda_by_timeint[ct, ti, i]) <- det_int
      }

      # Detection rate integral during monitoring with a Riemann sum
      lambda_integral[ct, i] <-
        sum(lambda_by_timeint[ct, 1:nb_monitored[ct, i], i]) *
            tempprec / (60 * 60 * 24)

      # Total number of detections
      nb_detec[ct, i] ~ dpois(lambda = lambda_integral[ct, i] * N[i])

      for (td in 1:cste_nb_detec[ct, i]) {
        # Detection rate at the time of each detection
        log(lambda_detecs[ct, td, i]) <- det_int

        # Proba of having those detections at those times (with a 1-trick)
        detecs_1[ct, td, i] ~ dbern(
          prob = lambda_detecs[ct, td, i] / lambda_integral[ct, i]
        )
      }
    }
  }
})
```

Model 9: CTabun model

```
ct_abun_code <- nimble::nimbleCode({
  # Priors
  ## Abundance parameters
  abun_int ~ dnorm(0, sd = 1.25)
  for (ks in 1:ncov_state) {
    abun_beta[ks] ~ dnorm(0, sd = 1)
  }

  ## Detection parameters
  det_int ~ dnorm(0, sd = 1)
  for (ka in 1:ncov_anthro) {
    det_a_beta[ka] ~ dnorm(0, sd = 1)
  }
  for (kw in 1:ncov_weather) {
    det_w_beta[kw] ~ dnorm(0, sd = 1)
  }

  # Hierarchical abunpancy model
  for (i in 1:nb_sites) {
    ## Abundance submodel (per site)
    log(phi[i]) <- abun_int + sum(
      abun_beta[1:ncov_state] * state_cams[i, 1:ncov_state]
    )
    N[i] ~ dpois(phi[i])

    ## Detection submodel per camtrap
    for (ct in 1:nb_cams[i]) {
      # Detection rate per timeint
      for (ti in 1:nb_monitored[ct, i]) {
        log(lambda_by_timeint[ct, ti, i]) <- det_int +
          sum(det_a_beta[1:ncov_anthro] *
                anthro_by_timeint[ct, ti, i, 1:ncov_anthro]) +
          sum(det_w_beta[1:ncov_weather] *
                weather_by_timeint[ct, ti, i, 1:ncov_weather])
      }

      # Detection rate integral during monitoring with a Riemann sum
      lambda_integral[ct, i] <-
        sum(lambda_by_timeint[ct, 1:nb_monitored[ct, i], i]) *
          tempprec / (60 * 60 * 24)

      # Total number of detections
      nb_detec[ct, i] ~ dpois(lambda = lambda_integral[ct, i] * N[i])

      for (td in 1:cste_nb_detec[ct, i]) {
        # Detection rate at the time of each detection
        log(lambda_detecs[ct, td, i]) <- det_int +
          sum(det_a_beta[1:ncov_anthro] *
                anthro_at_detecs[ct, td, i, 1:ncov_anthro]) +
          sum(det_w_beta[1:ncov_weather] *
                weather_at_detecs[ct, td, i, 1:ncov_weather])

        # Proba of having those detections at those times (with a 1-trick)
        detecs_1[ct, td, i] ~ dbern(
          prob = lambda_detecs[ct, td, i] / lambda_integral[ct, i]
        )
      }
    }
  }
})
```
