## Appendix 2 for "When does temporal resolution matter? Including detection covariates in discrete- versus continuous-time occupancy and N-mixture models"

### Appendix S2: supplementary figures
#### When does temporal resolution matter? Including detection covariates in discrete- versus continuous-time occupancy and N-mixture models

Léa Pautrel[1, 2], Sylvain Moulherat[2], Benoit Charrasse[3], Guillaume Debat[2], Lucie Gendron[2], Kenneth Kellner[4], Marie-Pierre Etienne[5, *], and Olivier Gimenez[1, *]

[1]Centre d'Ecologie Fonctionnelle et Evolutive (CEFE), University of Montpellier, CNRS, EPHE, IRD, 1919 Route de Mende, 34000 Montpellier, France
[2]OïkoLab, TerrOïko, 2 Place Dom Devic, BP 26, 81540 Sorèze, France
[3]CEA, DES, IRESNE, DTN, Laboratory for Environmental Transfer Modeling, Cadarache, 13108 Saint-Paul-lès-Durance, France
[4]Department of Fisheries and Wildlife, Michigan State University, 480 Wilson Rd, East Lansing, Michigan, USA
[5]Univ Rennes, Ensai, CNRS, CREST – UMR 9194, F-35000 Rennes, France
[*]Co-senior authors

01 September 2025

### List of Tables

S1 Image-level AI classification performance metrics per taxon . . . 8
S2 Sequence-level AI classification performance metrics by taxon . . . 8

#### S2.1 Simulation study details

To fit models, we estimated initial values based on simplified summaries of the simulated data. We first discretised the time-varying data to daily resolution: we summed detections per day and calculated the daily mean for each detection covariate.

- **For occupancy models**, we estimated the initial occupancy probability as the proportion of sites (among the 100 for which we simulated both occupancy and detection processes) with at least one detection across the full deployment period. This proportion was then logit-transformed. To estimate the detection rate parameters (intercept and three covariate effects), we subsetted to sites with at least one detection and fitted a generalised linear model (GLM) with Poisson distribution and log link. The response variable was the total number of daily detections per site, and the predictors were the daily means of the three covariates (`covday`, `covyear`, `covarma`).
- **For N-mixture models**, we estimated the initial mean abundance by dividing the mean number of detections per site by the expected number of detections per individual (a known simulation parameter). We used this parameter because we assumed that modellers would have some prior knowledge about the detectability of the studied species when choosing initial values. A log transformation was then applied. For the detection rate parameters, we used all sites and again fitted a Poisson GLM with log link, using total daily detections as the response and the daily mean values of the three covariates as predictors.

To assess model inference quality, we computed four complementary metrics by comparing estimated parameters to their true simulated values:

- **Root mean squared error (RMSE)**: Figure in the main text
- **Bias**: Figure S1
- **Coverage** of the 95% credible interval: Figure S2
- **Average width** of the 95% credible interval: Figure S3

All simulations were distributed across three Linux Debian 12 machines:

- ***Ringhal***: 2× AMD EPYC 9274F CPUs (48 cores, 96 threads total; up to 4.3,GHz; 512,MB L3 cache) with 515 GB RAM
- ***Crotale***: 2× AMD EPYC 7402 CPUs (48 cores, 96 threads total; up to 2.8,GHz; 256,MB L3 cache) with 515 GB RAM
- ***Cobra***: 2× Intel Xeon Gold 6154 CPUs (36 cores, 72 threads total; up to 3.7,GHz) with 385 GB RAM

Fitting durations for all model configurations across these machines are presented in Figure S4.

| | | Abundance models | | | | | | | | | | | | | Occupancy models | | | | | | | | | | | | |
|---|---|---|---|---|---|---|---|---|---|---|---|---|---|---|---|---|---|---|---|---|---|---|---|---|---|---|---|
| | | CTabun | | | | | DTabun | | | | | | | | CToccu | | | | | DToccu | | | | | | | |
| | | Gaussian interpolation | | Linear interpolation | | | 3 hours sessions | | Daily sessions | | | Weekly sessions | | | Gaussian interpolation | | Linear interpolation | | | 3 hours sessions | | Daily sessions | | | Weekly sessions | | |
| | | hour | day | 3 mins | hour | day | 3 mins | hour | 3 mins | hour | day | 3 mins | hour | day | hour | day | 3 mins | hour | day | 3 mins | hour | 3 mins | hour | day | 3 mins | hour | day |
| No impact / d0_y0_a0 | det_covarma | 0.001 | 0.001 | 0.001 | 0.001 | 0.001 | 0.001 | 0.001 | 0.001 | 0.002 | 0.001 | 0.001 | 0.001 | 0.001 | -0.013 | -0.013 | -0.013 | -0.013 | -0.013 | -0.013 | -0.013 | -0.013 | -0.013 | -0.013 | -0.013 | -0.014 | -0.013 |
| | det_covyear | -0.007 | -0.003 | -0.008 | -0.008 | -0.006 | -0.008 | -0.008 | -0.008 | -0.008 | -0.007 | -0.008 | -0.008 | -0.008 | -0.005 | -0.001 | -0.005 | -0.005 | -0.004 | -0.005 | -0.005 | -0.005 | -0.005 | -0.005 | -0.005 | -0.005 | -0.005 |
| | det_covday | 0.001 | -0.048 | 0.003 | 0.003 | -0.021 | 0.002 | 0.002 | -0.019 | -0.020 | -0.021 | -0.019 | -0.019 | -0.020 | 0.002 | -0.027 | -0.005 | -0.004 | -0.002 | -0.003 | -0.004 | 0.000 | 0.000 | -0.005 | -0.004 | -0.003 | -0.005 |
| | det_int | -0.153 | -0.170 | -0.151 | -0.151 | -0.148 | -0.153 | -0.151 | -0.154 | -0.153 | -0.145 | -0.156 | -0.155 | -0.148 | -0.003 | -0.039 | -0.002 | -0.003 | -0.025 | -0.003 | -0.003 | -0.004 | -0.004 | -0.022 | -0.004 | -0.004 | -0.025 |
| | state_int | 0.154 | 0.154 | 0.154 | 0.153 | 0.154 | 0.155 | 0.154 | 0.155 | 0.153 | 0.155 | 0.156 | 0.155 | 0.155 | 0.029 | 0.029 | 0.028 | 0.028 | 0.030 | 0.029 | 0.029 | 0.029 | 0.029 | 0.030 | 0.028 | 0.029 | 0.030 |
| 1 covariate / d0_y0_a2 | det_covarma | 0.069 | 0.082 | 0.068 | 0.069 | 0.084 | 0.067 | 0.068 | 0.070 | 0.070 | 0.093 | 0.071 | 0.071 | 0.073 | 0.044 | 0.051 | 0.044 | 0.044 | 0.052 | 0.044 | 0.045 | 0.045 | 0.045 | 0.055 | 0.046 | 0.045 | 0.046 |
| | det_covyear | -0.009 | -0.002 | -0.009 | -0.009 | -0.006 | -0.009 | -0.009 | -0.008 | -0.008 | -0.008 | -0.008 | -0.008 | -0.008 | -0.006 | -0.002 | -0.006 | -0.006 | -0.005 | -0.006 | -0.006 | -0.008 | -0.008 | -0.006 | -0.008 | -0.008 | -0.007 |
| | det_covday | 0.002 | -0.034 | 0.005 | 0.005 | -0.006 | 0.005 | 0.004 | -0.003 | -0.005 | -0.004 | -0.001 | -0.001 | 0.000 | 0.007 | 0.030 | 0.004 | 0.005 | 0.061 | 0.005 | 0.005 | 0.060 | 0.061 | 0.062 | 0.065 | 0.065 | 0.068 |
| | det_int | -0.478 | -0.500 | -0.477 | -0.476 | -0.484 | -0.474 | -0.475 | -0.481 | -0.481 | -0.488 | -0.495 | -0.492 | -0.482 | -0.060 | -0.040 | -0.059 | -0.059 | -0.026 | -0.059 | -0.061 | -0.060 | -0.060 | -0.028 | -0.072 | -0.072 | -0.015 |
| | state_int | 0.427 | 0.429 | 0.426 | 0.425 | 0.433 | 0.424 | 0.424 | 0.426 | 0.426 | 0.434 | 0.429 | 0.427 | 0.430 | 0.073 | 0.076 | 0.073 | 0.072 | 0.077 | 0.073 | 0.073 | 0.073 | 0.073 | 0.078 | 0.073 | 0.073 | 0.074 |
| 1 covariate / d0_y2_a0 | det_covarma | -0.003 | -0.003 | -0.003 | -0.003 | -0.002 | -0.003 | -0.003 | -0.003 | -0.002 | -0.002 | -0.002 | -0.002 | -0.002 | 0.004 | 0.005 | 0.004 | 0.004 | 0.005 | 0.004 | 0.004 | 0.004 | 0.004 | 0.004 | 0.004 | 0.004 | 0.003 |
| | det_covyear | 0.061 | 0.359 | 0.040 | 0.041 | 0.023 | 0.040 | 0.040 | 0.031 | 0.031 | 0.033 | 0.034 | 0.034 | 0.034 | 0.040 | 0.318 | 0.018 | 0.018 | 0.012 | 0.018 | 0.018 | 0.023 | 0.023 | 0.020 | 0.023 | 0.022 | 0.023 |
| | det_covday | 0.004 | -0.046 | 0.005 | 0.005 | 0.110 | 0.004 | 0.004 | 0.050 | 0.051 | 0.080 | 0.044 | 0.049 | 0.080 | 0.008 | -0.109 | 0.001 | 0.002 | 0.112 | 0.002 | 0.001 | 0.042 | 0.043 | 0.102 | 0.054 | 0.055 | 0.095 |
| | det_int | -0.214 | -0.549 | -0.195 | -0.195 | -0.295 | -0.195 | -0.194 | -0.188 | -0.189 | -0.223 | -0.219 | -0.218 | -0.258 | -0.032 | -0.322 | -0.012 | -0.012 | -0.054 | -0.012 | -0.012 | -0.018 | -0.018 | -0.007 | -0.047 | -0.047 | -0.022 |
| | state_int | 0.165 | 0.155 | 0.165 | 0.165 | 0.165 | 0.165 | 0.164 | 0.165 | 0.166 | 0.166 | 0.164 | 0.163 | 0.163 | 0.029 | 0.029 | 0.030 | 0.029 | 0.030 | 0.029 | 0.029 | 0.030 | 0.029 | 0.029 | 0.029 | 0.029 | 0.029 |
| 1 covariate / d2_y0_a0 | det_covarma | 0.004 | 0.005 | 0.005 | 0.005 | 0.005 | 0.005 | 0.005 | 0.005 | 0.005 | 0.005 | 0.008 | 0.008 | 0.008 | 0.003 | 0.004 | 0.004 | 0.004 | 0.003 | 0.004 | 0.004 | 0.004 | 0.004 | 0.003 | 0.003 | 0.003 | 0.003 |
| | det_covyear | -0.009 | 0.001 | -0.004 | -0.003 | 0.003 | -0.003 | -0.003 | -0.002 | -0.002 | -0.004 | -0.001 | -0.001 | -0.003 | 0.011 | 0.028 | 0.016 | 0.017 | 0.021 | 0.016 | 0.017 | 0.016 | 0.016 | 0.014 | 0.015 | 0.015 | 0.014 |
| | det_covday | 1.747 | 0.235 | 0.029 | 0.056 | 0.047 | 0.088 | 0.125 | 0.058 | 0.059 | 0.052 | 0.059 | 0.060 | 0.026 | 1.752 | 0.249 | 0.072 | 0.096 | 0.063 | 0.125 | 0.159 | 0.117 | 0.118 | 0.063 | 0.120 | 0.119 | 0.043 |
| | det_int | -1.676 | -1.388 | -0.281 | -0.312 | -1.304 | -0.398 | -0.434 | -1.454 | -1.454 | -1.298 | -1.458 | -1.457 | -1.302 | -1.467 | -1.266 | -0.083 | -0.112 | -1.167 | -0.193 | -0.227 | -1.250 | -1.250 | -1.161 | -1.252 | -1.251 | -1.153 |
| | state_int | 0.206 | 0.211 | 0.247 | 0.246 | 0.214 | 0.244 | 0.242 | 0.213 | 0.212 | 0.215 | 0.214 | 0.213 | 0.212 | 0.030 | 0.029 | 0.030 | 0.029 | 0.029 | 0.029 | 0.030 | 0.029 | 0.029 | 0.030 | 0.029 | 0.029 | 0.029 |
| 3 covariates / d1_y1_a1 | det_covarma | 0.006 | 0.009 | 0.006 | 0.006 | 0.010 | 0.006 | 0.007 | 0.005 | 0.005 | 0.010 | 0.004 | 0.004 | 0.005 | 0.009 | 0.012 | 0.010 | 0.010 | 0.012 | 0.011 | 0.010 | 0.010 | 0.010 | 0.012 | 0.010 | 0.009 | 0.010 |
| | det_covyear | 0.012 | 0.119 | 0.003 | 0.003 | 0.004 | 0.003 | 0.003 | 0.001 | 0.001 | 0.000 | 0.005 | 0.005 | 0.004 | 0.034 | 0.124 | 0.025 | 0.026 | 0.023 | 0.025 | 0.025 | 0.019 | 0.019 | 0.019 | 0.021 | 0.021 | 0.021 |
| | det_covday | 0.864 | 0.114 | 0.004 | 0.012 | 0.059 | 0.013 | 0.025 | 0.049 | 0.049 | 0.055 | 0.048 | 0.048 | 0.039 | 0.862 | 0.115 | 0.003 | 0.011 | 0.075 | 0.009 | 0.022 | 0.079 | 0.081 | 0.076 | 0.073 | 0.074 | 0.054 |
| | det_int | -0.671 | -0.648 | -0.253 | -0.260 | -0.549 | -0.278 | -0.284 | -0.613 | -0.612 | -0.514 | -0.625 | -0.625 | -0.525 | -0.457 | -0.447 | -0.025 | -0.032 | -0.345 | -0.047 | -0.055 | -0.398 | -0.397 | -0.314 | -0.409 | -0.408 | -0.333 |
| | state_int | 0.233 | 0.232 | 0.248 | 0.249 | 0.237 | 0.248 | 0.247 | 0.233 | 0.233 | 0.236 | 0.234 | 0.234 | 0.235 | 0.016 | 0.016 | 0.015 | 0.015 | 0.016 | 0.015 | 0.016 | 0.015 | 0.015 | 0.016 | 0.015 | 0.014 | 0.014 |
| 3 covariates / d2_y2_a2 | det_covarma | 0.094 | 0.109 | 0.107 | 0.105 | 0.108 | 0.100 | 0.099 | 0.090 | 0.091 | 0.119 | 0.091 | 0.092 | 0.096 | 0.074 | 0.082 | 0.090 | 0.089 | 0.085 | 0.087 | 0.087 | 0.077 | 0.077 | 0.087 | 0.077 | 0.077 | 0.080 |
| | det_covyear | 0.107 | 0.401 | 0.054 | 0.056 | 0.055 | 0.053 | 0.055 | 0.043 | 0.043 | 0.045 | 0.048 | 0.048 | 0.048 | 0.150 | 0.414 | 0.108 | 0.109 | 0.112 | 0.107 | 0.108 | 0.100 | 0.100 | 0.103 | 0.103 | 0.104 | 0.109 |
| | det_covday | 1.779 | 0.360 | 0.043 | 0.070 | 0.146 | 0.103 | 0.141 | 0.127 | 0.125 | 0.106 | 0.140 | 0.141 | 0.073 | 1.778 | 0.249 | 0.078 | 0.103 | 0.153 | 0.130 | 0.169 | 0.192 | 0.198 | 0.143 | 0.206 | 0.209 | 0.060 |
| | det_int | -2.062 | -2.084 | -0.710 | -0.740 | -1.749 | -0.810 | -0.848 | -1.825 | -1.823 | -1.671 | -1.868 | -1.866 | -1.684 | -1.675 | -1.673 | -0.315 | -0.342 | -1.336 | -0.415 | -0.453 | -1.453 | -1.453 | -1.280 | -1.495 | -1.496 | -1.304 |
| | state_int | 0.468 | 0.466 | 0.538 | 0.536 | 0.483 | 0.527 | 0.527 | 0.478 | 0.476 | 0.486 | 0.475 | 0.473 | 0.478 | 0.058 | 0.059 | 0.065 | 0.065 | 0.061 | 0.064 | 0.064 | 0.057 | 0.058 | 0.063 | 0.057 | 0.058 | 0.058 |

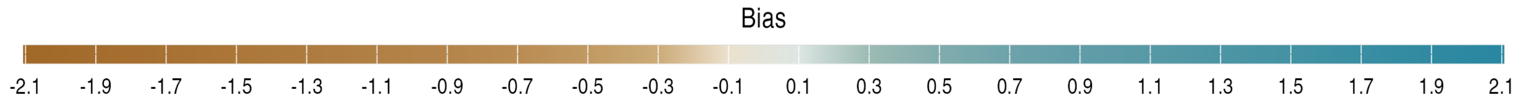

Figure S1: **Mean absolute bias for occupancy and abundance models.** Rows correspond to model terms. Columns represent model types and configurations, as indicated in the top facets: occupancy or abundance; CT or DT; interpolation method (CT) or discretisation (DT); and covariate temporal resolution (one value every 3 minutes, hour, or day). Right-hand facets show different simulated impact scenarios, as described in the manuscript. For example, scenario `d2_y0_a0` indicates a coefficient of 2 for `covday` (`d2`) on the detection rate, with no effect of `covyear` (`y0`) or `covarma` (`a0`). Cell colours and values indicate the mean absolute bias. A **positive value** (blue) means that the parameter has been **underestimated** across simulations.

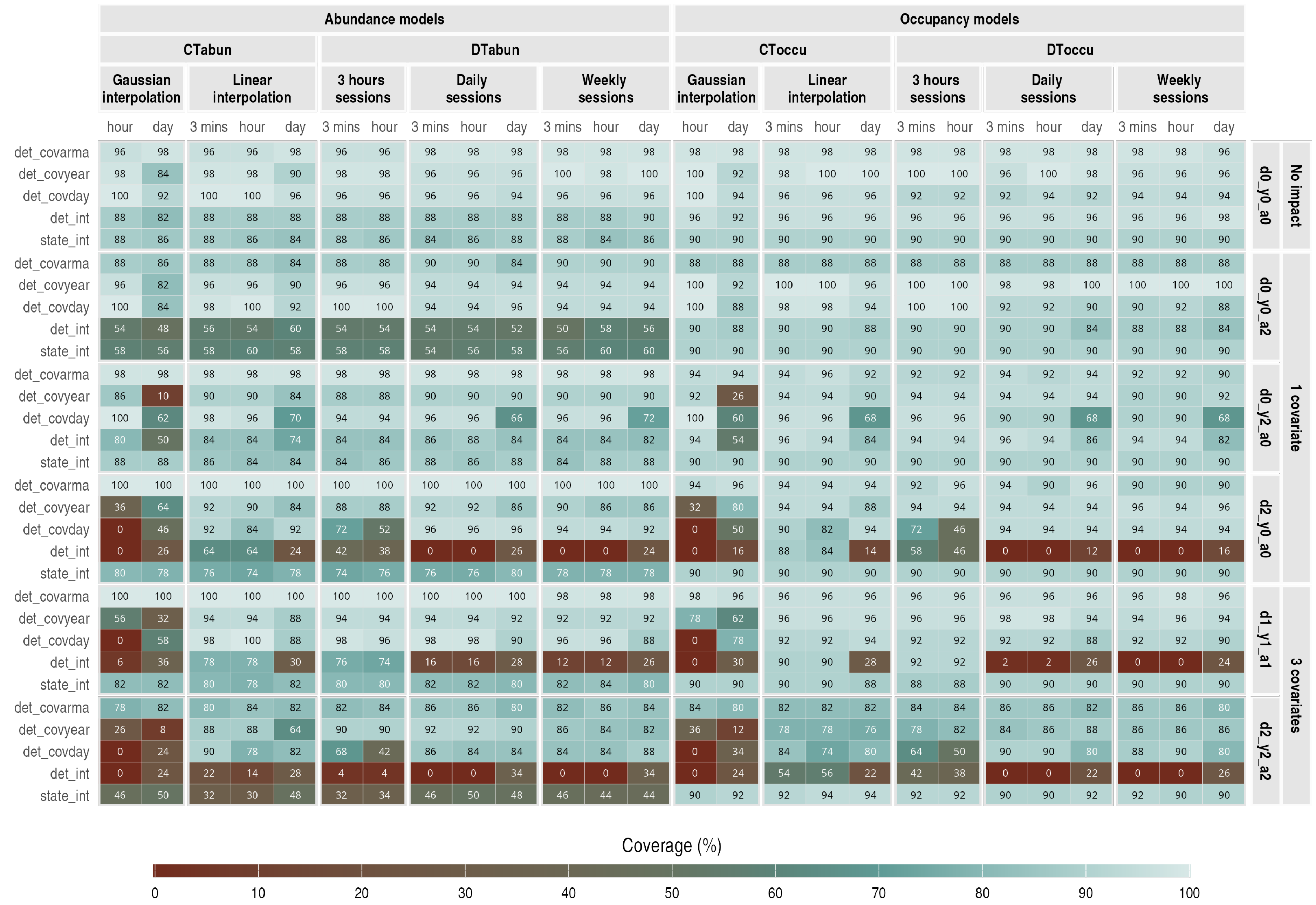

Figure S2: **Coverage for occupancy and abundance models**. Rows correspond to model terms. Columns represent model types and configurations, as indicated in the top facets: occupancy or abundance; CT or DT; interpolation method (CT) or discretisation (DT); and covariate temporal resolution (one value every 3 minutes, hour, or day). Right-hand facets show different simulated impact scenarios, as described in the manuscript. For example, scenario `d2_y0_a0` indicates a coefficient of 2 for `covday` (`d2`) on the detection rate, with no effect of `covyear` (`y0`) or `covarma` (`a0`). Cell colours and values indicate coverage, the percentage of simulations in which the simulated value was within the 95% credible interval.

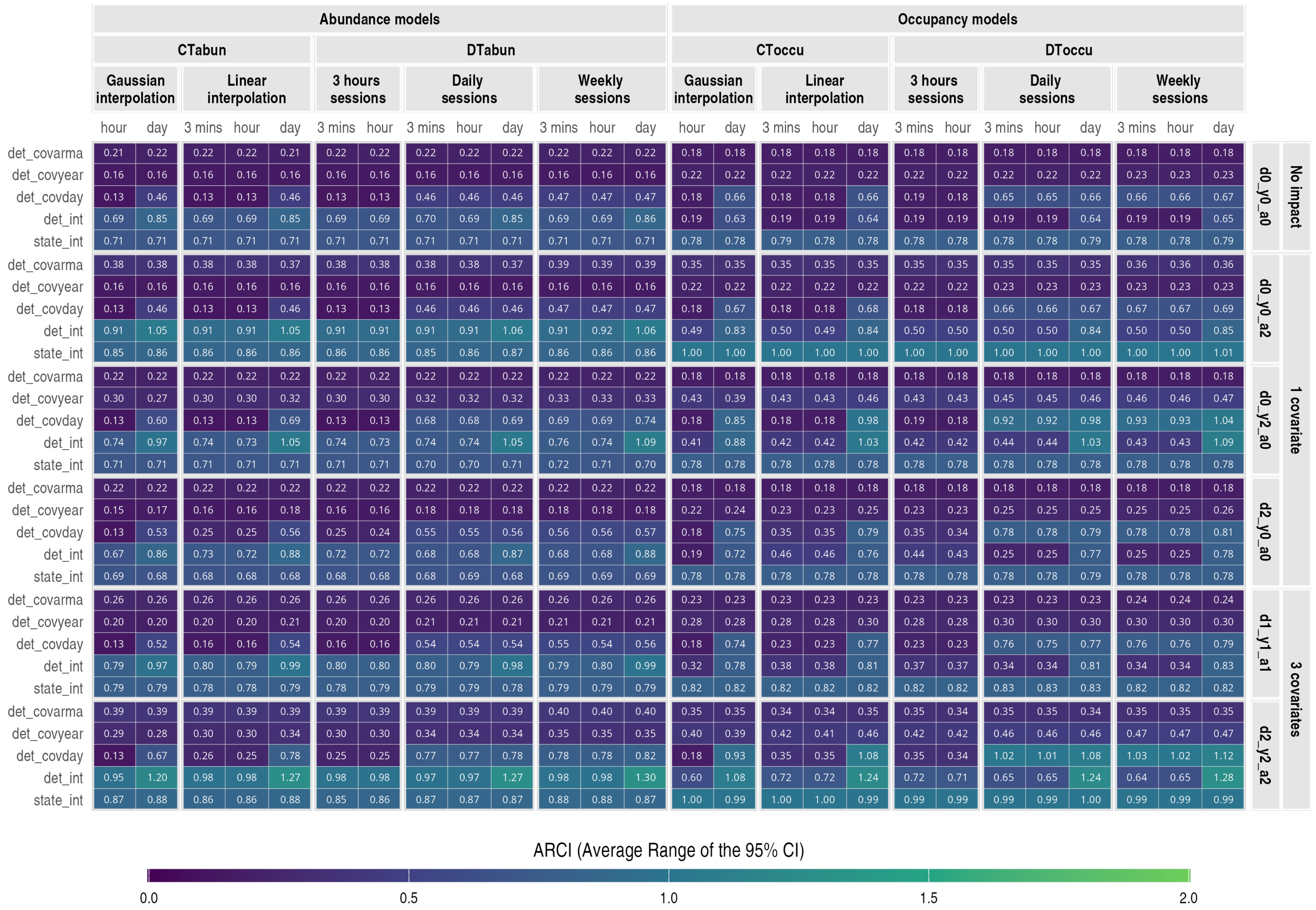

Figure S3: **Average range of the 95% credible interval for occupancy and abundance models.** Rows correspond to model terms. Columns represent model types and configurations, as indicated in the top facets: occupancy or abundance; CT or DT; interpolation method (CT) or discretisation (DT); and covariate temporal resolution (one value every 3 minutes, hour, or day). Right-hand facets show different simulated impact scenarios, as described in the manuscript. For example, scenario `d2_y0_a0` indicates a coefficient of 2 for `covday` (`d2`) on the detection rate, with no effect of `covyear` (`y0`) or `covarma` (`a0`). Cell colours and values indicate the mean width of the 95% credible interval across simulations.

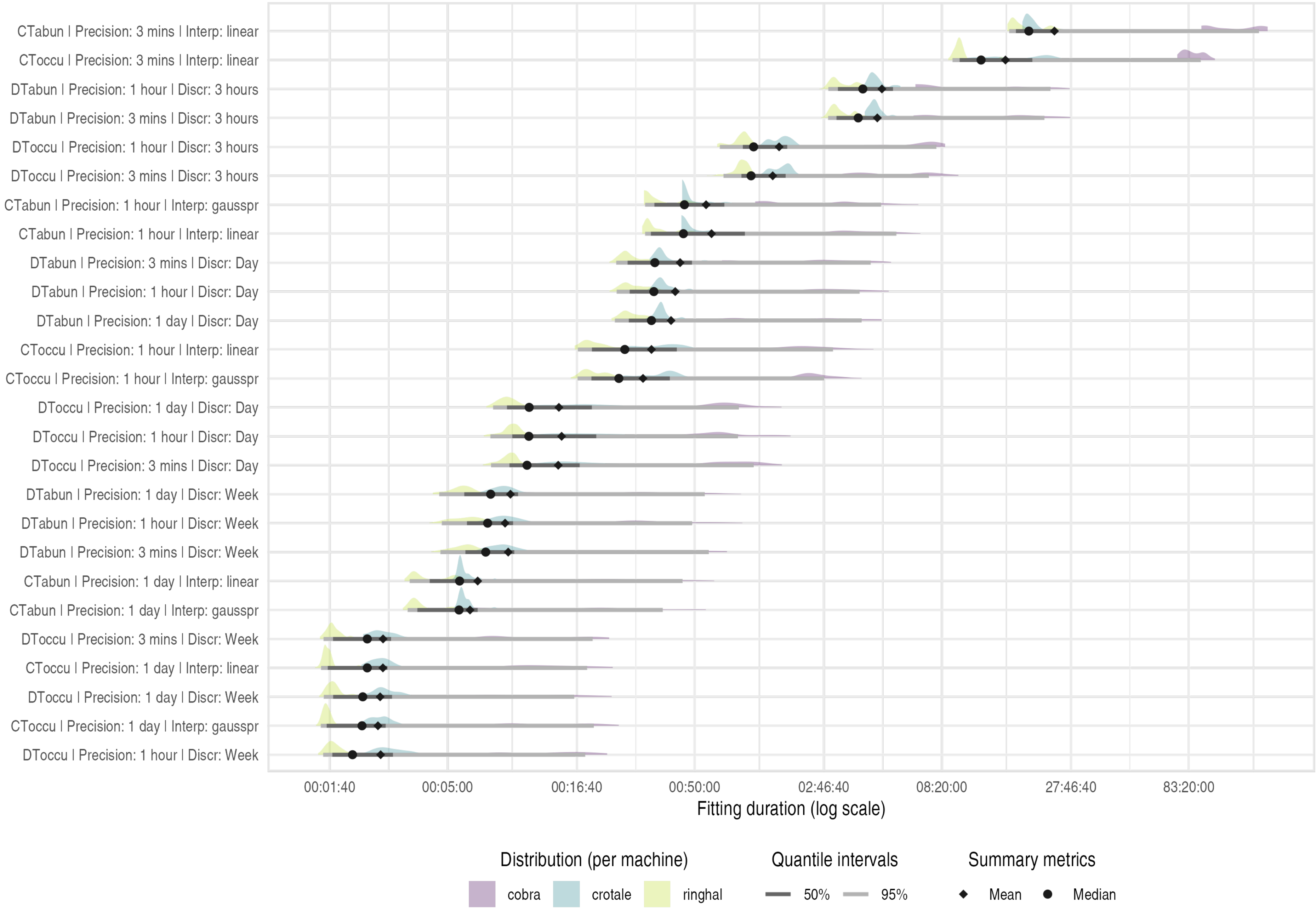

Figure S4: **Fitting duration**. Each row corresponds to a model configuration, ordered from shortest (bottom) to longest (top) runtimes. Distributions show log-transformed fitting durations (in seconds), grouped by computing machine. Horizontal lines indicate the 50% and 95% quantiles of the overall distribution (all machines combined). Points show the mean (triangle) and median (circle) durations across all machines.

#### S2.2 Use-case details

##### S2.2.1 Detection data

- Figure S5 shows the confusion matrix between AI-predicted labels and annotated labels in the subset of 9,180 annotated images.
- Table S1 shows the image-level performance of the AI for the selected taxa (wild boar, mouflon, red fox and small mustelids) and for humans, used to measure anthropic disturbance.
- Table S2 shows the sequence-level performance of the AI for those taxa.
- Figures S6, S7, S8, S9 show the detection histories of the selected taxa after filtering observations seen in all images of a given sequence.

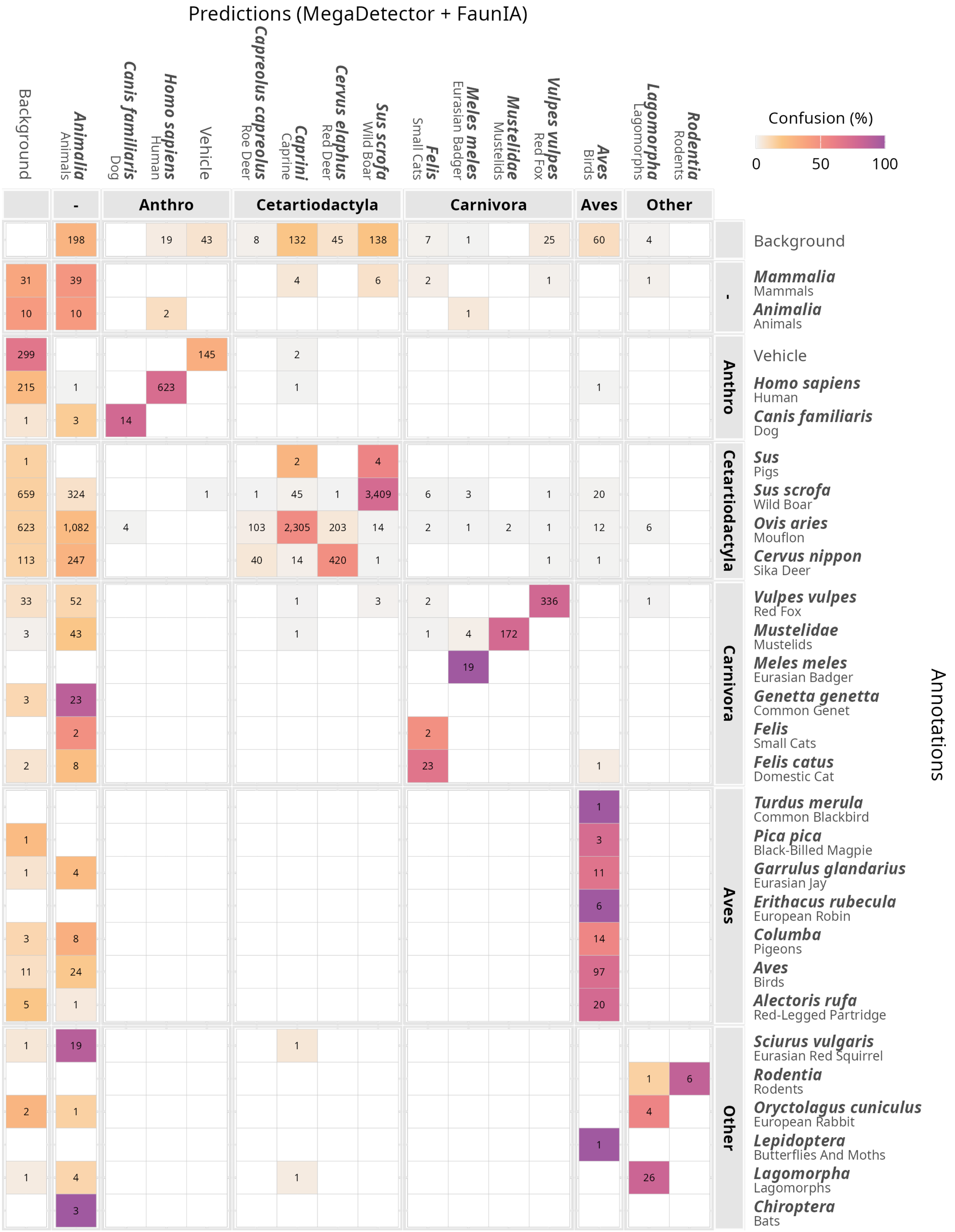

Figure S5: **Confusion matrix between annotated (right) and predicted (MegaDetetor + FaunIA, top) labels.** With 9,180 annotated images. Colour indicates the percentage of confusion relative to the number of annotations per label; numbers show the absolute number of overlapping bounding boxes. Anthro stands for anthropogenic. Some labels match despite name differences, *e.g.*, FaunIA predicts *caprini*, a tribe including goats and sheep, but since only mouflon (*Ovis aries*) occurs on site, these predictions match mouflon annotations. Annotations were made at the most precise level the annotator was confident about. For example, if unsure whether it was a wildcat (*Felis silvestris*) or a domestic cat (*Felis catus*), it was labelled as *Felis* (genus level).

Table S1: **Image-level AI classification performance metrics per taxon**. Metrics computed from 9,180 annotated images (out of 795,159 total). With **TP** = true positives, **FP** = false positives, **FN** = false negatives. **bg** = background; **unk** = unknown animal (labelled as Mammalia or Animalia in the confusion matrix, Figure S5); **other** = misclassified as another species. For example, **FP bg** = 25 for red fox means that 25 boxes were predicted as fox but actually contained no animal. **FN other** = 348 for mouflon indicates that 348 boxes were predicted as another animal (mostly roe deer: 103 boxes, and red deer: 203; see the row for mouflon's annotations in Figure S5) but were annotated as mouflon. **Annotator unsure** = 6 for wild boar means that 6 boxes were predicted as boar but were annotated as an unknown animal or mammal because the annotator was unsure, and are thus excluded from the precision and recall calculations.
*Note: Mouflon annotations were matched with caprini predictions, as FaunIA does not recognise mouflon specifically, and because mouflon is the only species from the Ovis and Capra taxa present at the site.*

| Taxon | Precision | Recall | TP | FP bg | FP other | FN bg | FN unk | FN other | Annotator unsure |
|---|---|---|---|---|---|---|---|---|---|
| Wild boar<br>*Sus scrofa* | 95.52 % | 76.26 % | 3409 | 138 | 22 | 659 | 324 | 78 | 6 |
| Mouflon<br>*Ovis aries* | 92.02 % | 52.89 % | 2305 | 132 | 68 | 623 | 1082 | 348 | 4 |
| Red fox<br>*Vulpes vulpes* | 92.31 % | 78.5 % | 336 | 25 | 3 | 33 | 52 | 7 | 1 |
| Small mustelids<br>*Martes/Mustela* spp. | 98.85 % | 76.79 % | 172 | 0 | 2 | 3 | 43 | 6 | 0 |
| Human<br>*Homo sapiens* | 97.04 % | 74.08 % | 623 | 19 | 0 | 215 | 1 | 2 | 2 |

Table S2: **Sequence-level AI classification performance metrics by taxon**. Metrics computed from 2,240 fully annotated sequences (out of 168,916 total). With **TP seq** = true positives at the sequence level (all images in the sequence are annotated and predicted with the label); **FP seq** = false positives (not all images are annotated with the label, but all are predicted with it); **FN seq** = false negatives (all images are annotated with the label, but not all are predicted with it); **TN seq** = true negatives (not all images are annotated or predicted with the label).

| Taxon | Precision | Recall | TP seq | FP seq | FN seq | TN seq |
|---|---|---|---|---|---|---|
| Wild boar<br>*Sus scrofa* | 99.88 % | 89.7 % | 845 | 1 | 97 | 1297 |
| Mouflon<br>*Ovis aries* | 98.11 % | 59.92 % | 468 | 9 | 313 | 1450 |
| Red fox<br>*Vulpes vulpes* | 100 % | 78.64 % | 81 | 0 | 22 | 2137 |
| Small mustelids<br>*Martes/Mustela* spp. | 100 % | 68.57 % | 24 | 0 | 11 | 2205 |
| Human<br>*Homo sapiens* | 100 % | 100 % | 23 | 0 | 0 | 2217 |

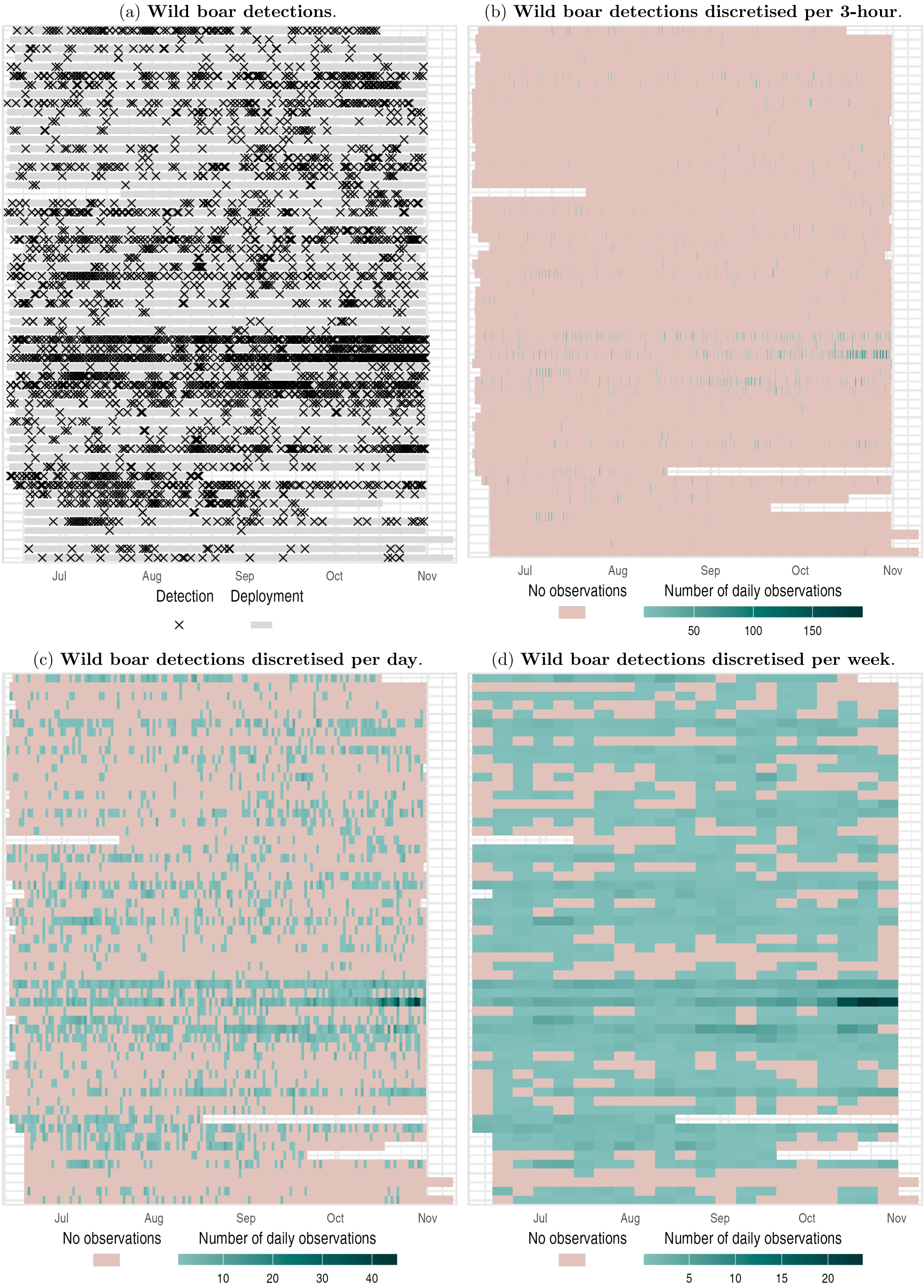

Figure S6: **Detection history for the wild boar**. Rows represent cameras. With 5,219 detections across 58 of 59 cameras. Panels show observations in continuous-time and aggregated in discrete-time with 3-hour, 1-day, and 1-week intervals. In discrete-time plots, to reflect incomplete sessions, the number of daily observations is displayed instead of the counts.

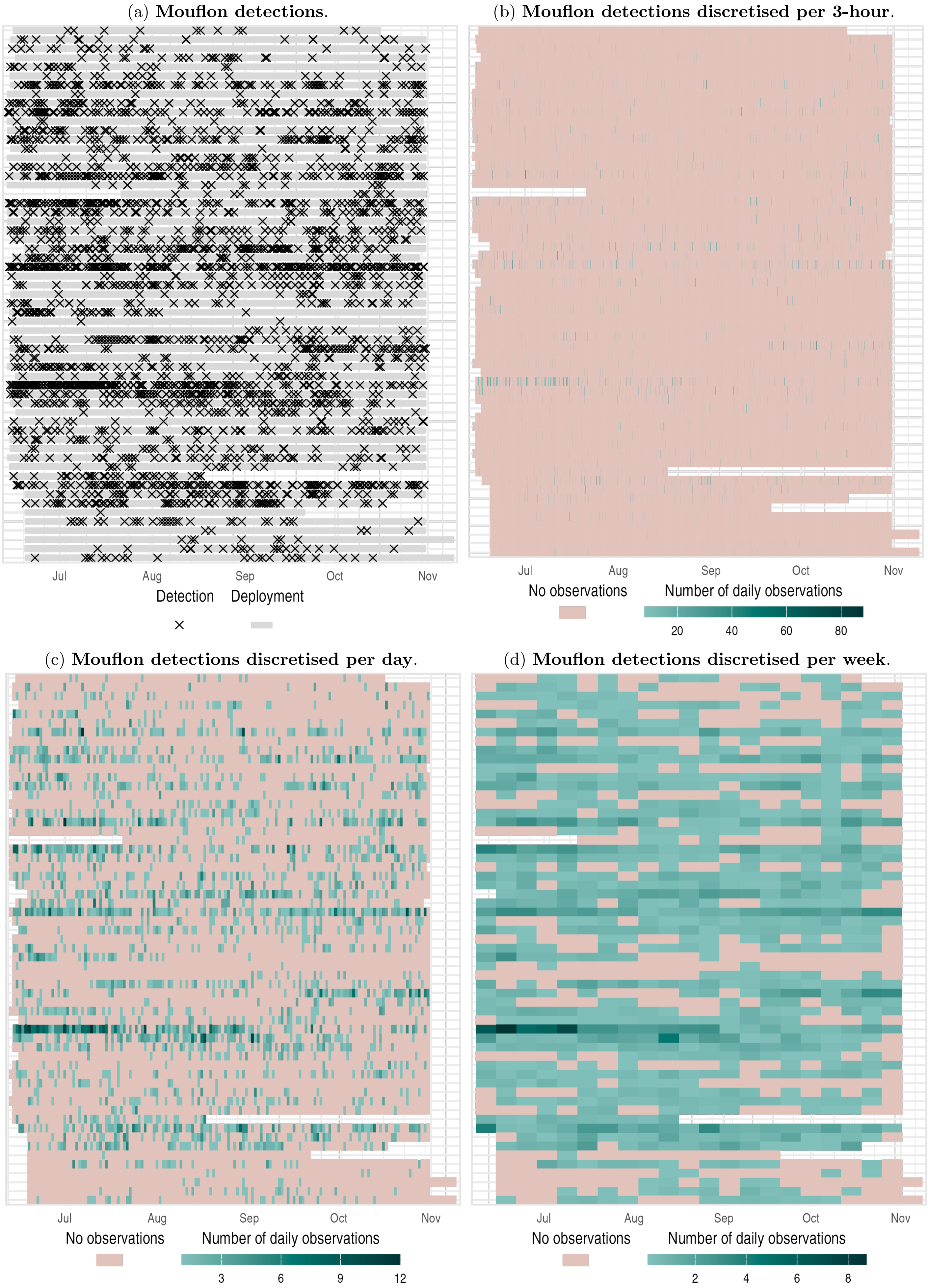

Figure S7: **Detection history for the mouflon**. Rows represent cameras. With 4,032 detections across 59 of 59 cameras. Panels show observations in continuous-time and aggregated in discrete-time with 3-hour, 1-day, and 1-week intervals. In discrete-time plots, to reflect incomplete sessions, the number of daily observations is displayed instead of the counts.

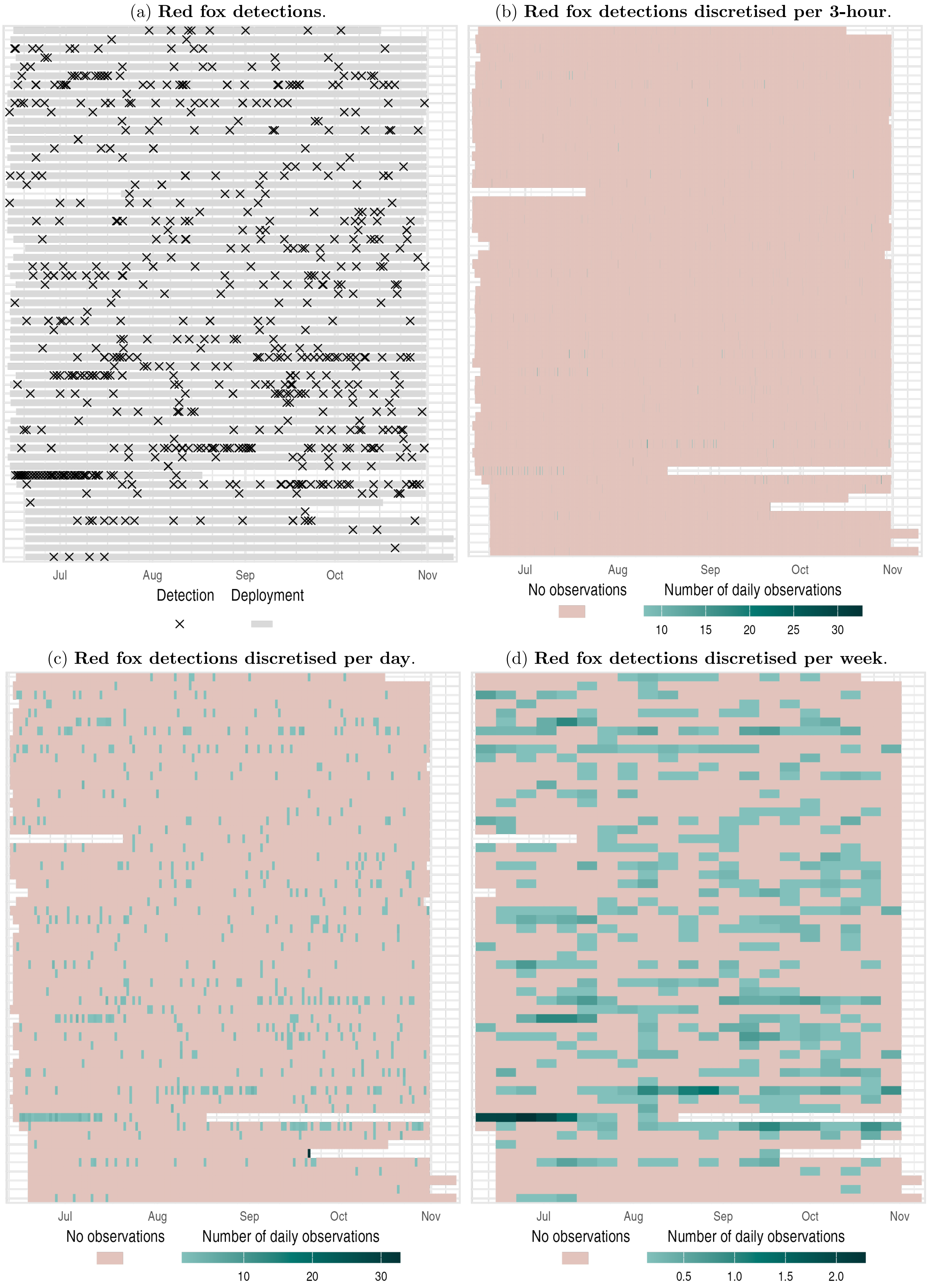

Figure S8: **Detection history for the red fox.** Rows represent cameras. With 689 detections across 59 of 59 cameras. Panels show observations in continuous-time and aggregated in discrete-time with 3-hour, 1-day, and 1-week intervals. In discrete-time plots, to reflect incomplete sessions, the number of daily observations is displayed instead of the counts.

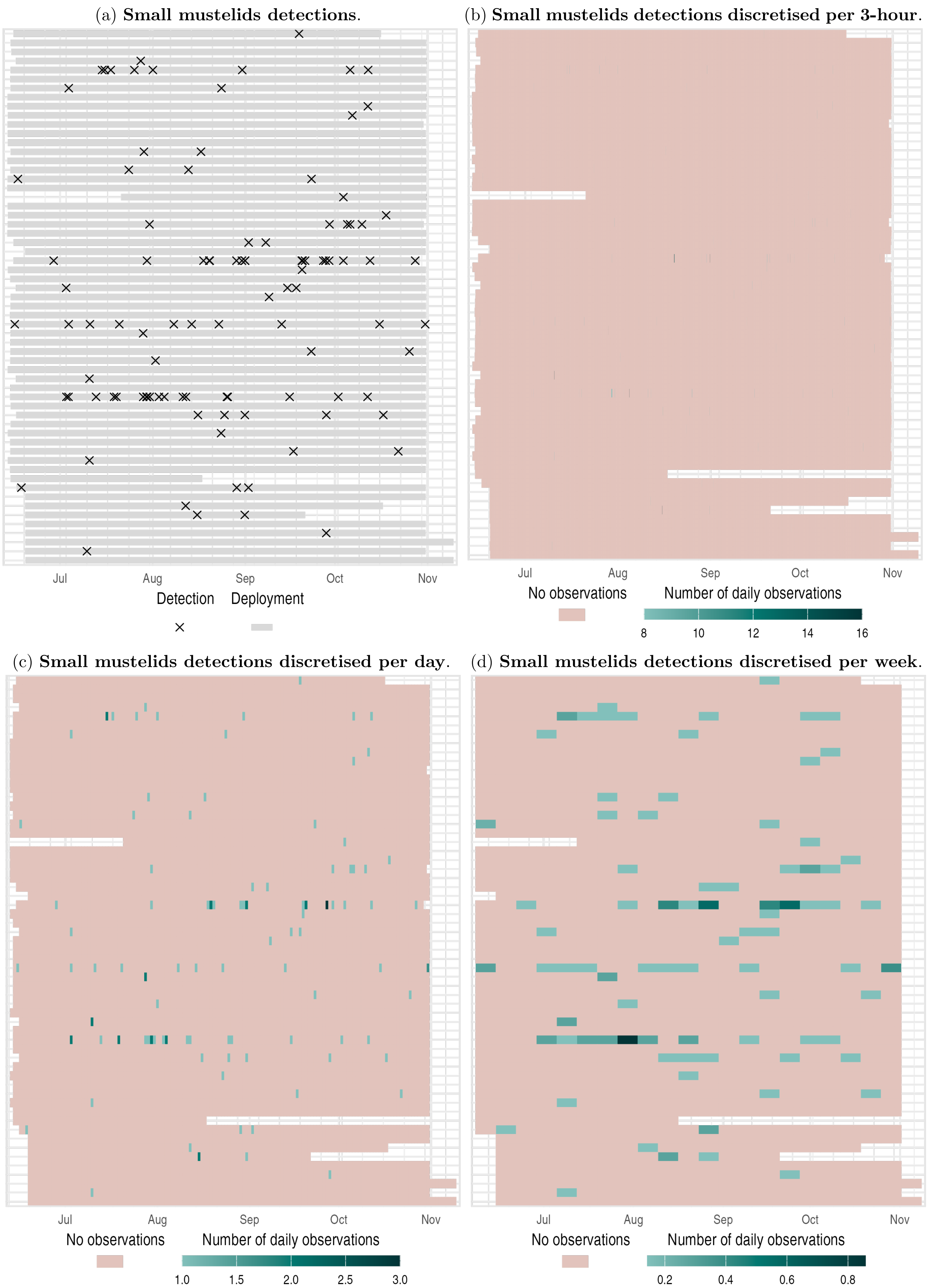

Figure S9: **Detection history for the small mustelids**. Rows represent cameras. With 109 detections across 32 of 59 cameras. Panels show observations in continuous-time and aggregated in discrete-time with 3-hour, 1-day, and 1-week intervals. In discrete-time plots, to reflect incomplete sessions, the number of daily observations is displayed instead of the counts.

###### S2.2.2 Covariates preparation

The numerous covariates were reduced in dimensionality using three separate PCAs: one for habitat covariates (spatial), one for weather covariates (temporal), and one for human disturbance covariates (spatiotemporal). In the models, we used varying temporal precisions, two interpolations methods for CT models, and three discretisation periods for DT models.

- **Habitat covariates (spatial)**
  - Figure S10: land-use map of the study area and sampling design
  - Figure S11: PCA eigenvalues and variable projections
  - Figure S12: spatial representation of selected PCA dimensions
  - Figure S13: correlations between original variables and PCA dimensions
- **Weather covariates (temporal)**
  - Figure S14: PCA eigenvalues and variable projections
  - Figure S15: correlations between original variables and PCA dimensions
  - Figure S16: temporal visualisation at different temporal precisions
- **Human disturbance covariates (spatiotemporal)**
  - Figure S17: PCA eigenvalues and variable projections
  - Figure S18: correlations between original variables and PCA dimensions
  - Figure S19: temporal visualisation at different temporal precisions (3 example sites)

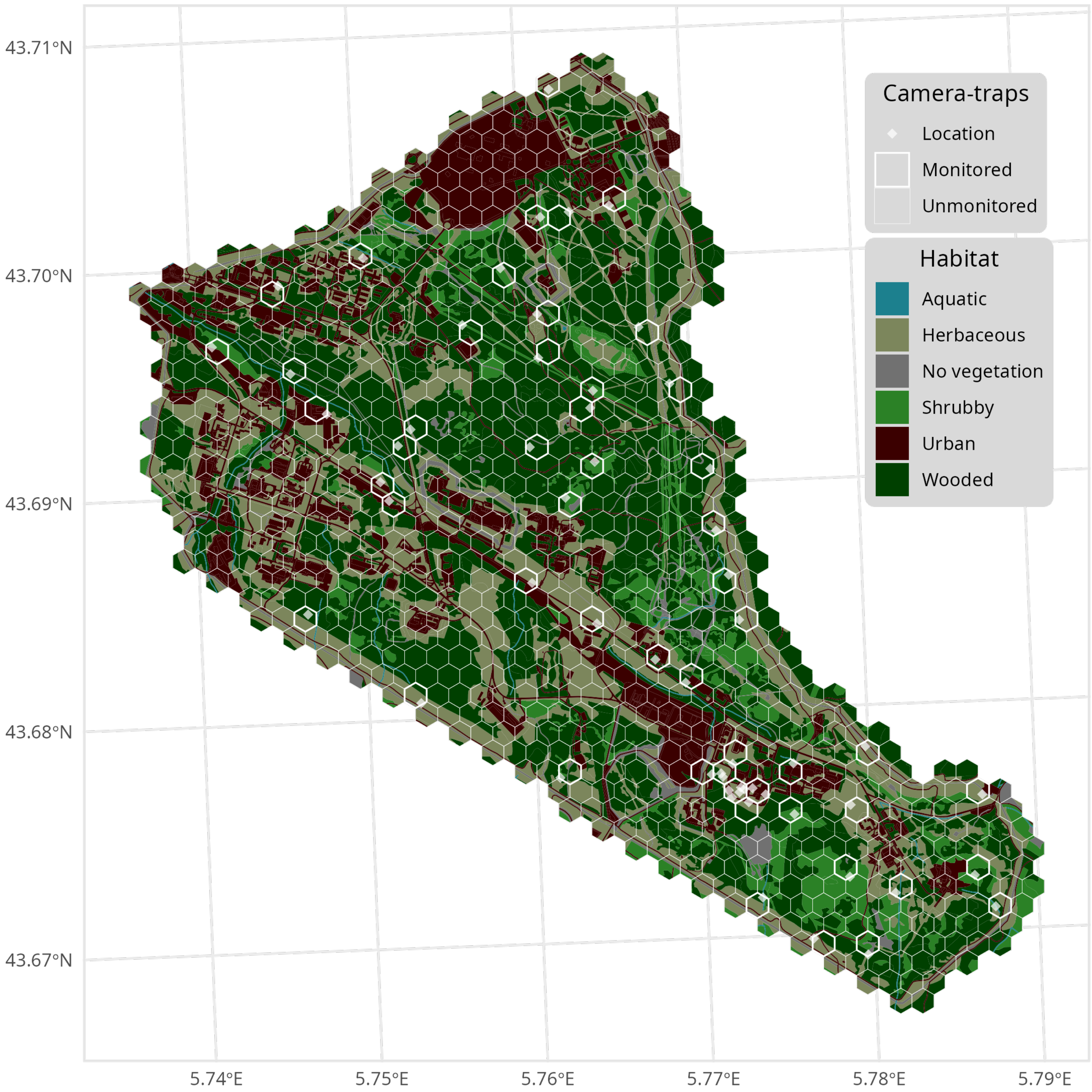

Figure S10: **Land-use map of the study area and sampling design**. Hexagonal cells with a 1-ha area overlay the land-use map, coloured by habitat (6 classes). Points show camera traps. Cells with thick lines highlight monitored cells.

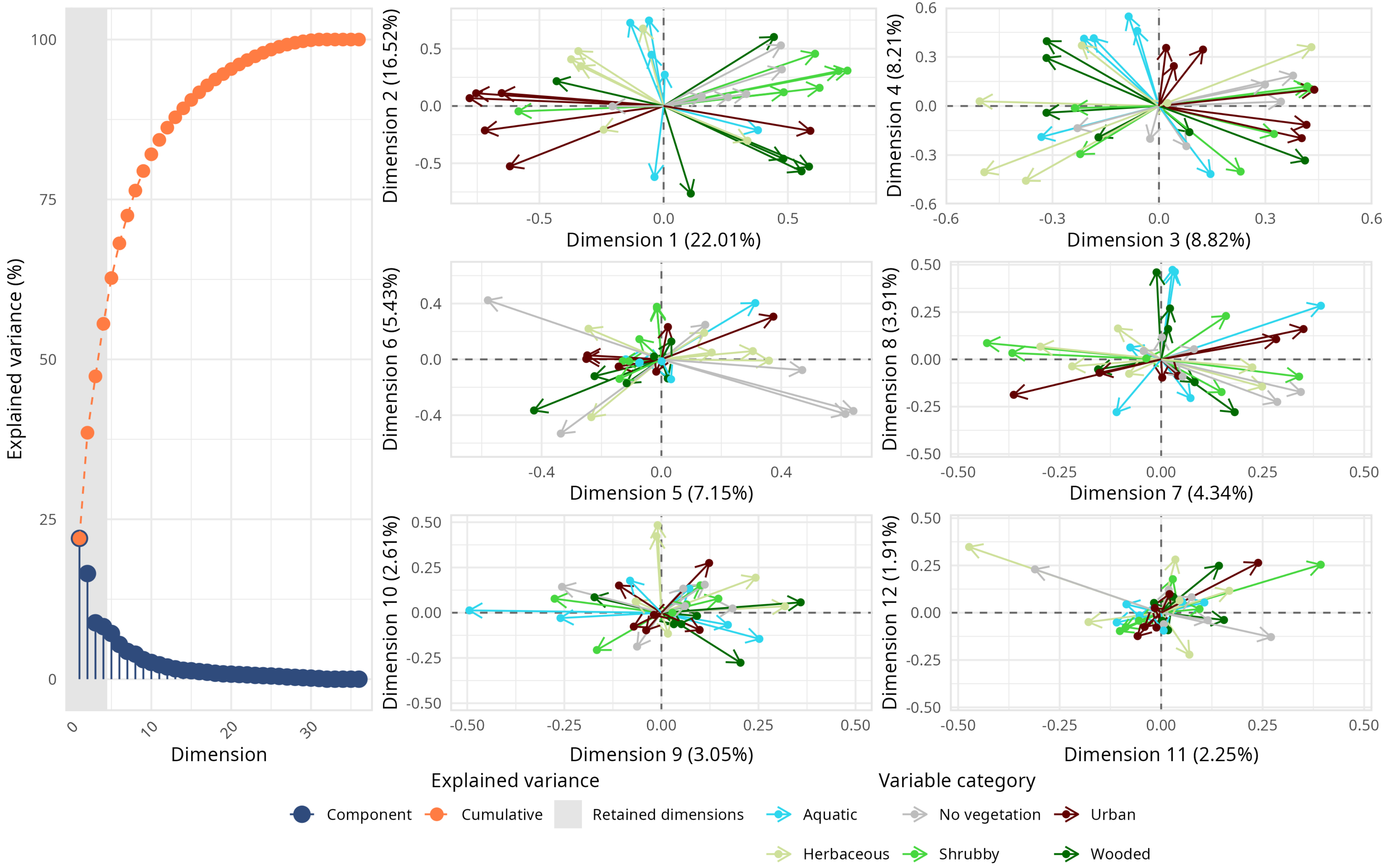

Figure S11: **Habitat PCA eigenvalues and variable projections**. Four dimensions were retained for the rest of the analysis.

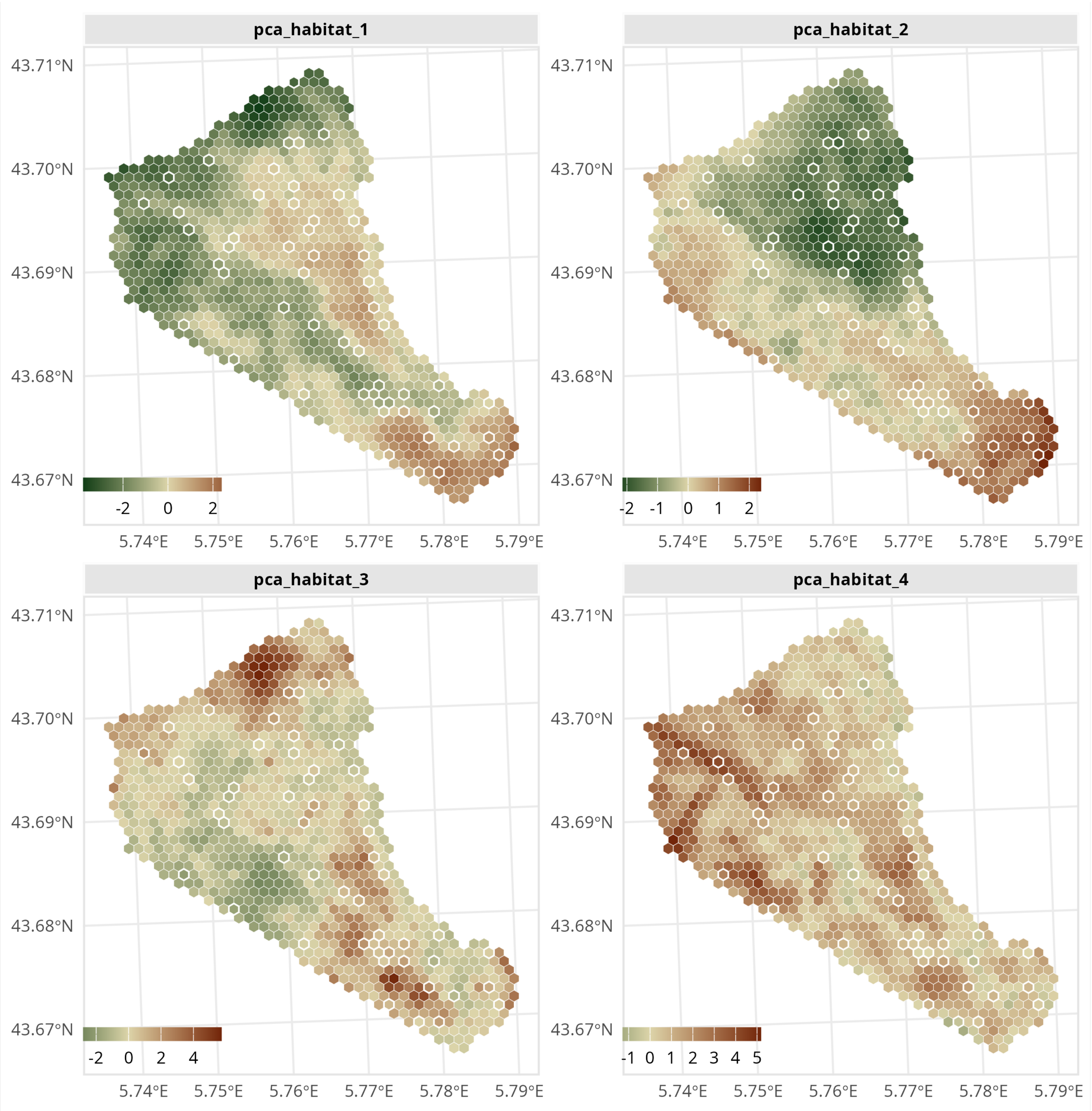

Figure S12: **Habitat PCA spatial visualisation**. Shown are the four retained PCA dimensions, projected within each 1-ha cell of the gridded study area.

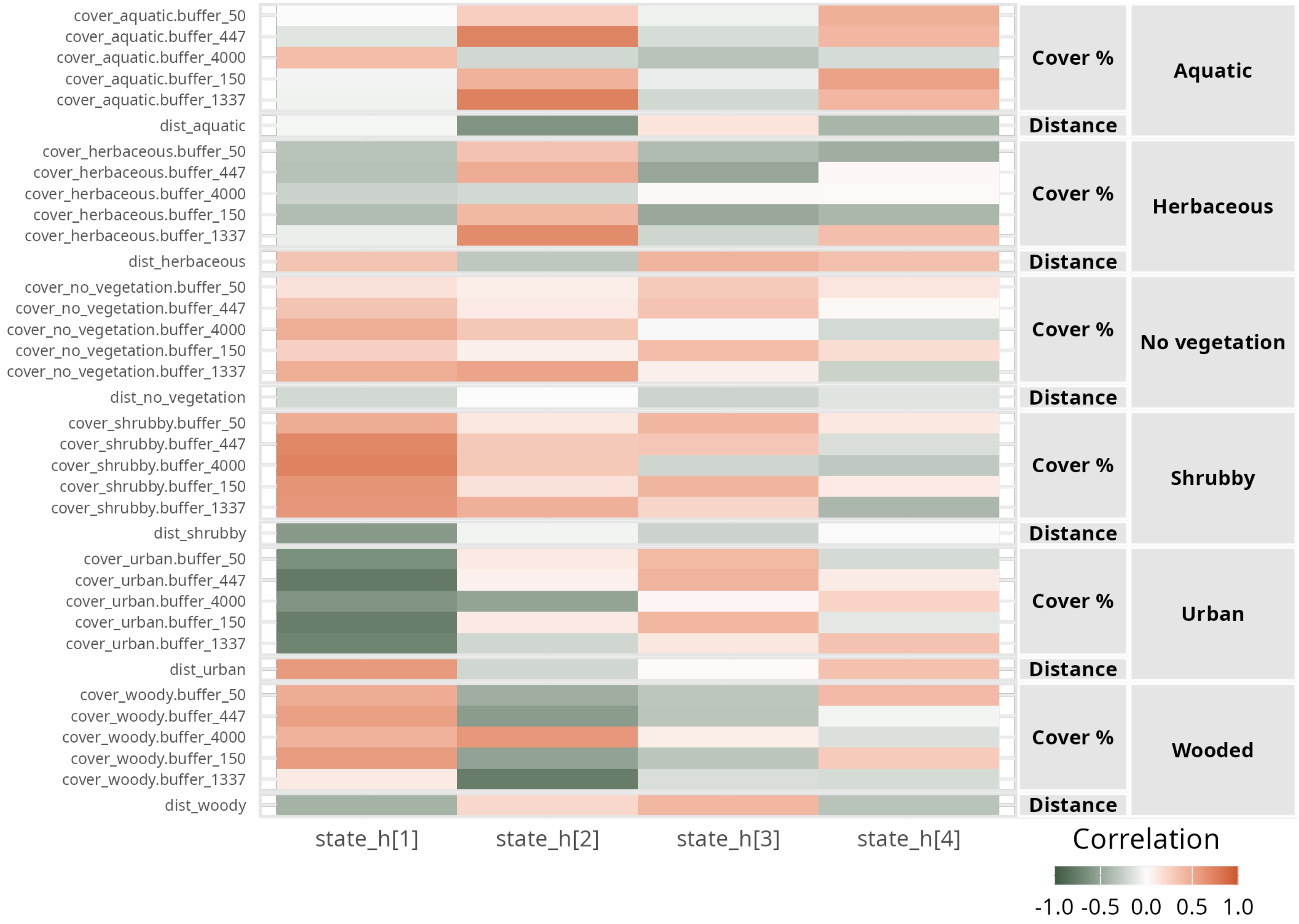

Figure S13: **Habitat PCA: correlations between variables and PCA dimensions**. The four retained PCA dimensions are shown. Darker colours indicate stronger contributions of variables to a given dimension; green indicates negative correlation, orange positive. Variables are prefixed with `dist` when they represent the distance to the nearest area of a given habitat type (e.g. `dist_woody` is the distance to the closest woody area), and with `cover` when they represent the percentage cover of a habitat type within a defined area (e.g. `cover_woody.buffer_150` is the percentage of forest within a 150-m buffer around the cell centroid).

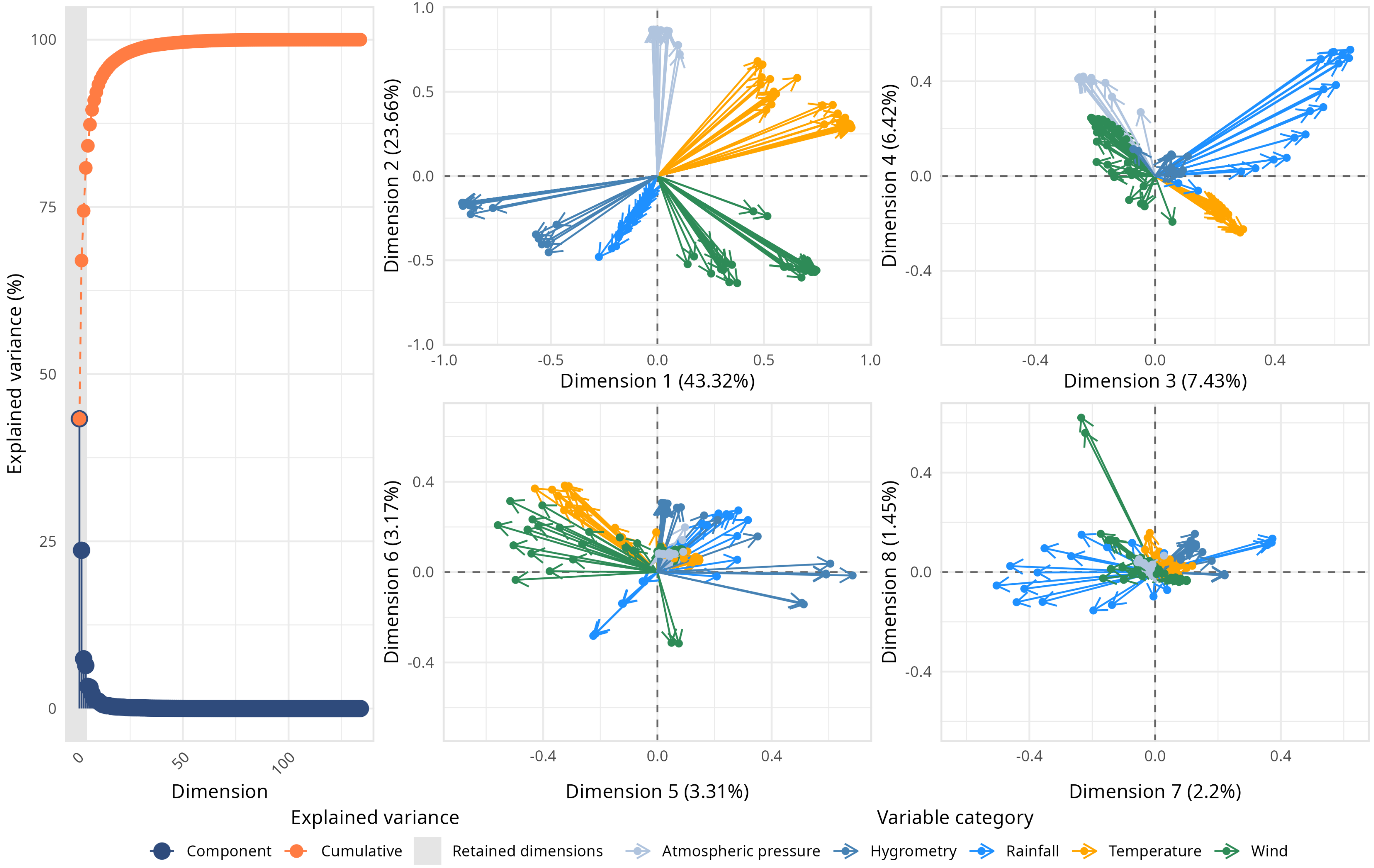

Figure S14: **Weather PCA eigenvalues and variable projections.** Four dimensions were retained for the rest of the analysis.

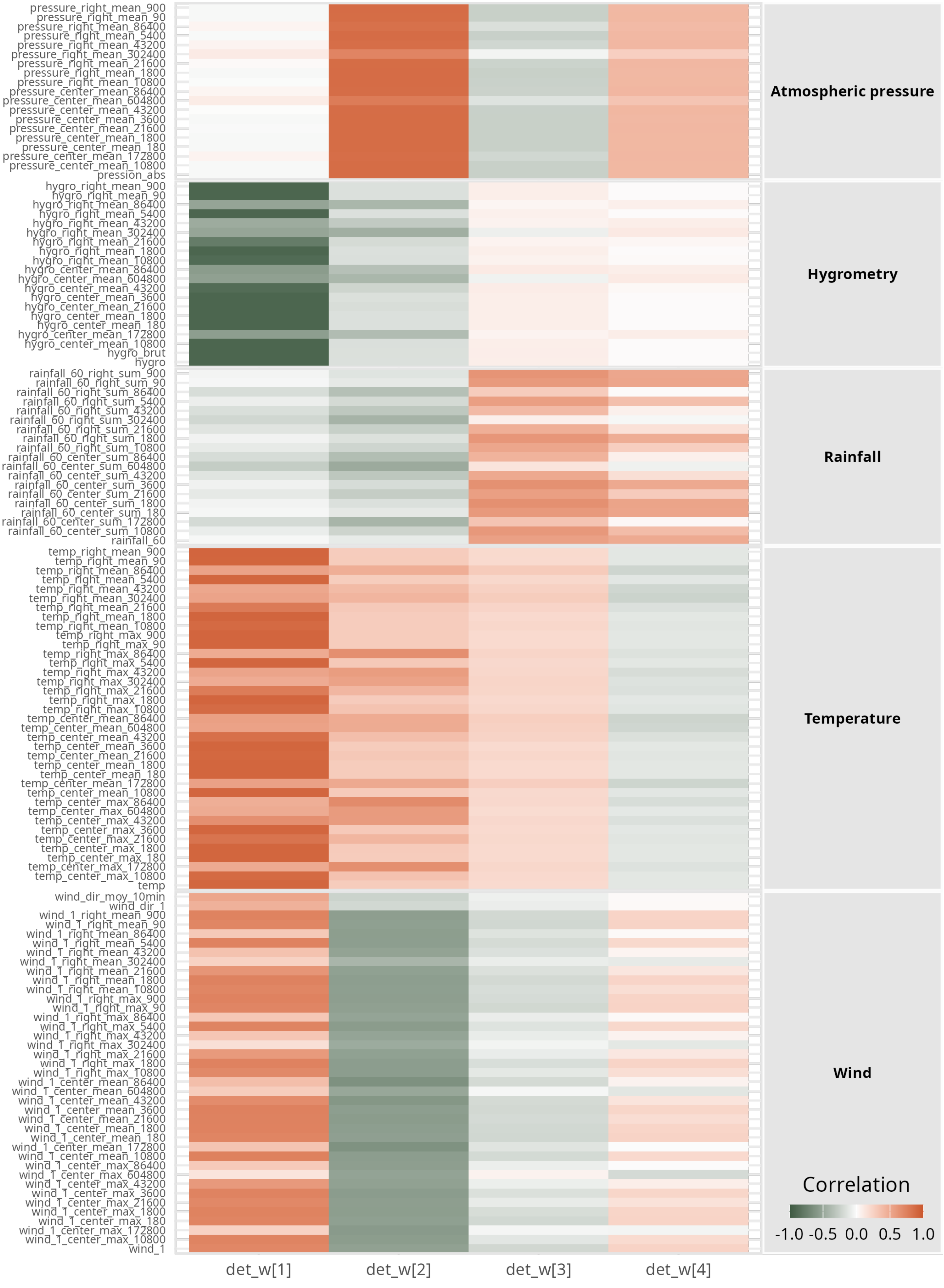

Figure S15: **Weather PCA: correlations between variables and PCA dimensions**. Shown are the four retained PCA dimensions, hereafter used as detection covariates (`det_w[1]` to `det_w[4]`). Darker colours indicate stronger retention of variable information in a given PCA dimension; green indicates negative correlation, orange positive. Variables are named with the rolling function (max/sum/mean) and how it was applied (center/right, and the duration in seconds).

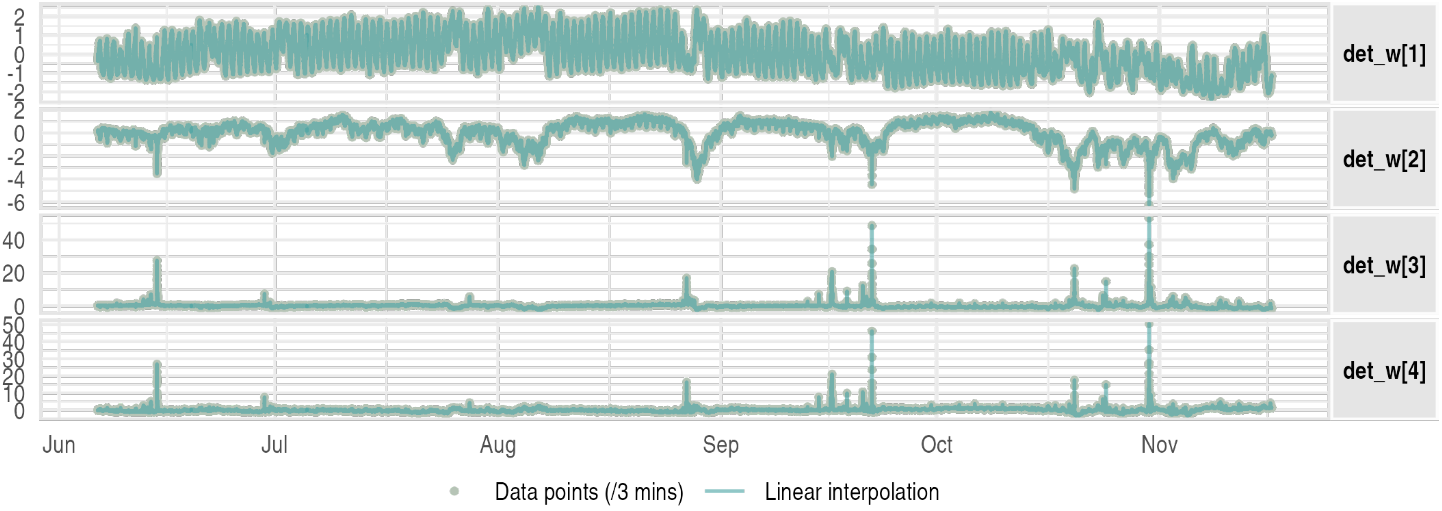

(a) **1 point each 3 minutes**.

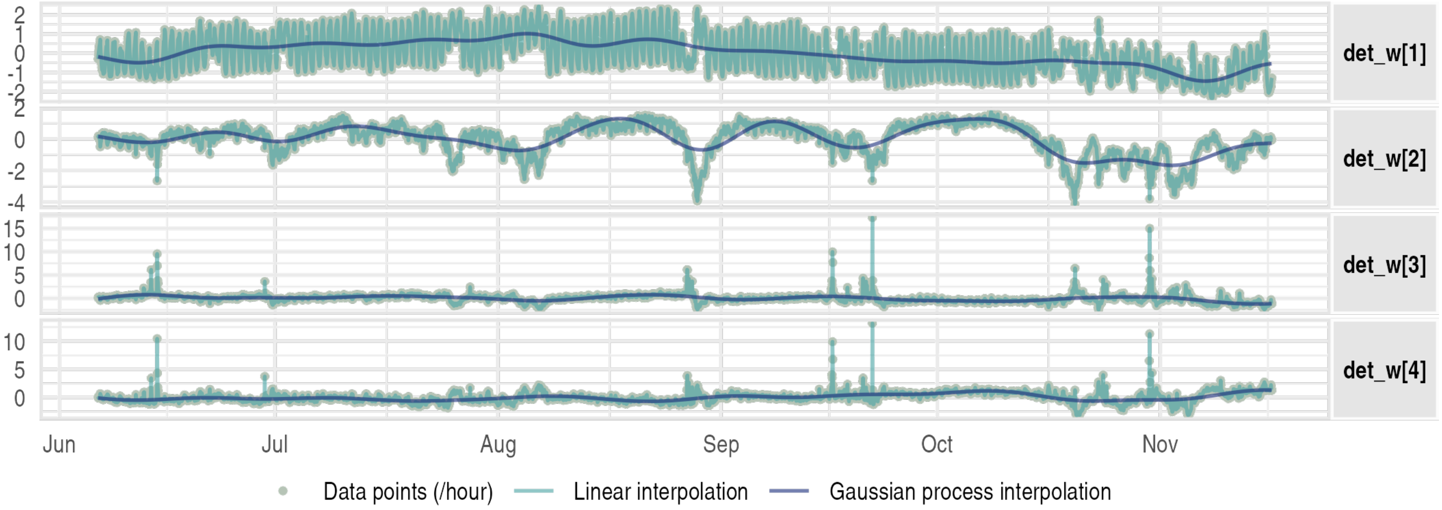

(b) **1 point each hour**.

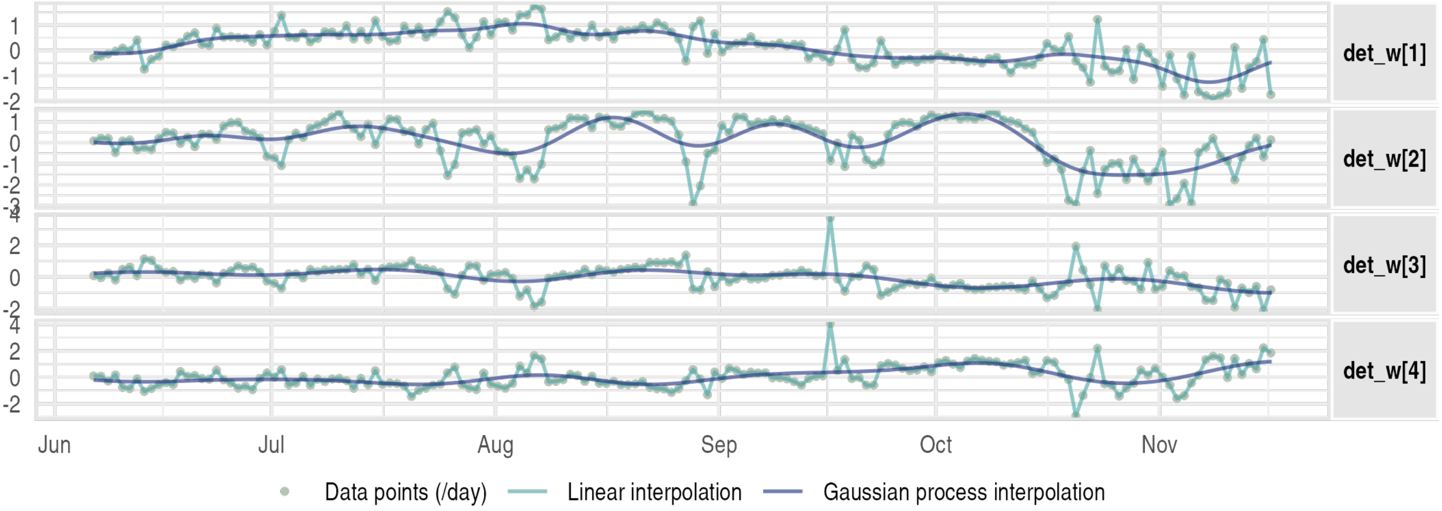

(c) **1 point each day**.

Figure S16: **Weather covariates: temporal visualisation at different temporal precisions**. Data points, linear interpolation and gaussian interpolation are shown. No gaussian interpolation was done for the 3-mins level of temporal precision.

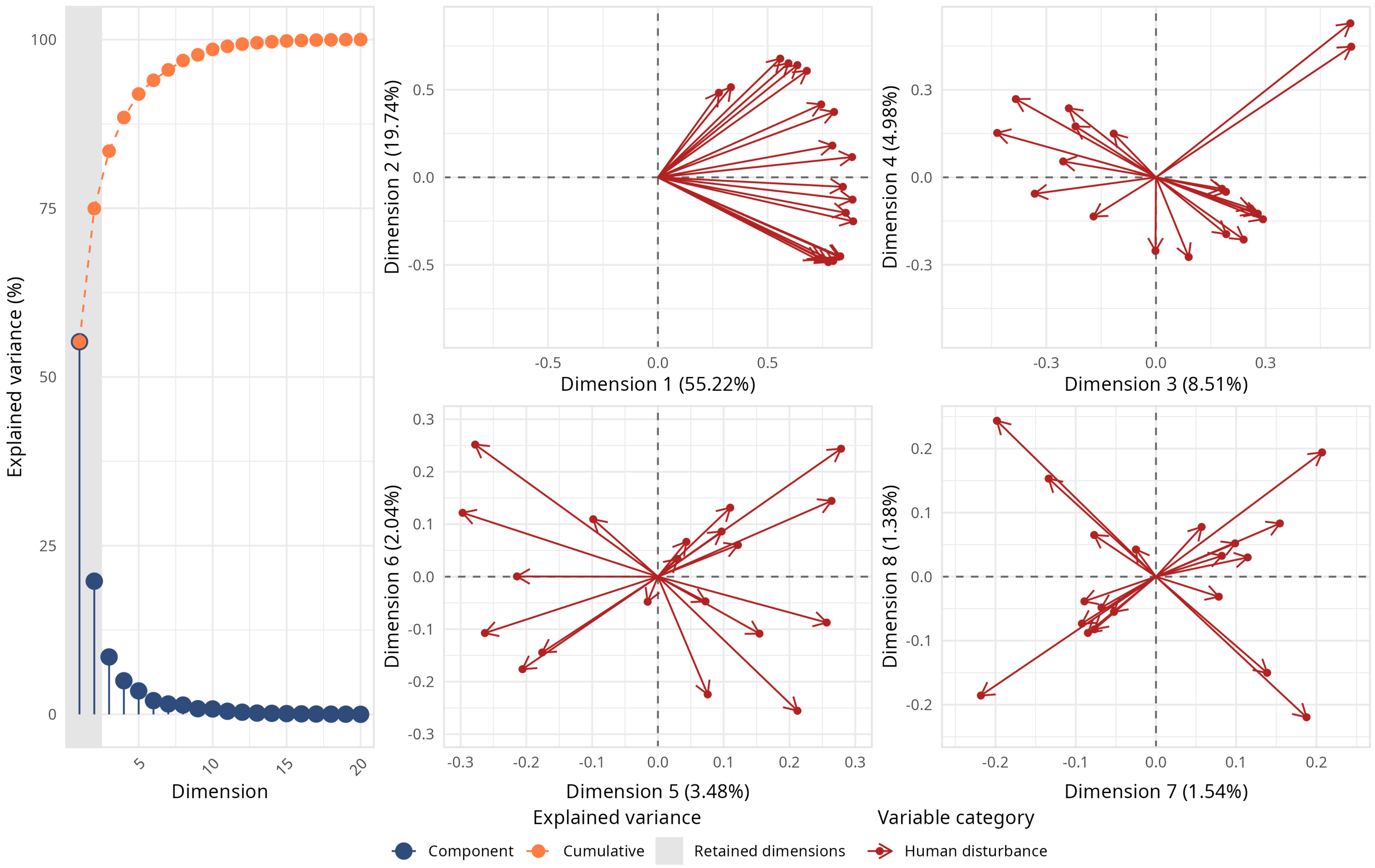

Figure S17: **Human disturbance PCA eigenvalues and variable projections.** Two dimensions were retained for the rest of the analysis.

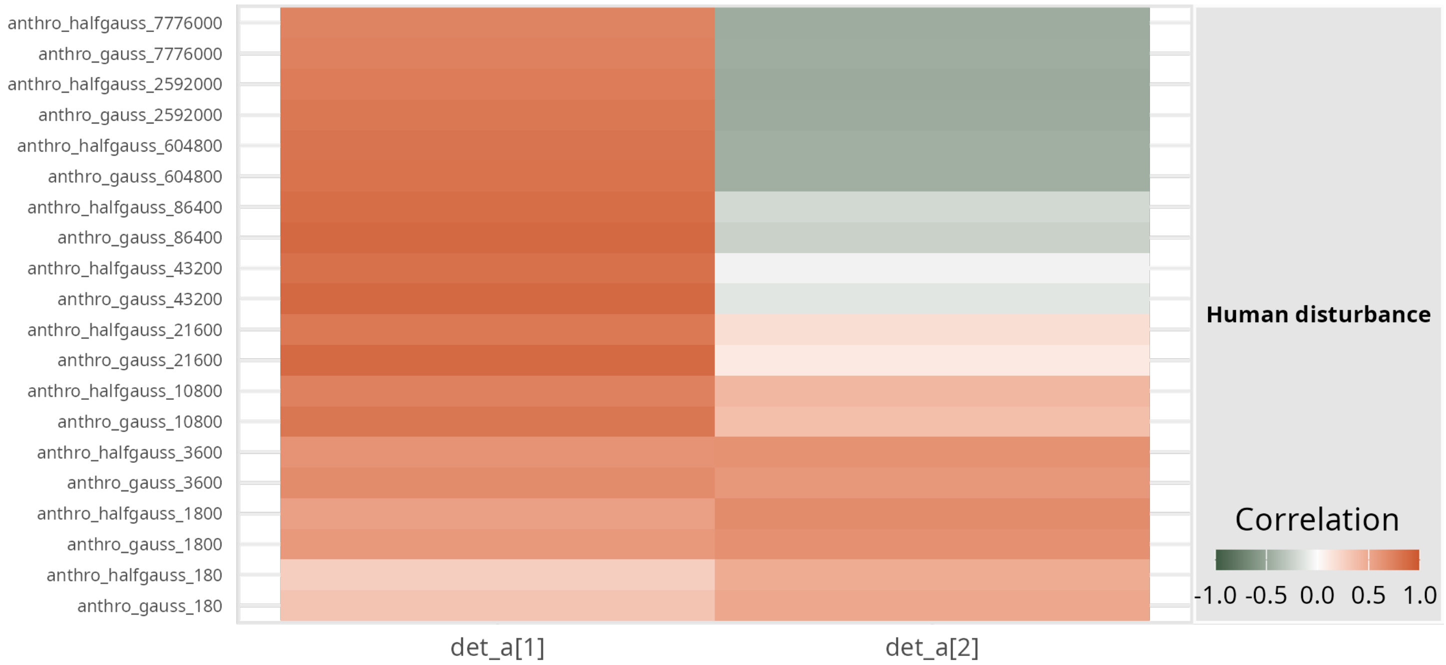

Figure S18: **Human disturbance PCA: correlations between variables and PCA dimensions.** Shown are the two retained PCA dimensions, hereafter used as detection covariates (`det_a[1]`, `det_a[2]`). Darker colours indicate stronger retention of variable information in a given PCA dimension; green indicates negative correlation, orange positive. Variables are named with the kernel (gaussian or half-gaussian) and the duration over which the kernel is applied (in seconds).

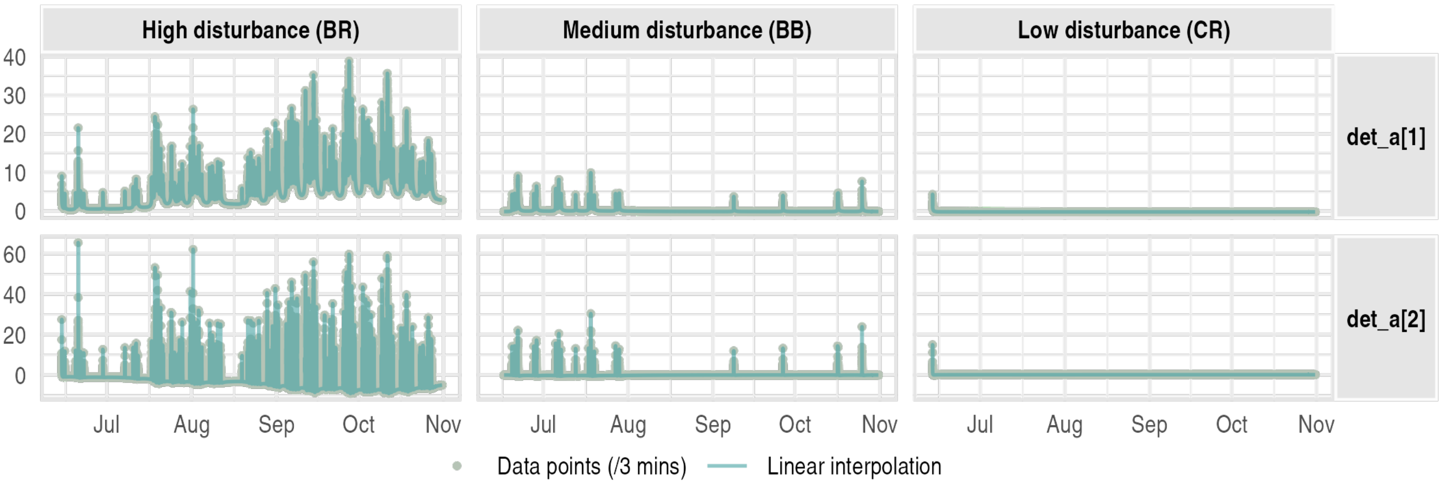

(a) **1 point each 3 minutes**.

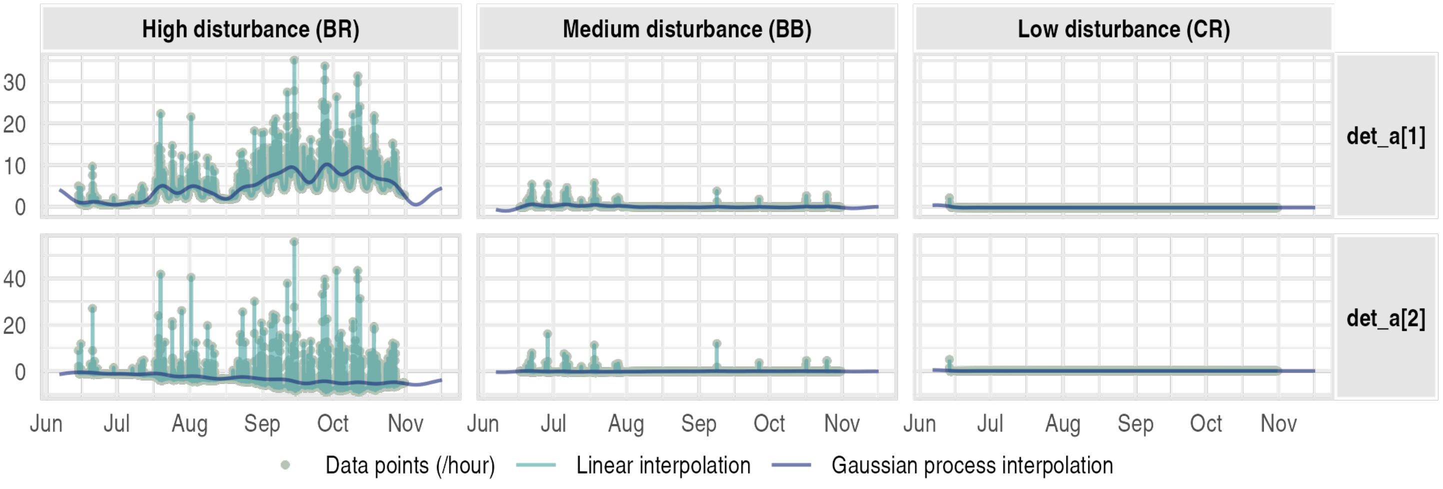

(b) **1 point each hour**.

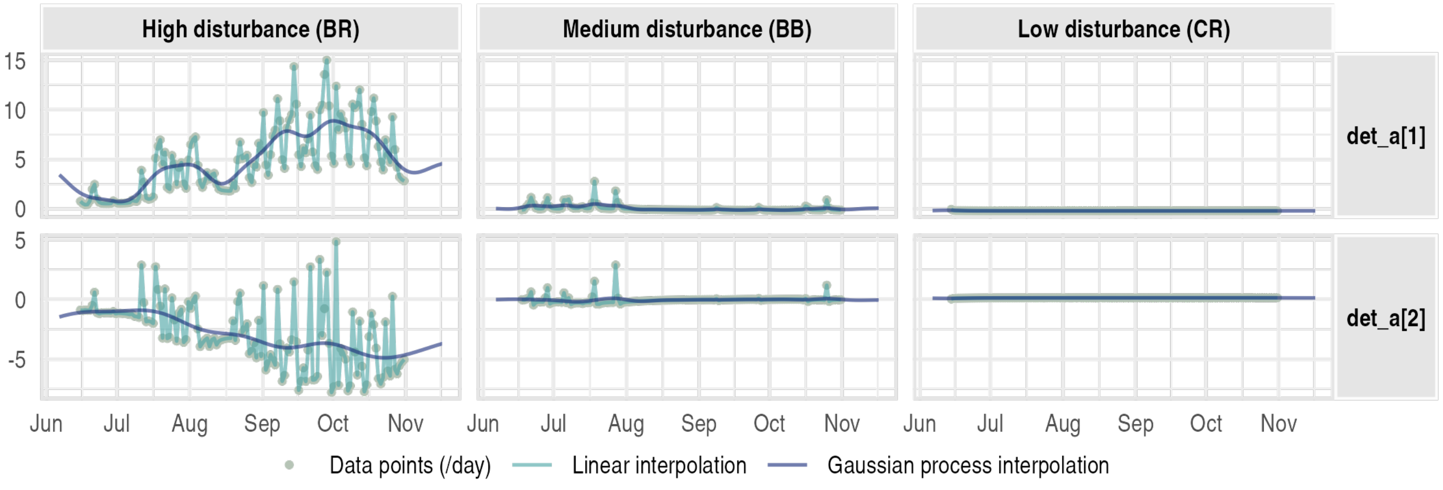

(c) **1 point each day**.

Figure S19: **Human disturbance covariates: temporal visualisation at different temporal precisions**. Data points, linear interpolation and gaussian interpolation are shown. No gaussian interpolation was done for the 3-mins level of temporal precision. Only 3 example sites are shown.

##### S2.2.3 Coefficients estimation

Coefficient estimates for each focal taxon (wild boar, mouflon, red fox, and small mustelids) are presented below. For each species, we display results from four models: the abundance NULL model, the abundance model with covariates, the occupancy NULL model, and the occupancy model with covariates. Only the abundance model with covariates for wild boar is included in the main manuscript figures; all others are presented in this appendix.

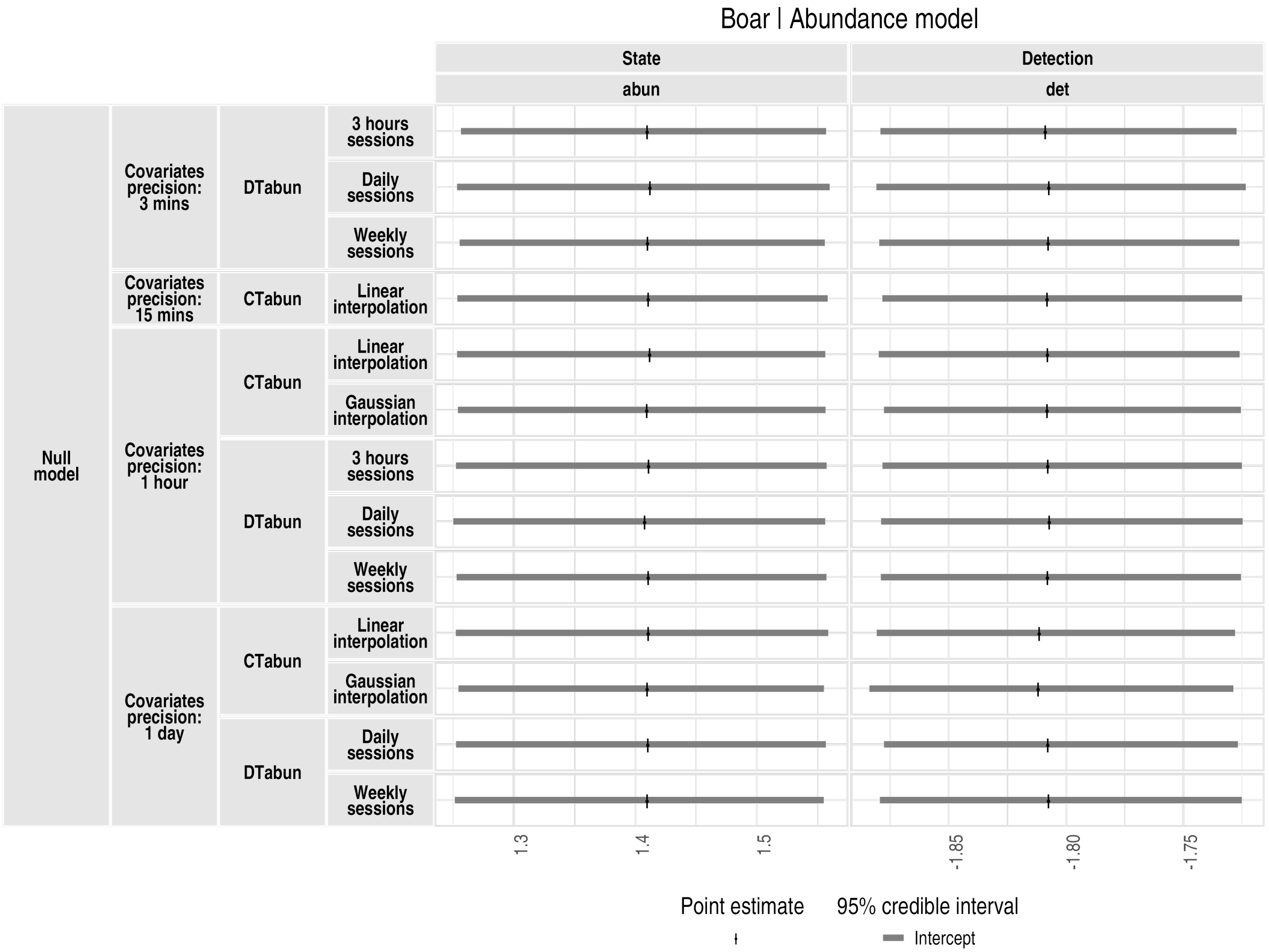

Figure S20: **Abundance model results for the wild boar (null model)**. `abun` refers to the abundance intercept (log scale) and `det` to the detection rate intercept (log scale). Grey horizontal lines show 95% credible intervals (posterior distribution quantiles), and black vertical lines indicate point estimates (posterior distribution medians). Model types and configurations are listed on the left side of the plot.

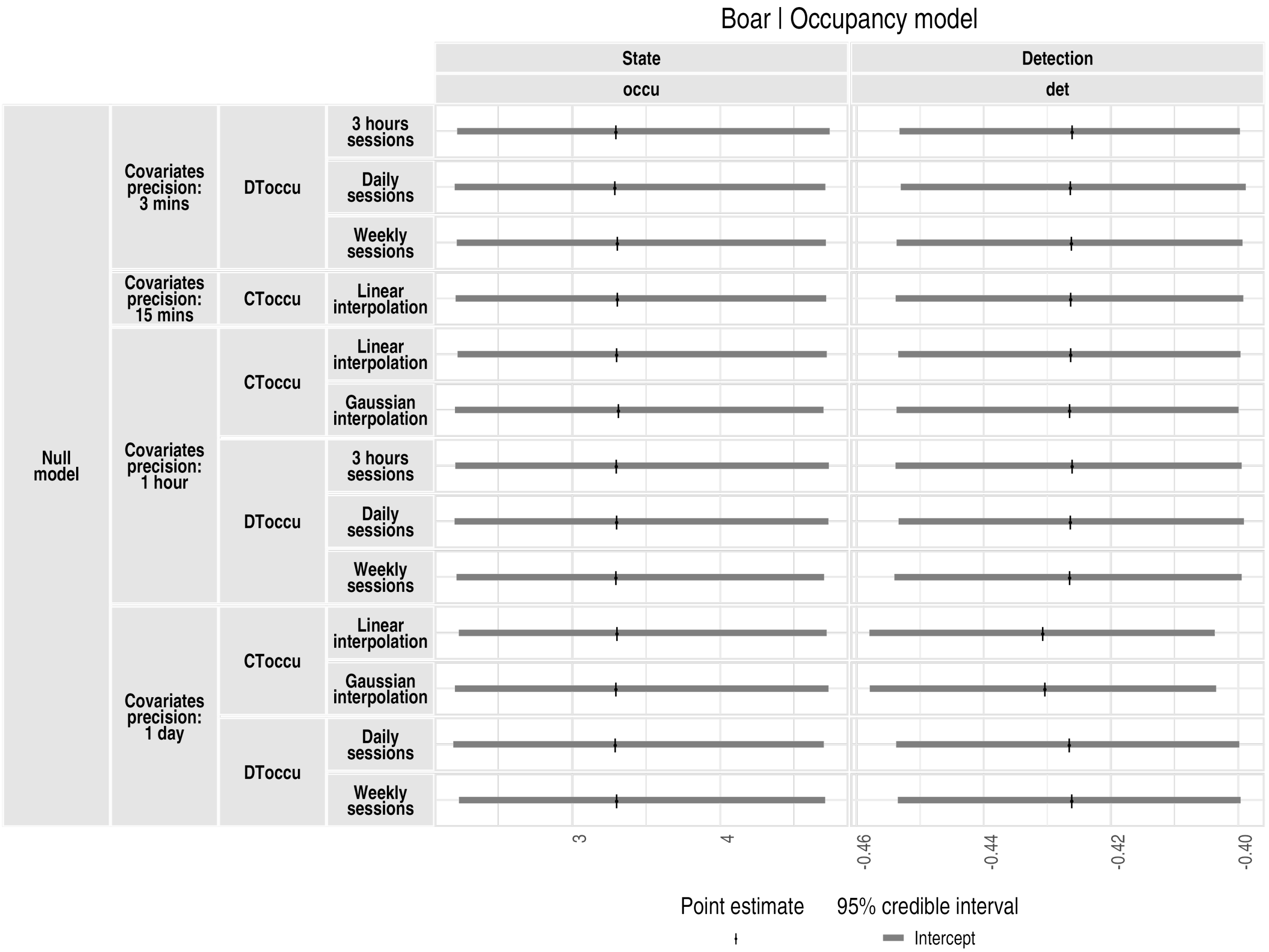

Figure S21: **Occupancy model results for the wild boar (null model)**. `occu` refers to the occupancy intercept (logit scale) and `det` to the detection rate intercept (log scale). Grey horizontal lines show 95% credible intervals (posterior distribution quantiles), and black vertical lines indicate point estimates (posterior distribution medians). Model types and configurations are listed on the left side of the plot.

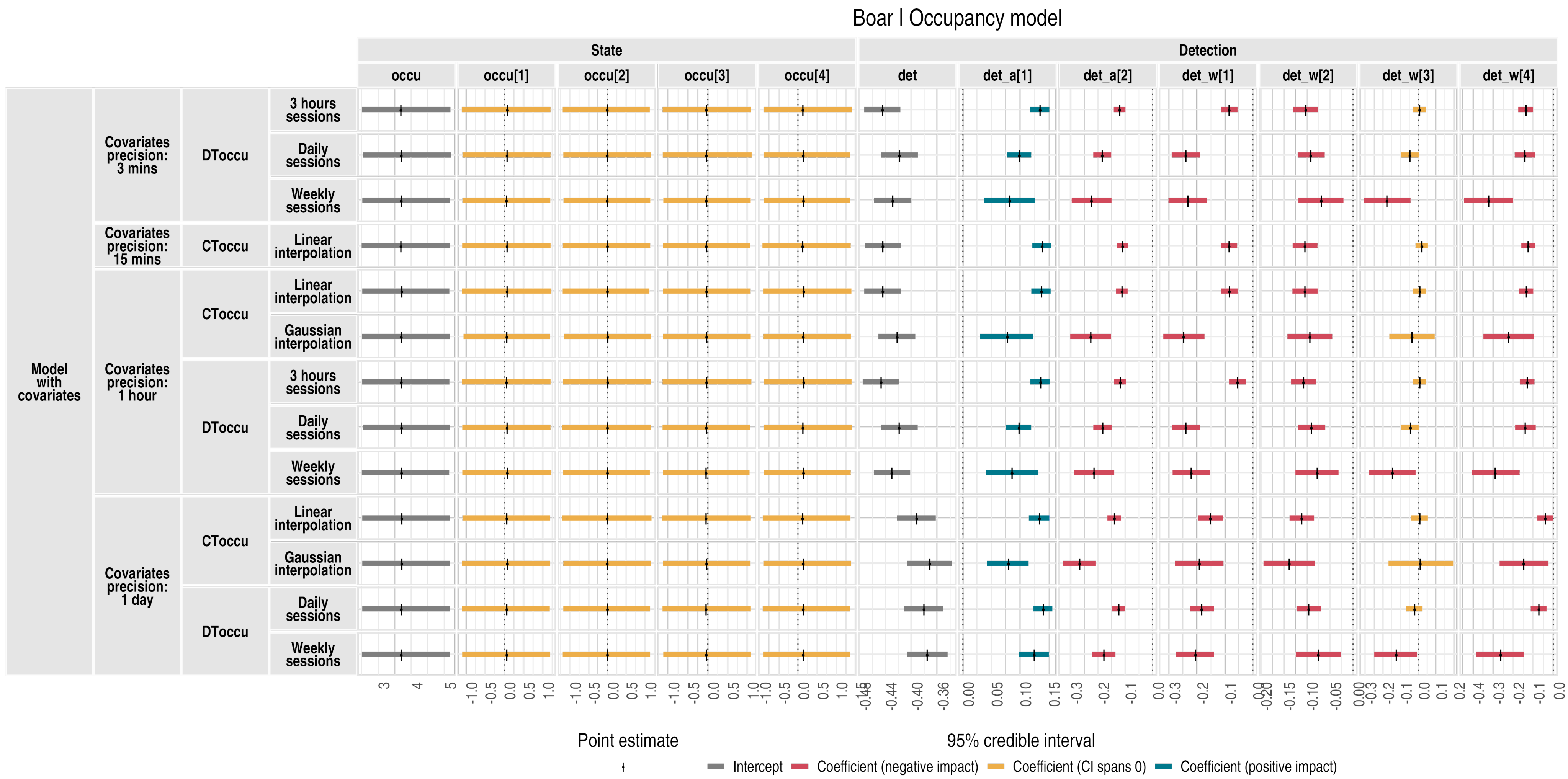

Figure S22: **Occupancy model results for the wild boar**. `occu` is the occupancy intercept, and `occu[1]` to `occu[4]` are the coefficients for habitat covariates (named `state_h[1]` to `state_h[4]` in the covariate plots). `det` is the detection rate intercept; `det_a[1]` and `det_a[2]` are the anthropogenic (human disturbance) detection covariate coefficients; `det_w[1]` to `det_w[4]` are the weather detection covariate coefficients. Grey horizontal lines show 95% credible intervals (posterior distribution quantiles), and black vertical lines indicate point estimates (posterior distribution medians). Model types and configurations are listed on the left side of the plot.

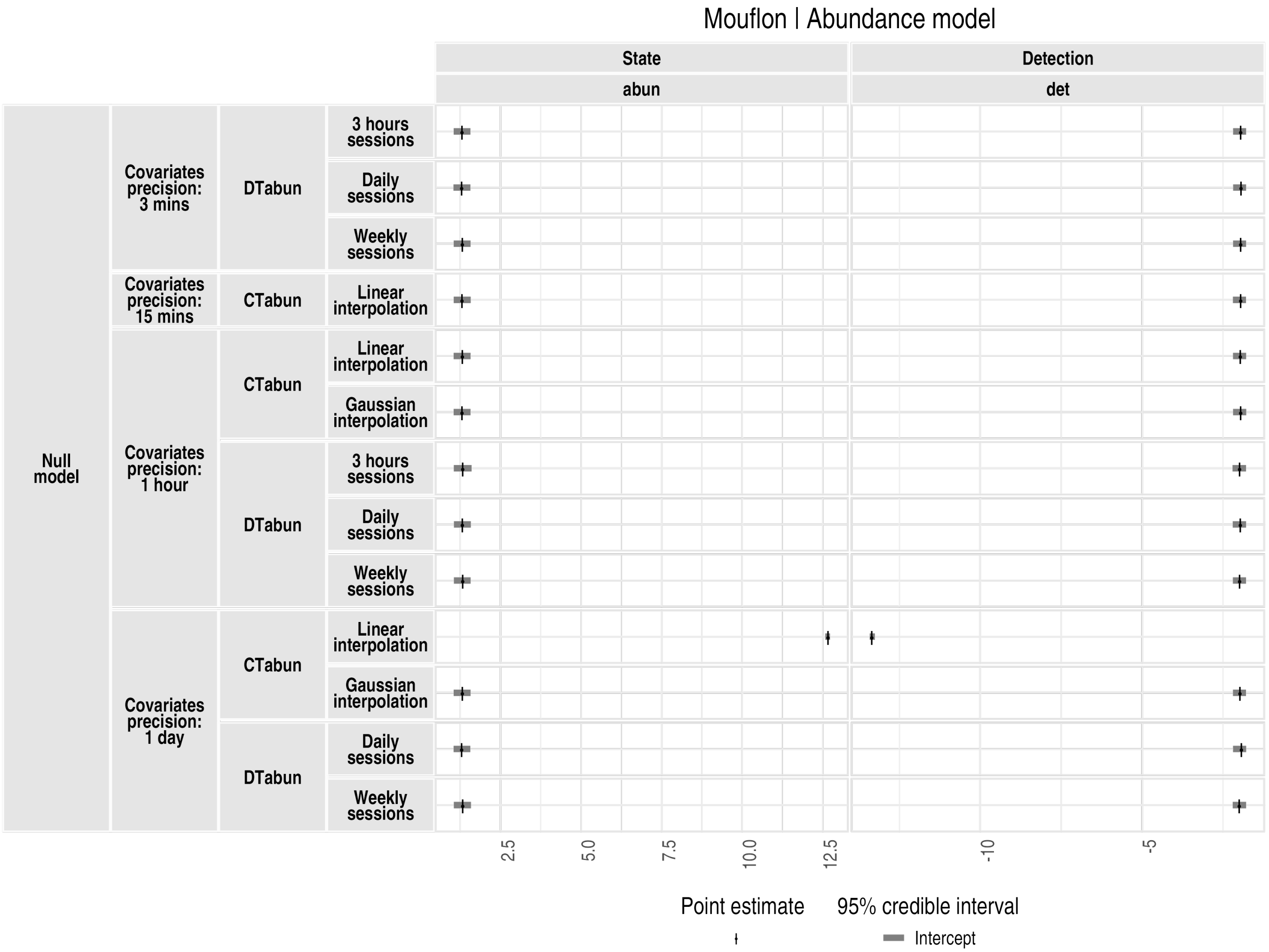

Figure S23: **Abundance model results for the mouflon (null model)**. `abun` refers to the abundance intercept (log scale) and `det` to the detection rate intercept (log scale). Grey horizontal lines show 95% credible intervals (posterior distribution quantiles), and black vertical lines indicate point estimates (posterior distribution medians). Model types and configurations are listed on the left side of the plot.

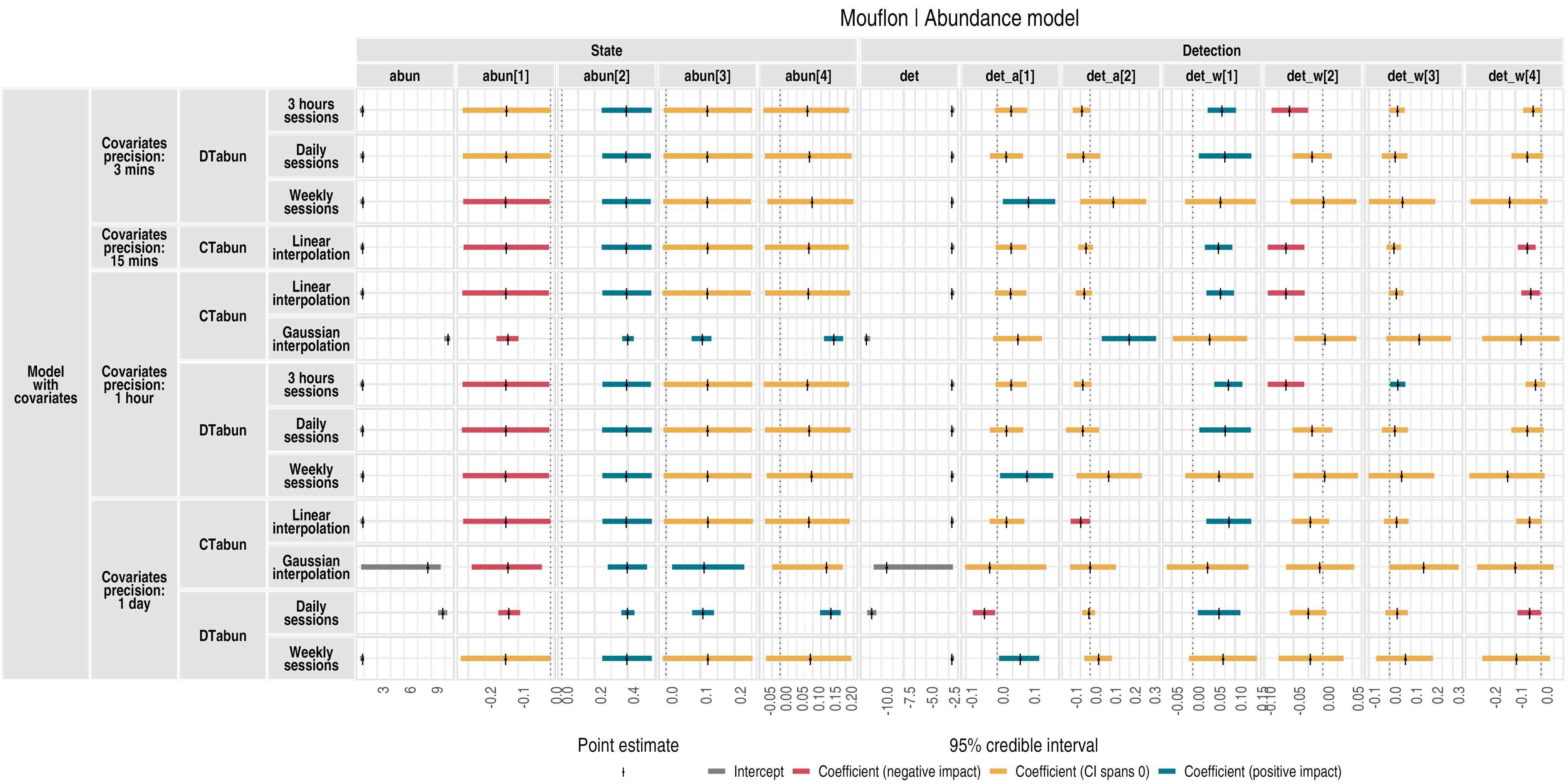

Figure S24: **Abundance model results for the mouflon**. `abun` is the abundance intercept, and `abun[1]` to `abun[4]` are the coefficients for habitat covariates (named `state_h[1]` to `state_h[4]` in the covariate plots). `det` is the detection rate intercept; `det_a[1]` and `det_a[2]` are the anthropogenic (human disturbance) detection covariate coefficients; `det_w[1]` to `det_w[4]` are the weather detection covariate coefficients. Grey horizontal lines show 95% credible intervals (posterior distribution quantiles), and black vertical lines indicate point estimates (posterior distribution medians). Model types and configurations are listed on the left side of the plot.

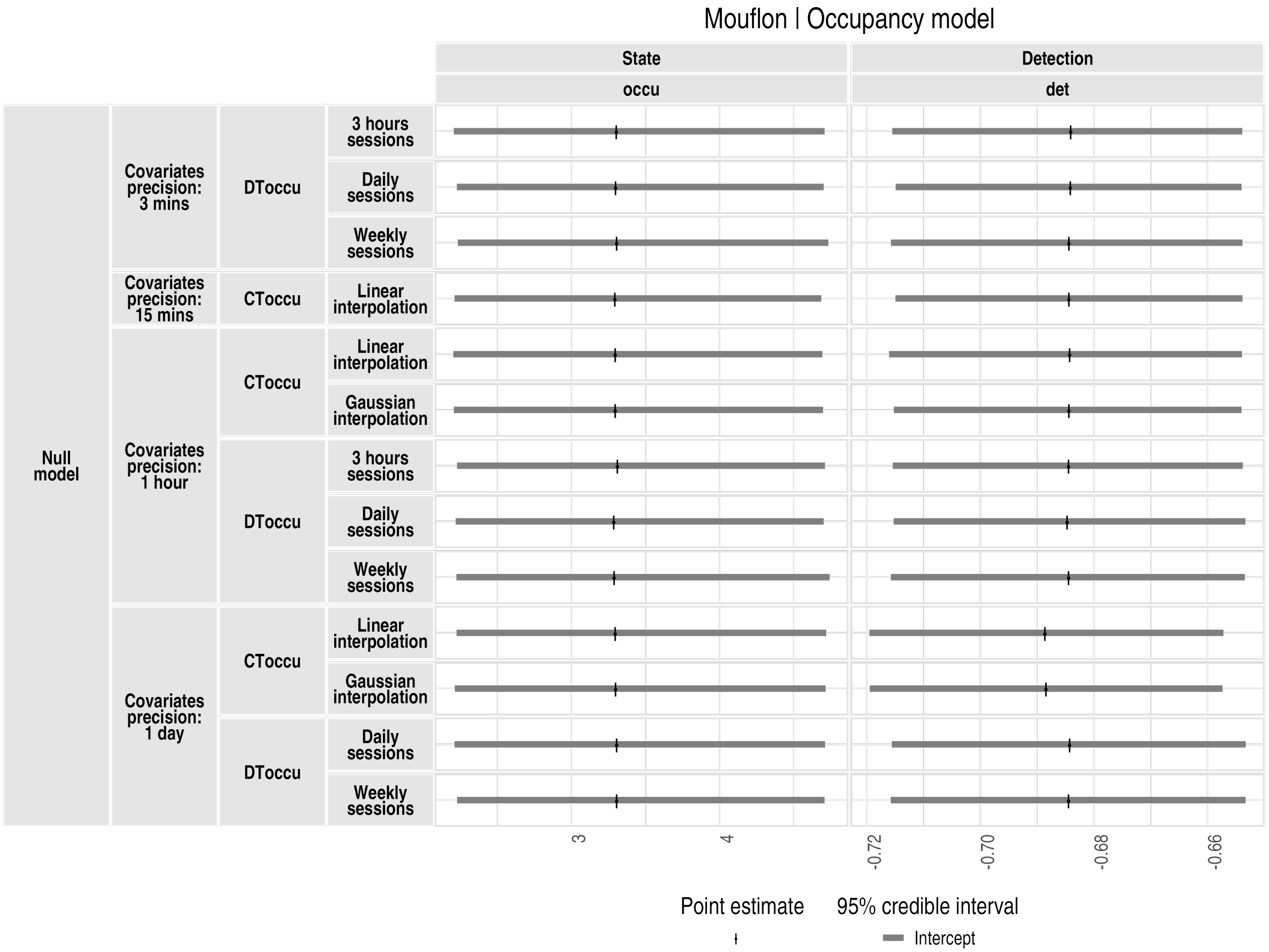

Figure S25: **Occupancy model results for the mouflon (null model)**. `occu` refers to the occupancy intercept (logit scale) and `det` to the detection rate intercept (log scale). Grey horizontal lines show 95% credible intervals (posterior distribution quantiles), and black vertical lines indicate point estimates (posterior distribution medians). Model types and configurations are listed on the left side of the plot.

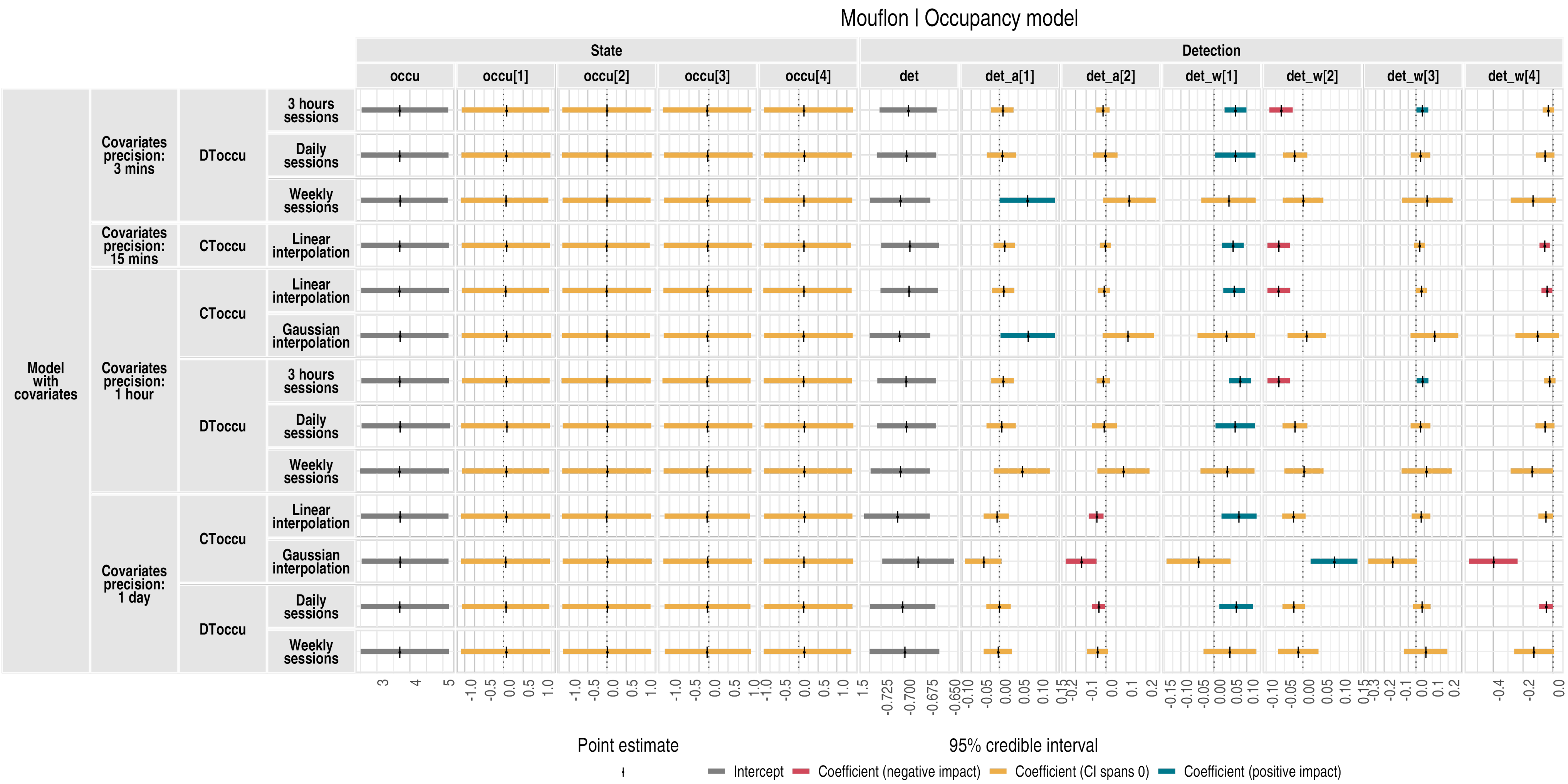

Figure S26: **Occupancy model results for the mouflon**. `occu` is the occupancy intercept, and `occu[1]` to `occu[4]` are the coefficients for habitat covariates (named `state_h[1]` to `state_h[4]` in the covariate plots). `det` is the detection rate intercept; `det_a[1]` and `det_a[2]` are the anthropogenic (human disturbance) detection covariate coefficients; `det_w[1]` to `det_w[4]` are the weather detection covariate coefficients. Grey horizontal lines show 95% credible intervals (posterior distribution quantiles), and black vertical lines indicate point estimates (posterior distribution medians). Model types and configurations are listed on the left side of the plot.

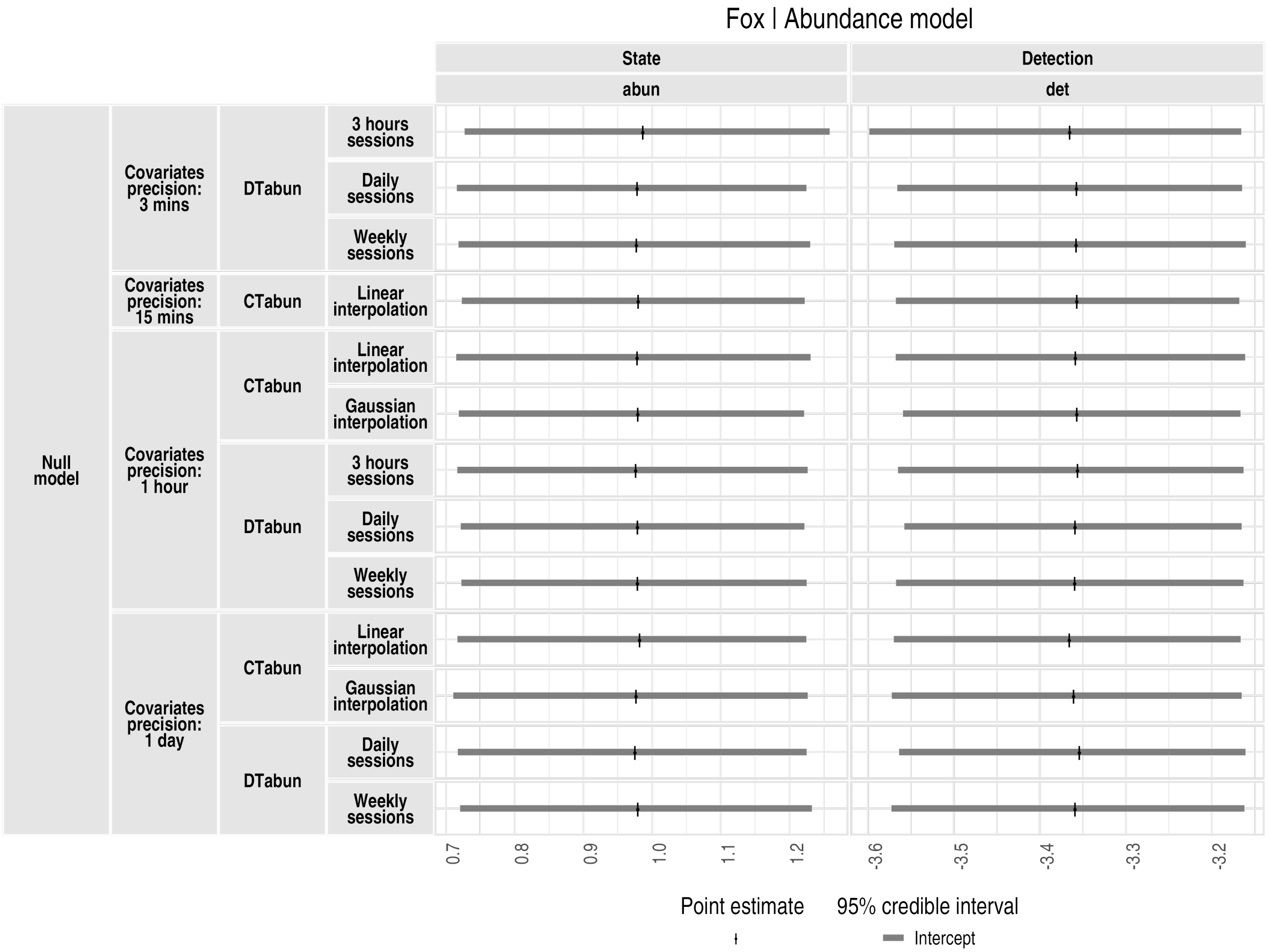

Figure S27: **Abundance model results for the fox (null model)**. `abun` refers to the abundance intercept (log scale) and `det` to the detection rate intercept (log scale). Grey horizontal lines show 95% credible intervals (posterior distribution quantiles), and black vertical lines indicate point estimates (posterior distribution medians). Model types and configurations are listed on the left side of the plot.

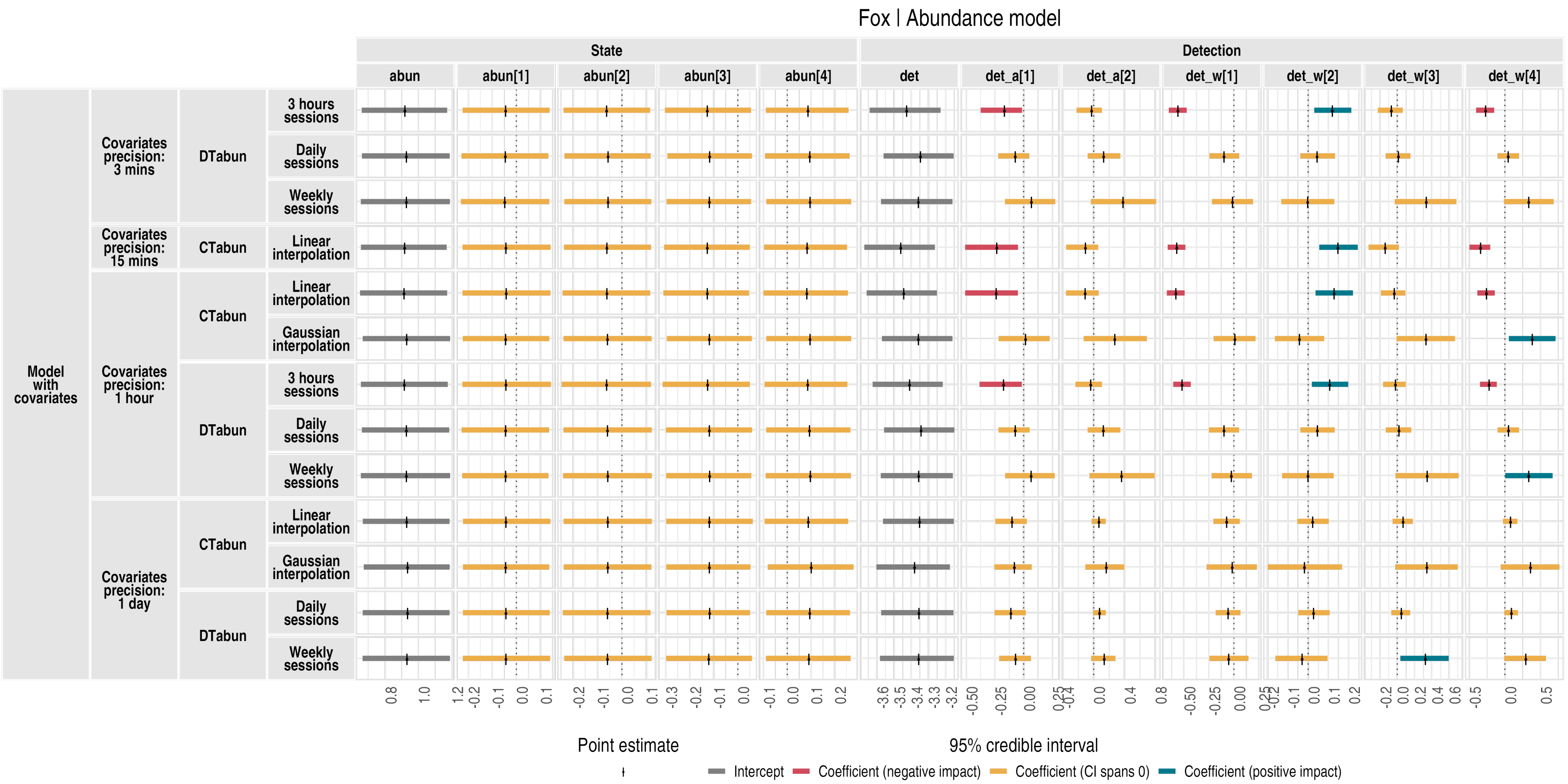

Figure S28: **Abundance model results for the fox.** `abun` is the abundance intercept, and `abun[1]` to `abun[4]` are the coefficients for habitat covariates (named `state_h[1]` to `state_h[4]` in the covariate plots). `det` is the detection rate intercept; `det_a[1]` and `det_a[2]` are the anthropogenic (human disturbance) detection covariate coefficients; `det_w[1]` to `det_w[4]` are the weather detection covariate coefficients. Grey horizontal lines show 95% credible intervals (posterior distribution quantiles), and black vertical lines indicate point estimates (posterior distribution medians). Model types and configurations are listed on the left side of the plot.

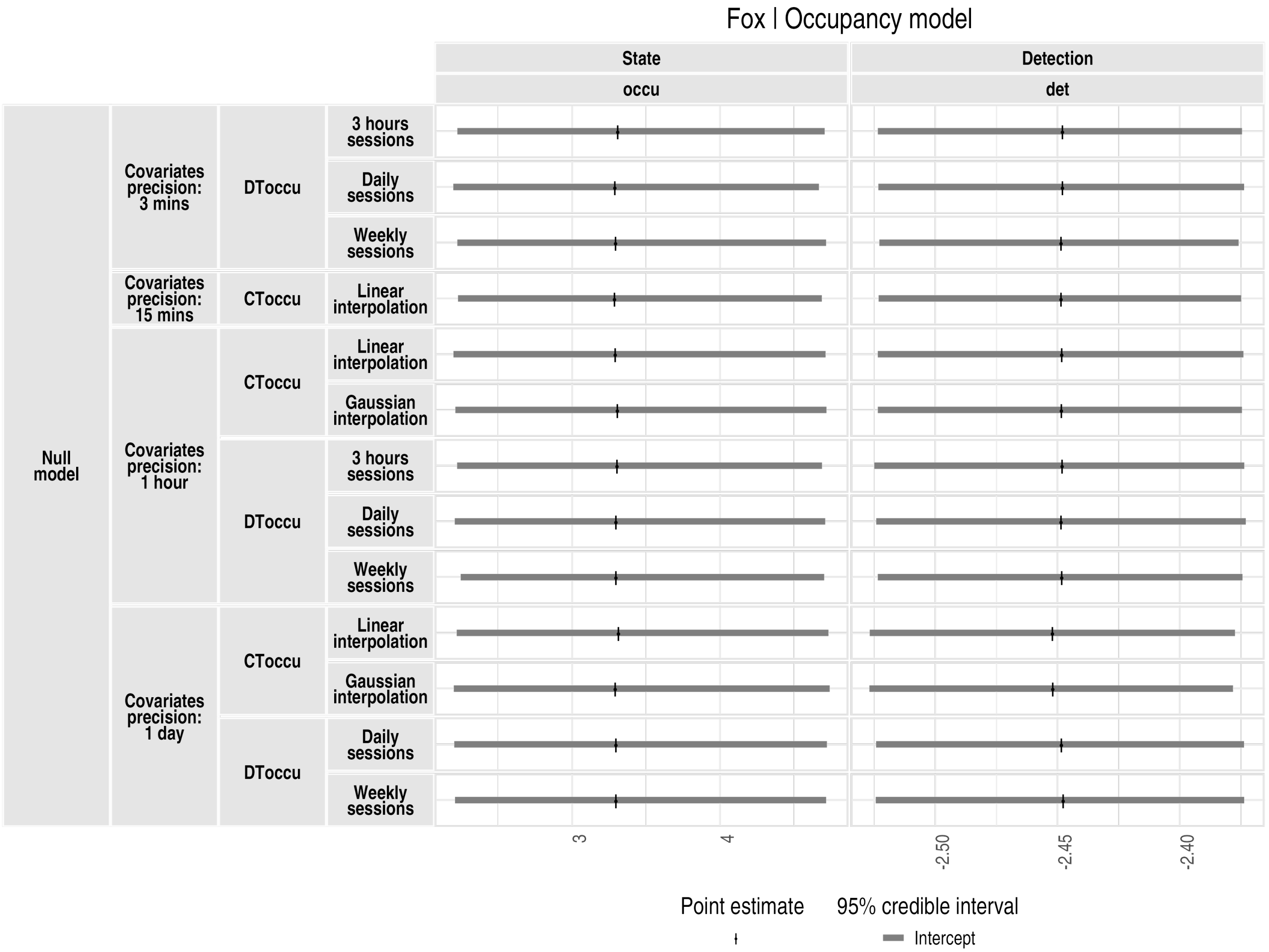

Figure S29: **Occupancy model results for the fox (null model)**. `occu` refers to the occupancy intercept (logit scale) and `det` to the detection rate intercept (log scale). Grey horizontal lines show 95% credible intervals (posterior distribution quantiles), and black vertical lines indicate point estimates (posterior distribution medians). Model types and configurations are listed on the left side of the plot.

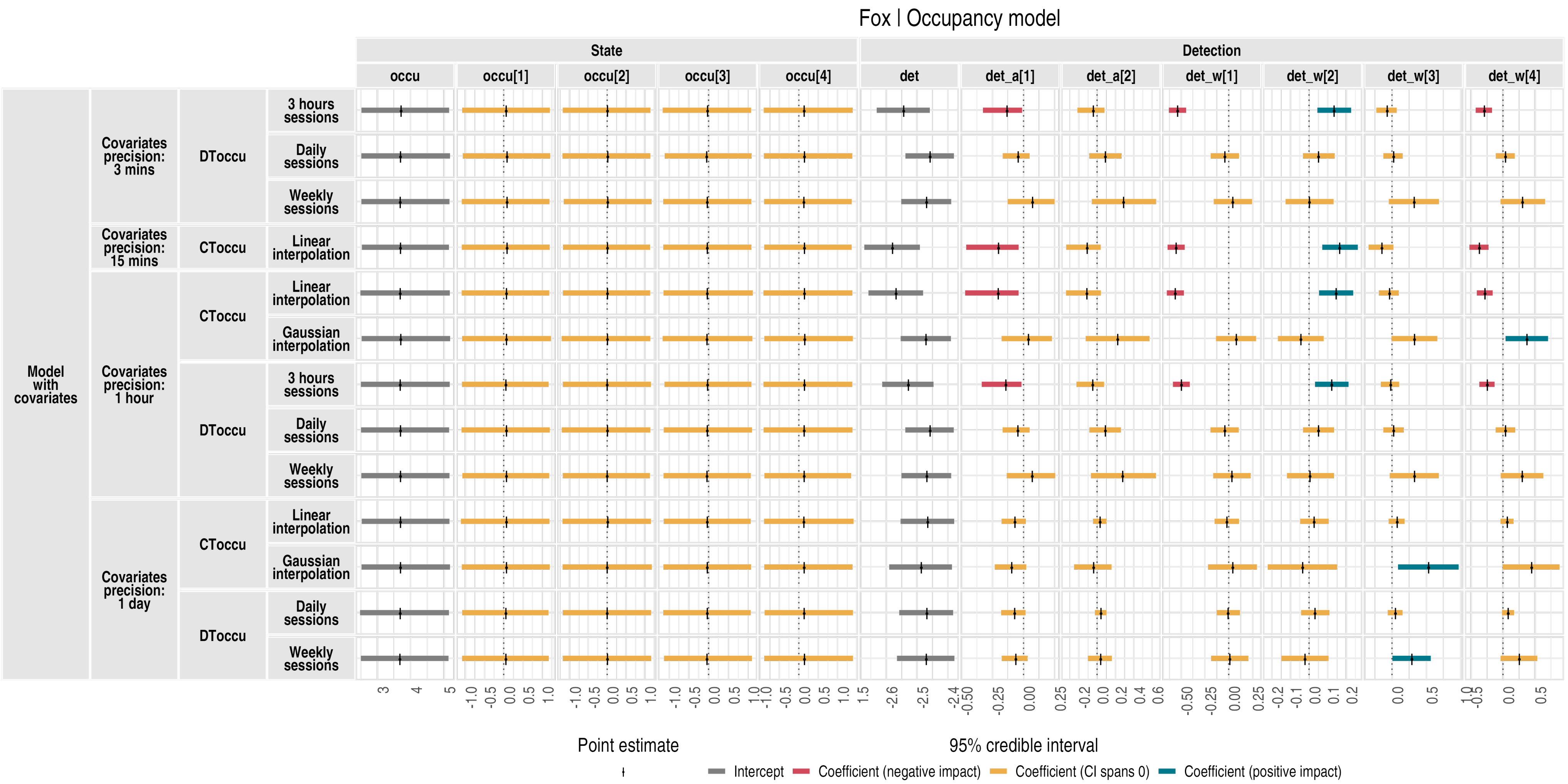

Figure S30: **Occupancy model results for the fox**. `occu` is the occupancy intercept, and `occu[1]` to `occu[4]` are the coefficients for habitat covariates (named `state_h[1]` to `state_h[4]` in the covariate plots). `det` is the detection rate intercept; `det_a[1]` and `det_a[2]` are the anthropogenic (human disturbance) detection covariate coefficients; `det_w[1]` to `det_w[4]` are the weather detection covariate coefficients. Grey horizontal lines show 95% credible intervals (posterior distribution quantiles), and black vertical lines indicate point estimates (posterior distribution medians). Model types and configurations are listed on the left side of the plot.

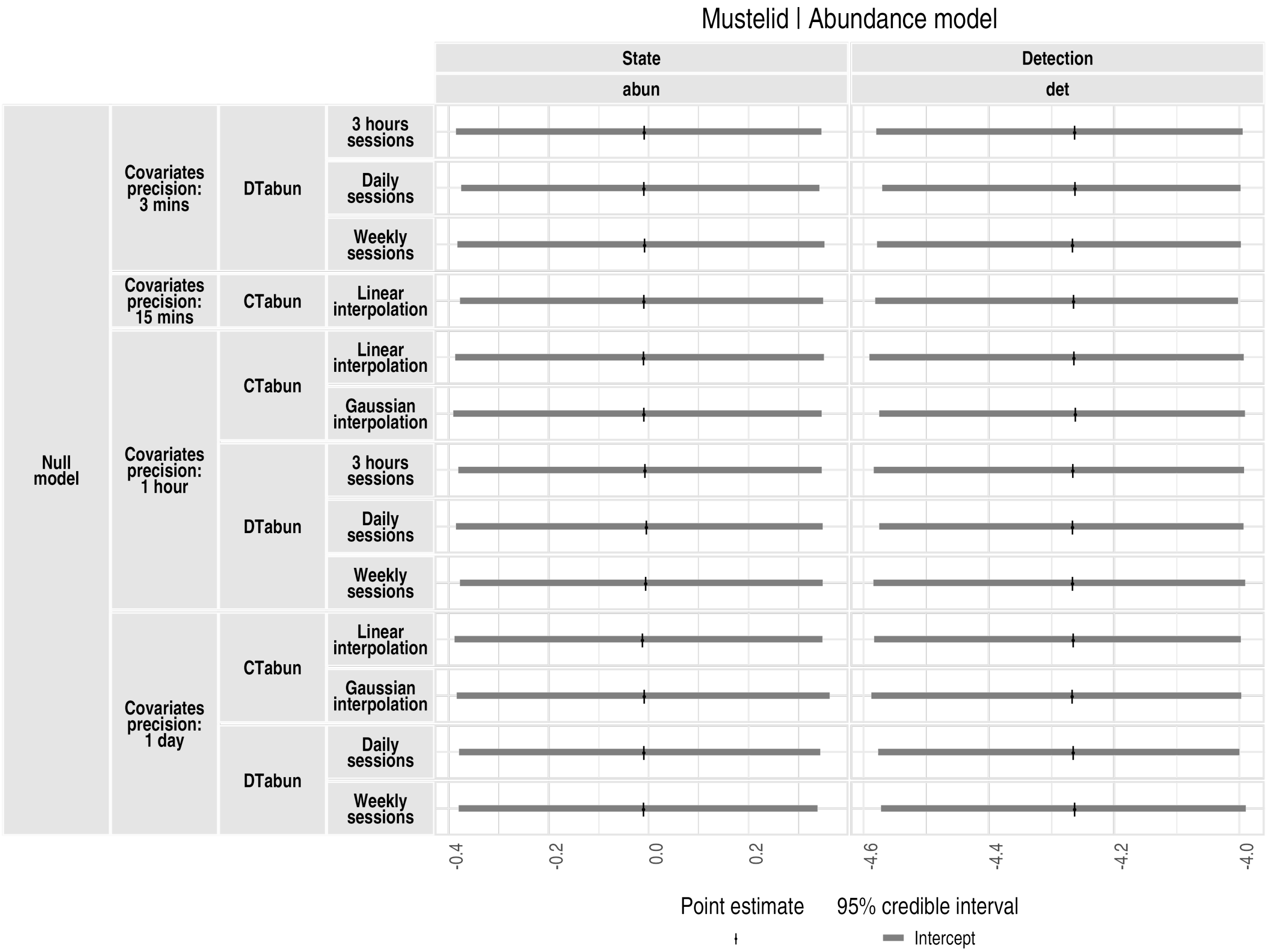

Figure S31: **Abundance model results for the small mustelids (null model)**. `abun` refers to the abundance intercept (log scale) and `det` to the detection rate intercept (log scale). Grey horizontal lines show 95% credible intervals (posterior distribution quantiles), and black vertical lines indicate point estimates (posterior distribution medians). Model types and configurations are listed on the left side of the plot.

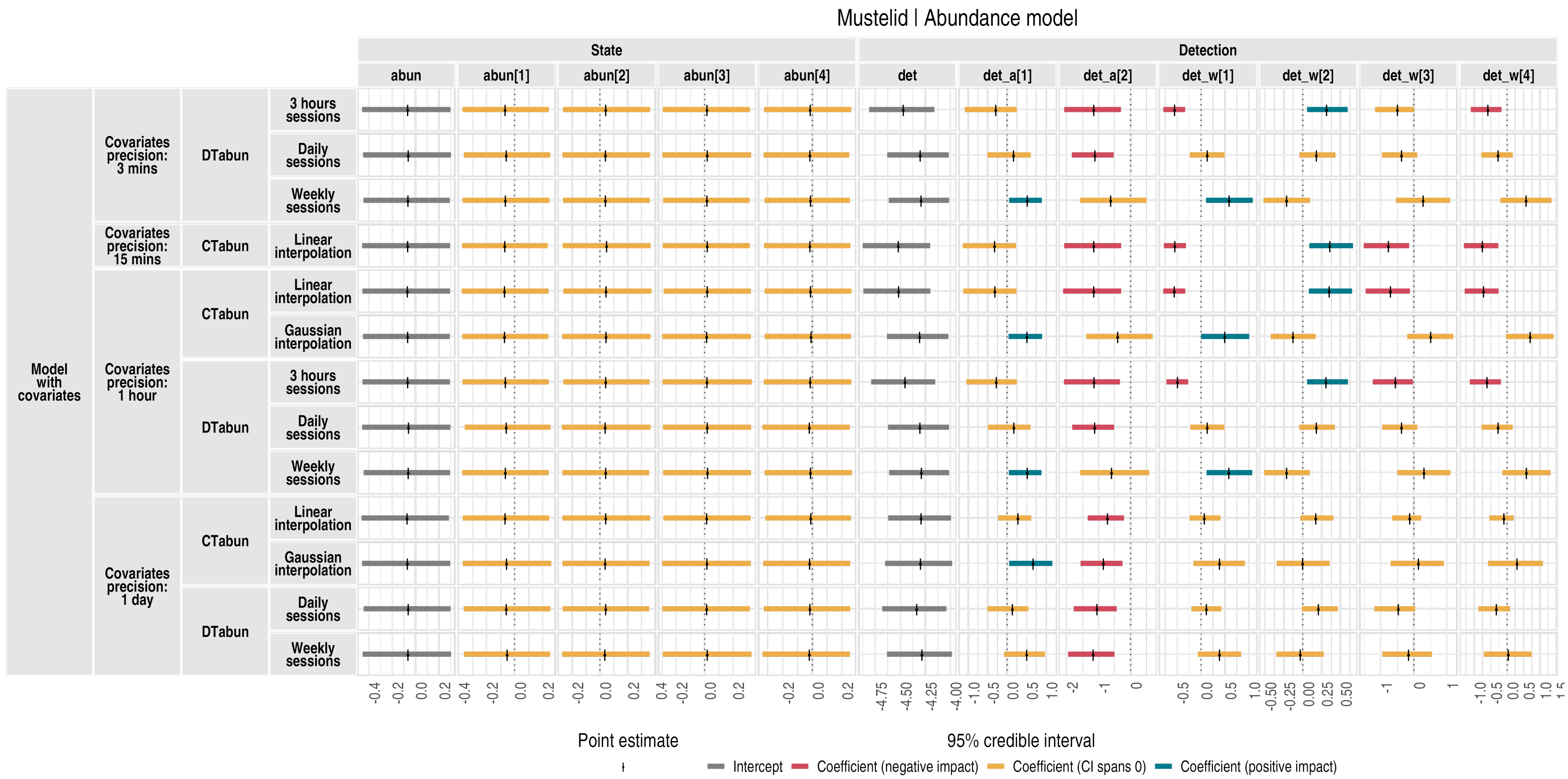

Figure S32: **Abundance model results for the small mustelids**. `abun` is the abundance intercept, and `abun[1]` to `abun[4]` are the coefficients for habitat covariates (named `state_h[1]` to `state_h[4]` in the covariate plots). `det` is the detection rate intercept; `det_a[1]` and `det_a[2]` are the anthropogenic (human disturbance) detection covariate coefficients; `det_w[1]` to `det_w[4]` are the weather detection covariate coefficients. Grey horizontal lines show 95% credible intervals (posterior distribution quantiles), and black vertical lines indicate point estimates (posterior distribution medians). Model types and configurations are listed on the left side of the plot.

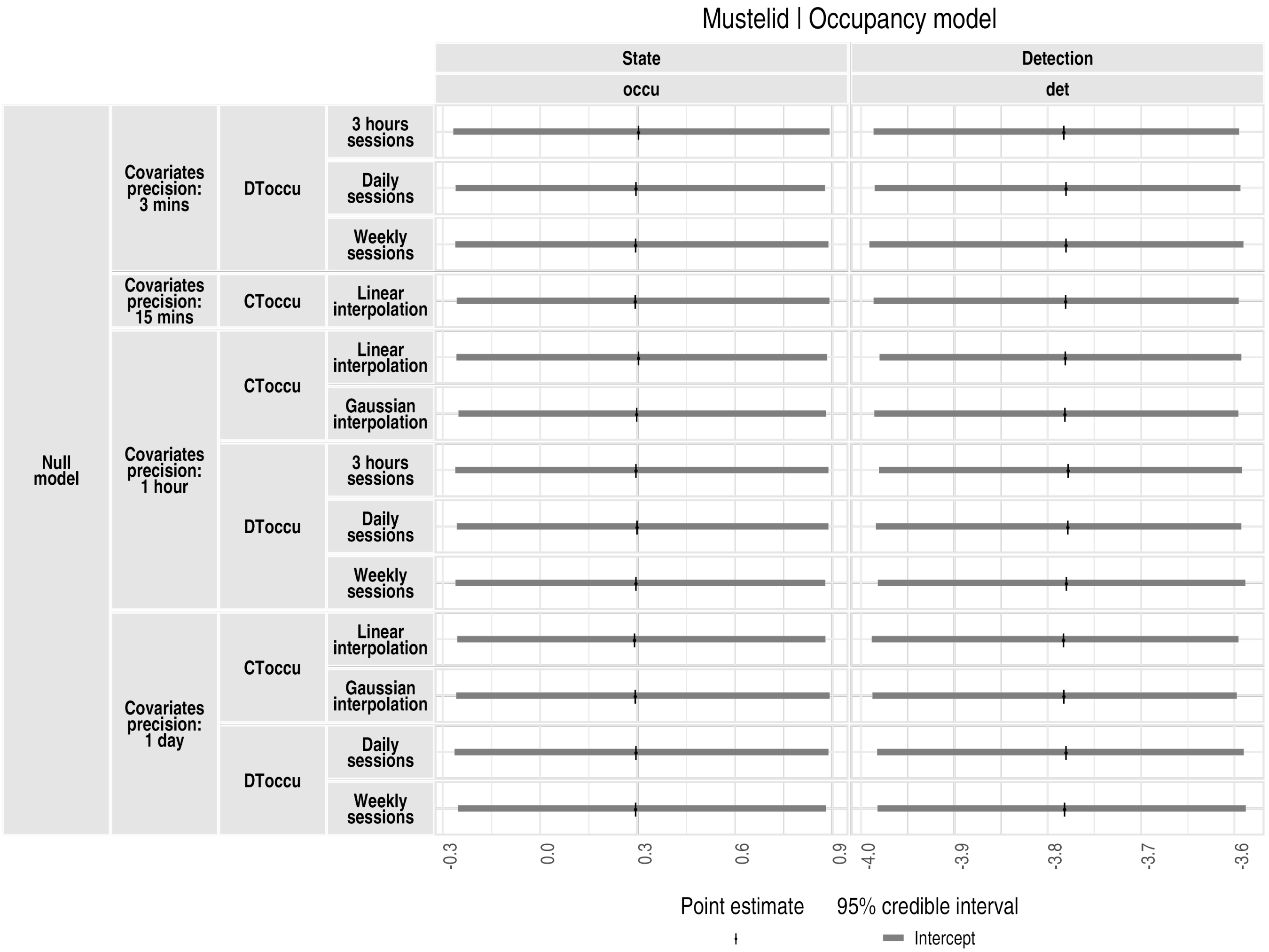

Figure S33: **Occupancy model results for the small mustelids (null model)**. `occu` refers to the occupancy intercept (logit scale) and `det` to the detection rate intercept (log scale). Grey horizontal lines show 95% credible intervals (posterior distribution quantiles), and black vertical lines indicate point estimates (posterior distribution medians). Model types and configurations are listed on the left side of the plot.

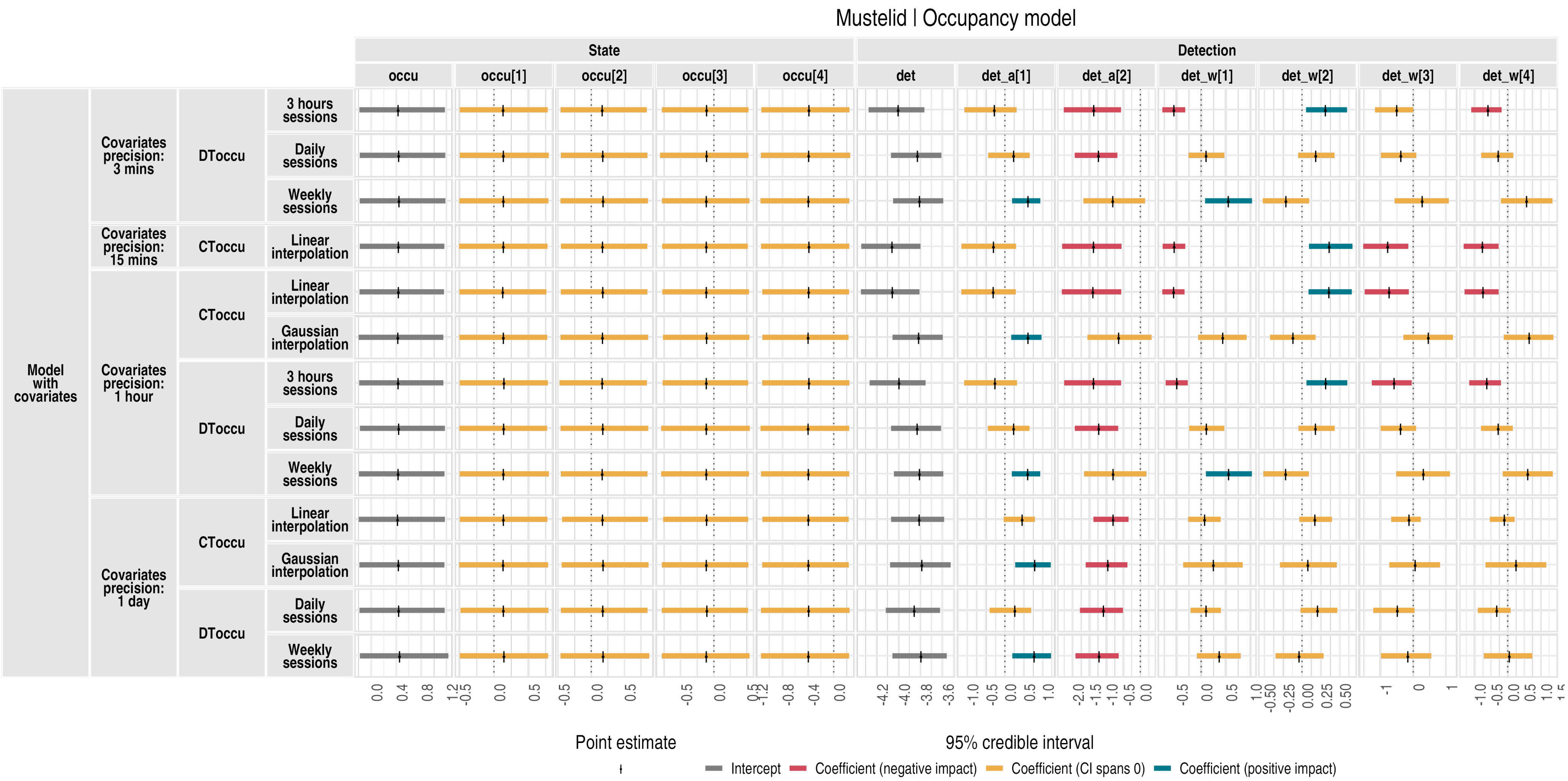

Figure S34: **Occupancy model results for the small mustelids**. `occu` is the occupancy intercept, and `occu[1]` to `occu[4]` are the coefficients for habitat covariates (named `state_h[1]` to `state_h[4]` in the covariate plots). `det` is the detection rate intercept; `det_a[1]` and `det_a[2]` are the anthropogenic (human disturbance) detection covariate coefficients; `det_w[1]` to `det_w[4]` are the weather detection covariate coefficients. Grey horizontal lines show 95% credible intervals (posterior distribution quantiles), and black vertical lines indicate point estimates (posterior distribution medians). Model types and configurations are listed on the left side of the plot.
